# Analysis of Rare Coding Variation Identifies New Genetic Contributors to Schizophrenia

**DOI:** 10.64898/2026.09.18.752670

**Authors:** Julia M. Sealock, Connor Dowd, Calwing Liao, Robert Ye, Daniel Howrigan, F. Kyle Satterstrom, Chiara Auwerx, Soyeon Kim, Cong Huai, Lin He, Melkam Alemayehu, Stella Gichuru, Rehema M. Mwende, Charles R.J.C. Newton, Nastassja Koen, Zukiswa Zingela, Joseph Kyebuzibwa, Anne Stevenson, Arsalan Hassan, Kai Wang, Makoto Arai, Yasue Horiuchi, Masanari Itokawa, Nora Ayola Serrano, Carrie E. Bearden, Sintia I. Belangero, Saulo G. Castor Albuquerque, Nicolas Crossley, Andre L. de Souza Rodrigues, Ana M. Diaz-Zuluaga, Mateus M. Diniz, Thiago H. Freitas, Ary Gadelha, Juliana Gomez-Makhinson, Nayana Holanda, Alex Kopelowicz, Pedro G. Lorencetti, Lucas C. Quarantini, Ana M. Ramirez-Diaz, Victor I. Reus, Marcos L. Santoro, Terri Teshiba, Carolina Ziebold, Carlo Esteban Sotelo-Ramirez, Marco Antonio Sanabrais-Jiménez, Eric Hahn, Van Phi Nguyen, Penelope A. Lind, Jennifer Forsyth, Adeniran Okewole, Rodney C.P. Go, Raquel Gur, Ruben Gur, Nicholas Craddock, Aarno Palotie, Eija Hämäläinen, Olli Pietiläinen, Stephen J. Glatt, Christina Hultman, Celso Arango López, Andrew McQuillin, Nick Bass, Ann E. Pulver, David St. Clair, Bruce Cohen, Douglas H. Blackwood, Andrew McIntosh, Tõnu Esko, Elizabeth Karlson, Aiden P. Corvin, Derek W. Morris, Dara Manoach, Mikael Landen, SCHEMA Consortium, Michael Boehnke, Anders D. Børglum, Claire Churchhouse, David Curtis, Michael C. O’Donovan, Michael J. Owen, Elliott Rees, Patrick F. Sullivan, Marquis P. Vawter, James T.R. Walters, Laura Scott, Sarah E. Medland, Van Tuan Nguyen, Thi Minh Tam Ta, Beatriz Camarena, Nelson Freimer, Carlos Lopez-Jaramillo, Loes Olde Loohuis, Roel A. Ophoff, Akira Sawa, Carlos N. Pato, Michele T. Pato, Muhammad Ayub, James A. Knowles, Rocky Stroud, Lukoye Atwoli, Akena Dickens, Symon M. Kariuki, Karestan C. Koenen, Dan J. Stein, Solomon Teferra, Shengying Qin, Mark J. Daly, Hailiang Huang, Benjamin M. Neale

## Abstract

Schizophrenia is a severe psychiatric disorder, affecting nearly 27 million individuals worldwide and imposing substantial personal and societal burden, yet its biological mechanisms remain incompletely understood. Rare coding variant association studies are a powerful approach for identifying genes contributing to schizophrenia risk, but detecting such associations requires large sample sizes given the polygenic nature of schizophrenia. In this study, we present the largest rare coding variant association study of schizophrenia to date, with 87,959 schizophrenia cases and 150,587 controls. We identify 16 genes at exome-wide significance: *SETD1A, ZMYM2, HERC1, RB1CC1, SCAF1, XPO7, SP4, FYN, PPP3CA, CUL1, HDAC9, JARID2, ATP9A, PTK2, STAG1,* and *SCN2A,* and an additional 24 at a 5% false discovery rate. Cases carrying ultra-rare, damaging variants, primarily protein-truncating and deleterious missense mutations, show strong enrichment in constrained genes with consistent effects across ancestries. Half of the 40 identified genes overlap with those implicated in developmental delay, autism, or bipolar disorder, underscoring shared neurodevelopmental risk. All 40 genes identified are highly expressed in excitatory and inhibitory neurons, and their expression patterns span diverse developmental stages. Functionally, the identified genes implicate chromatin regulation, protein degradation, and synaptic function. Overall, our results advance understanding of the genetic architecture of schizophrenia, provide a foundation for modeling risk genes in cellular and animal systems, and offer a framework for future work toward biologically informed patient stratification.

## Introduction

Schizophrenia is a complex and severe psychiatric disorder characterized by heterogeneous combinations of symptoms, including hallucinations, delusions, blunted affect, and avolition^1^. Typically diagnosed in late adolescence or early adulthood, schizophrenia is associated with significant long-term personal and societal costs, including decreased life expectancy of approximately 15 years and a higher lifetime risk of death by suicide^2,3^. Our limited understanding of the biological mechanisms of schizophrenia contributes to inadequate treatments and prevention strategies.

Genetics can offer insights into the foundational biological changes that lead to schizophrenia. The most recent genome-wide association study (GWAS) of schizophrenia identified 354 distinct common variant loci^4^; however, these variants are primarily located in non-coding regions with small effect sizes, making biological interpretation challenging. Despite the success of GWAS in schizophrenia, its full genetic architecture remains undiscovered^5^.

Rare variant association studies (RVAS) of coding variants are a powerful way to identify genes associated with schizophrenia^6^. Rare variants often have larger effect sizes because deleterious variants with large effects are subject to strong negative selection, which keeps their population frequency low^7^. By focusing on damaging coding (i.e., gene-altering) variants, RVAS directly identifies genes where damaging changes have an impact on disease risk.

Identifying damaging variants via annotations is key to RVAS. Variants predicted to be damaging are classified as protein-truncating variants (PTVs) or highly damaging missense variants by algorithms such as LOFTEE for PTVs, and MPC, AlphaMissense, or MisFit for missense variants^8–11^. RVAS typically aggregate predicted damaging variants within genes to increase power to detect gene-level associations with disease, particularly when individual variants are too rare to test independently. The genes identified by RVAS provide strong evidence for follow-up functional analysis to uncover disease mechanisms.

The Schizophrenia Exome Meta-Analysis (SCHEMA) Consortium is a global collaboration dedicated to identifying genes associated with schizophrenia by studying ultra-rare, gene-disrupting mutations. The initial SCHEMA analysis identified 10 genes significantly associated with schizophrenia^6^. Follow-up work in the Psychiatric Genomics Consortium Phase 3 Targeted Sequencing of Schizophrenia Study (PGC3SEQ) identified three additional genes,^12^ and a subsequent reanalysis of SCHEMA 1.0 with additional cases yielded two more genes^13^. Building on the successes of RVAS for gene discovery for schizophrenia, we present results for our second analysis wave, SCHEMA 2.0, based on updated methods and a greatly expanded sample size of 87,959 cases and 150,587 controls.

## Results

### Identifying ultra-rare, damaging variants

We aggregated new exome sequencing data with existing data from SCHEMA 1.0 and additional schizophrenia cases and controls included in gnomAD v4. In total, the SCHEMA 2.0 dataset includes sequencing data from 103,441 schizophrenia cases and 189,856 controls generated using five exome capture methods across 120 cohorts, including 19,919 cases and 35,519 controls from SCHEMA 1.0.

We performed extensive filtering to ensure quality of samples, variants, and genotypes, retaining 87,959 schizophrenia cases and 150,587 controls for case-control analysis, including 58,151 cases new to this report (Supplementary Methods, Supplementary Tables 1-3). We classified damaging variants as either high-confidence protein-truncating variants (PTVs) or deleterious missense variants. We defined deleterious missense variants using MisMeanRank, a composite rank-based score across multiple pathogenicity metrics (Methods).

Variants with large deleterious effects, particularly those in developmentally important genes or those whose associated phenotypes emerge before or during reproductive years, tend to be selected out of the population due to natural selection, leading to extremely low frequencies^7^. We extracted PTVs and damaging missense variants with minor allele count (MAC) ≤ 15 and examined their frequencies. Most damaging variants in our dataset are singletons, confirming that deleterious variants are maintained at extremely low frequencies in the population (Supplementary Figure 11). As a result, ultra-rare variants (MAC ≤ 15) are predominantly ancestry-specific. However, cross-ancestry sharing increases with allele count, with 38% of non-singleton variants present in multiple populations, increasing to 48% at MAC ≥ 3 and 63% at MAC ≥ 10. This pattern is consistent with recurrent recent mutational events across populations, rather than ancestry-specific variation typical of common variants (Supplementary Figure 12).

To rule out the possibility that variants that are uncommon but not ultra-rare could appear ultra-rare due to smaller ancestry sample sizes, we conducted sensitivity analyses excluding variants with a population-maximum (popmax) allele frequency >0.1% in external reference datasets (gnomAD v4 and the Regeneron Million Exome Cohort). Among ultra-rare PTVs in SCHEMA 2.0, only 555 of 550,916 variants (0.10%) exceeded this threshold in gnomAD, and 991 variants (0.17%) exceeded it in the Regeneron dataset. Furthermore, when calculating population-specific allele frequencies within the SCHEMA 2.0 dataset, only 205 PTVs observed only in the Ashkenazi Jewish population exceed a frequency of 0.1%, consistent with the well-documented bottlenecked population history of this group (Supplementary Figure 13). These results indicate that variants that are higher-frequency in sub-populations are infrequently misclassified as ultra-rare in our dataset.

Using ultra-rare variants, we assessed the enrichment of rare, damaging variants in our dataset across the ultra-rare MAC bins and ancestry. Constrained genes are those under strong purifying selection, evidenced by a depletion of deleterious variation relative to expectation in population sequencing data. Because functional disruption of constrained genes is more likely to harm the organism, we restricted enrichment analyses to genes with predicted loss-of-function intolerance (pLI ≥ 0.9). Across all samples, damaging singletons in constrained genes showed the strongest enrichment in schizophrenia cases versus controls, with PTVs exhibiting the strongest enrichment (PTV: OR = 1.21, p-value = 7.92 x 10^-92^; PTV + missense: OR = 1.19, p-value = 5.12 x 10^-154^, Missense: OR = 1.17, p-value = 1.75 x 10^-67^; Synonymous: OR = 1.00, p-value = 0.982). PTVs with 2 ≤ MAC ≤ 5 showed significant but reduced enrichment in cases (OR = 1.08, p-value = 1.67 x 10^-4^). The effect of variants with MAC 6 ≤ MAC ≤ 10 and MAC 11 ≤ MAC ≤ 15 indicated no significant enrichment in cases over controls, however, these analyses of strata are limited in power due to the small number of variants compared to the lower MAC groups (Supplementary Figure 14A, Supplementary Table 4).

When stratified by genetic ancestry groupings, all ancestries demonstrated significant enrichment of PTV singletons in schizophrenia cases versus controls. These findings support a model in which the most damaging mutations are of very recent origin with independent mutational events across populations (Supplementary Figure 14B, Supplementary Table 5).

### Identifying genes implicated by RVAS

After confirming the enrichment of rare, damaging mutations across annotations in schizophrenia cases versus controls, we sought to identify genes implicated in schizophrenia using a meta-analysis approach. In case-control RVAS, we restricted to ultra-rare variants with a MAC ≤ 15 across the entire SCHEMA 2.0 dataset. We performed an RVAS for PTV variants only and for PTV and damaging missense variants for all protein-coding genes in the genome. For each gene, we selected the annotation with the most significant p-value.

We identified 16 genes with exome-wide Bonferroni significance and an additional 24 genes with FDR 5% significance (Supplementary Table 6). Six of the 10 exome-wide significant genes from SCHEMA 1.0 remained significant in SCHEMA 2.0: *SETD1A, HERC1, RB1CCI, SP4, XPO7,* and *CUL1.* New exome-wide significant associations include *SCAF1, FYN, PPP3CA, HDAC9, JARID2, ATP9A, PTK2,* and *SCN2A* (Figure 1A and Figure 1B). Notably, we report an FDR 5% association with *CHRM4* (cholinergic receptor muscarinic 4), one of the targets of the recently approved antipsychotic xanomeline/trospium^14^.

**Figure 1.**
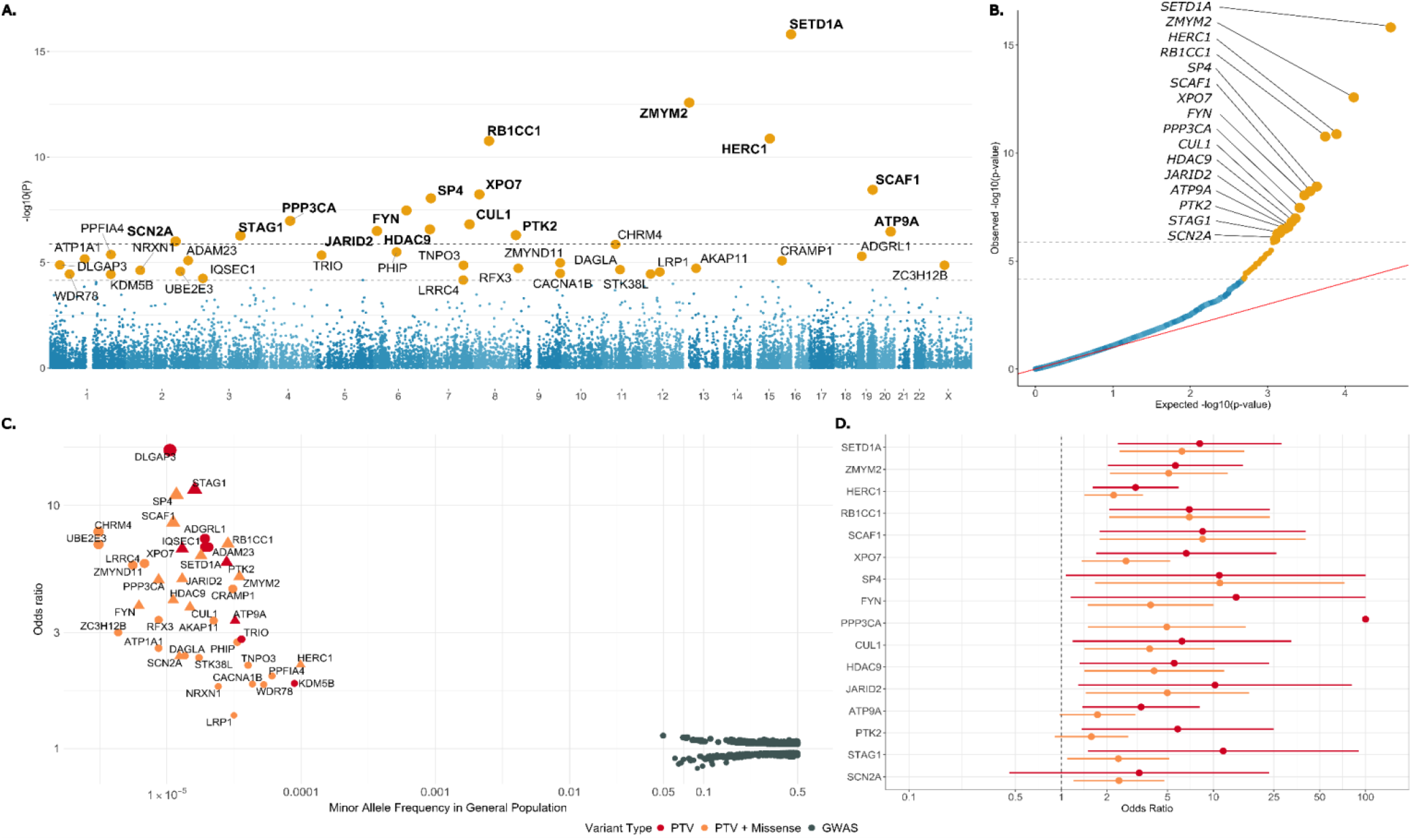
Results from analysis of ultra rare variants in SCHEMA 2.0. **A.** Manhattan plot with -log_10_-transformed p-values plotted against chromosomal location of each gene. The per-gene p-values were calculated by selecting the most significant case-control p-value between PTV only CMH tests and PTV + damaging missense CMH tests cases and controls. Genes in bold represent exome-wide significant associations. The dark gray line indicates Bonferroni significance (p < 1.31 x 10^-6^) and the light gray dashed line indicates FDR 5% significance (p < 6.84 x 10^-5^). **B.** Quantile-quantile plot of observed -log_10_ p-values plotted against the expectation given a uniform distribution. The dotted lines indicate FDR 5% significance and Bonferroni significance. Labelled genes represent exome-wide significant associations. **C.** Genetic architecture of schizophrenia risk plotted as odds ratios against the minor allele frequency in the general population for each type of genetic variants, with PTVs in red, PTV + MisMeanRank 93% in orange, and GWAS common variants in gray. Genes reaching exome-wide significance are denoted as triangles. The frequency in the general population for PTVs was extracted from gnomAD v4.0. The frequency in the general population for common variants was extracted from GWAS MAF estimates. The size of each dot is proportional to the odds ratio. OR estimates with no control carriers were capped at an OR of 100. **D.** Forest plot of log-transformed PTV odds ratios and PTV + MisMeanRank 93% for SCHEMA 2.0 exome-wide significance. OR and upper 95% confidence interval estimates were capped at OR = 100. Confidence intervals were estimated using the Robins-Breslow-Greenland variance estimator.

Among the 40 FDR 5% genes, 35 demonstrated an odds ratio greater than 2, indicating that, on average, rare, damaging mutations in these genes are associated with a twofold increase in the predicted odds of schizophrenia (Figure 1C). Across all cases, only 1.5% carry a PTV in one or more of the 40 genes significant at FDR 5% compared to 0.5% of controls, for an overall odds ratio of 3.43 (95% CI = 3.12 - 3.76). Individual variant effects, however, may range from neutral to much more severe than this average.

We report all high-quality variants, annotations, carrier frequencies, and gene-level results in a web browser at https://schema.broadinstitute.org/.

Variant annotation plays a critical role in the interpretation of RVAS results. As expected, the most deleterious annotation, PTV, drives the association signal for many genes, however, statistical evidence often increases when damaging missense variants are included (Supplementary Figure 15). In fact, for 27 of the 40 FDR 5% genes, the PTV-only signal is weaker than the signal from the combined PTV and damaging missense burden, suggesting that both types of damaging variation in the same gene contribute to schizophrenia risk.

We conducted sensitivity analyses using trio data from SCHEMA 1.0. We updated the *de novo* mutations from 3,402 schizophrenia trios to match the genome build and variant annotations of the case-control data (Supplementary Methods). Using mutation rates from gnomAD v2, we calculated the Poisson p-value for PTV variants, and PTV plus damaging missense variants on 457 genes with at least one *de novo* variant. We combined the p-values of the case-control analysis with the *de novo* Poisson p-values using a z-score weighted meta-analysis. Overall, the inclusion of *de novo* data had minimal impact on the results, with only two notable changes. *TRIO* rose to exome-wide significance (p = 1.07 x 10^-6^), supported by two *de novo* variants, and *YLPM1* reached the FDR 5% significance threshold (p = 6.70 x 10^-5^) (Supplementary Figure 5)

### Comparing SCHEMA 2.0 to SCHEMA 1.0

SCHEMA 2.0 methods differ from SCHEMA 1.0 in key ways, including changes to the damaging missense definition from MPC only to a rank-based annotation, exclusion of ‘other missense’ mutations, and exclusion of missense-only RVAS testing in SCHEMA 2.0. Comparison of the top gene findings from SCHEMA 1.0 to SCHEMA 2.0 showed that 7 of the top 10 genes remained at or above FDR 5% significance, with 6 genes, *SETD1A, HERC1, RB1CC1, XPO7, SP4,* and *CUL1,* remaining exome-wide significant (Supplementary Figure 19).

In the published SCHEMA 1.0 results, restricting the analysis to Class 1 case–control variants (PTVs and damaging missense variants with MPC ≥ 3), without incorporating *de novo* mutations, would have yielded only four genes reaching exome-wide significance, *XPO7, SETD1A, HERC1,* and *TRIO*. In comparison, SCHEMA 2.0 case-control data identified 16 such genes, an increase driven by the larger case-control sample size. Several genes that required *de novo* support in SCHEMA 1.0 (such as *CUL1*, *RB1CC1*, and *SP4*) are now identified by case– control data alone, suggesting that *de novo* evidence in SCHEMA 1.0 helped prioritize true risk genes that are subsequently supported by larger case–control datasets, rather than introducing spurious associations. Incorporating *de novo* information into SCHEMA 2.0 does not substantially increase the number of significant genes. As noted above, *TRIO* is the only additional exome-wide significant gene when *de novo* data are included, indicating that the increased sample size in SCHEMA 2.0 accounts for the majority of the discovery power, with *de novo* data contributing comparatively modest additional signal relative to SCHEMA 1.0.

To evaluate the impact of missense annotation definitions between SCHEMA 1.0 and SCHEMA 2.0, we compared results using the different missense definitions. When repeating the SCHEMA 2.0 RVAS using the SCHEMA 1.0 definition of MPC > 3 in place of MisMeanRank ≥ 93 (methods), the number of exome-wide significant genes decreases to 13, and the number of genes reaching FDR 5% declines to 25 (Supplementary Figure 20). This reduction suggests that reliance on a single missense metric captures a narrower subset of deleterious variation and consequently reduces statistical power. In contrast, MisMeanRank integrates information across multiple missense annotations, enabling more comprehensive prioritization of damaging variants and improved sensitivity to gene-level burden.

We compared the odds ratios and confidence intervals for the PTV and PTV + MisMeanRank ≥ 93 analyses across exome-wide significant genes. Both models demonstrated 2-10 fold increased odds of schizophrenia per gene, consistent with strong effects for these ultra-rare and deleterious mutations. However, confidence intervals remain wide despite the large SCHEMA 2.0 sample size, reflecting the low carrier counts inherent in rare variant studies; with few carriers per gene, odds ratio estimates become unstable and confidence intervals widen dramatically. We next compared PTV odds ratios for the ten exome-wide significant genes from SCHEMA 1.0 against their corresponding estimates in SCHEMA 2.0 (Supplementary Figure 21). Odds ratios were broadly consistent across both versions, with all genes demonstrating odds ratios greater than 1 in both SCHEMA 1.0 and SCHEMA 2.0. Although the odds ratios are slightly attenuated in SCHEMA 2.0 compared to SCHEMA 1.0, the wide confidence intervals overlap across both versions.

Notably, three genes that reached exome-wide significance in SCHEMA 1.0 did not meet the FDR 5% threshold in SCHEMA 2.0, *GRIN2A, GRIA3,* and *CACNA1G.* This change may be driven by more stringent sample and genotype quality control in the updated analysis, but the inclusion of newly ascertained cohorts likely also plays a role. All three genes are established risk factors for developmental delay/intellectual disability (DD/ID). When examining pLoF carrier frequencies in DD/ID genes across SCHEMA 2.0 samples, we observed higher case frequencies in cohorts that were part of SCHEMA 1.0 compared to newly added cohorts (Supplementary Figure 22, Supplementary Table 7). This suggests that newer cohorts may be milder schizophrenia cases or less enriched for schizophrenia cases with comorbid DD/ID, thereby diluting the signal for genes with cross-disorder effects. Additionally, a population-based analysis of the UK Biobank showed a higher frequency of PTVs in *CACNA1G* than reported in SCHEMA 1.0 cases, indicating a potentially lower impact on schizophrenia risk than previously reported^15^. Importantly, despite not reaching statistical significance in SCHEMA 2.0, *GRIA3* and *GRIN2A* still indicate substantial case enrichment: *GRIA3* has 6 PTV case carriers and 1 PTV control carrier (OR = 5.80), while *GRIN2A* has 10 case carriers and 5 control carriers (OR = 2.74), supporting persistent, though attenuated, enrichment of damaging mutations in schizophrenia cases.

### Overlap Across Neuropsychiatric Genes

We tested for schizophrenia case enrichment of damaging rare variants (MAC <= 15) in genes implicated in other neurodevelopment disorders, developmental delay, intellectual disability (DD/ID), and autism spectrum disorder (ASD). DD/ID has 309 genes implicated^16^, while the most recent ASD rare variant analysis implicated 253 genes^17^. Within DD/ID associated genes, schizophrenia cases were significantly enriched for PTVs (OR = 1.27, p-value = 1.23 x 10^-26^), PTV + damaging missense variants (OR = 1.21, p-value = 1.53 x 10^-53^), and damaging missense only variants (OR = 1.16, p-value = 3.74 x 10^-30^). Singletons within ASD-associated genes demonstrated slightly higher enrichment, with the strongest association in PTV only variants (OR = 1.47, p-value = 4.10 x 10^-46^), followed by the combined signal of PTV and damaging missense variants (OR = 1.27, p-value = 4.05 x 10^-60^), and damaging missense only variants (OR = 1.19, p-value = 1.31 x 10^-24^ (Figure 2A, Supplementary Table 8).

**Figure 2.**
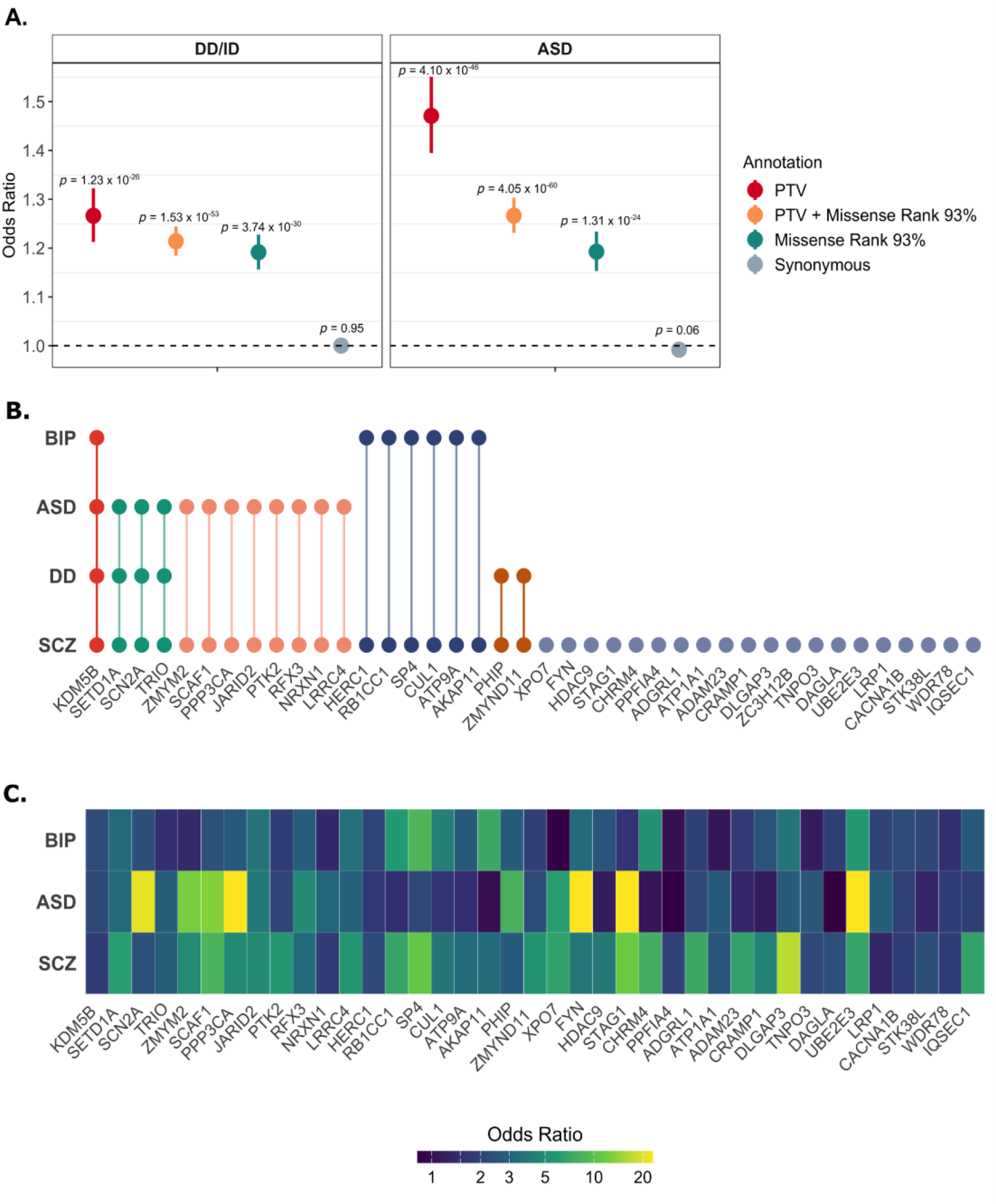
Shared genetic architecture between schizophrenia and other psychiatric disorders. **A.** Case-control enrichment of damaging singleton variants in genes associated with DD/ID (309 genes) or ASD (genes) stratified by annotation. DD/ID and ASD genes were identified from the most recent exome sequencing studies (Kaplanis et al, Fu et al). Logistic regressions were conducted within each ancestry separately, controlling for sex, capture, and top five principal components. The odds ratios were meta-analyzed using a fixed-effect inverse weighting meta-analysis for each annotation with the meta-analysis p-values reported above each annotation. The dots represent the meta-analyzed odds ratios, and the bars represent the 95% confidence intervals of the point estimates. **B.** Overlap of genes identified in schizophrenia (SCZ) with genes identified in developmental delay/intellectual disability (DD), autism spectrum disorders (ASD), and bipolar disorder (BIP). Dots represent whether the gene met an FDR 5% significance within each trait, and colors represent groupings based on overlap across the same traits. **C.** Heatmap of odds ratios (ORs) for genes identified in schizophrenia (SCZ) across autism spectrum disorders (ASD) and bipolar disorder (BIP). Genes are ordered based on the overlap identified in Figure 2B. Genes with infinite odds ratios were capped at 20.

Among the 40 FDR 5% significant genes, half overlap with DD/ID, ASD, or bipolar disorder (BD), indicating substantial shared genetic architecture between neurodevelopmental and psychiatric disorders. Notably, *KDM5B* is associated with all four disorders. Among the shared genes, three are associated with schizophrenia, DD/ID, and ASD, *SETD1A*, *SCN2A,* and *TRIO*. In the newest RVAS of bipolar disorder (BipEx 2.0), six genes overlap between schizophrenia and bipolar, *RB1CC1, HERC1, SP4, CUL1, ATP9A,* and *AKAP11* (Figure 2B).

Using the overlapping genes, we compared odds ratios (ORs) across schizophrenia, ASD, and bipolar disorder (Figure 2C). Because most of the DD/ID data were derived from trios, ORs could not be calculated for this trait. Across the three disorders, 12 genes exhibited ORs ≥ 2: *SETD1A, SCAF1, SP4, FYN, PPP3CA, JARID2, SCN2A, PHIP, DLGAP3, RFX3, UBE2E3,* and *LRC3.* Notably, one gene, *UBE2E3,* demonstrated an OR > 5 in all three traits. Four additional genes (*ZMYM2, SCAF1, XPO7,* and *STAG1*) had ORs ≥ 5 in schizophrenia and ASD, and three additional genes (*RB1CC1, SP4,* and *CHRM4*) showed ORs ≥ 5 in schizophrenia and bipolar disorder. These results highlight both shared biology across disorders through high-effect risk genes and schizophrenia-specific biology, where several high effect genes show limited evidence with other traits.

### Shared GWAS signal

Common and rare variants both play a role in increasing risk for schizophrenia, and we expect that GWAS and RVAS signals will converge on similar genetic regions. We used the latest GWAS of schizophrenia^18^ to examine the overlap between SCHEMA genes and GWAS loci. We selected the 287 independent GWAS loci (GRCh38 coordinates), and determined which index SNPs fell within ± 500 kb of the 40 SCHEMA 2.0 FDR 5% genes. We identified GWAS index SNPs near the following six SCHEMA 2.0 genes: *STAG1, FYN, SP4, CHRM4, LRP1,* and *SCAF1* (Table 1, Supplementary Figure 23A). The overlap between GWAS and RVAS signals at *CHRM4* provides convergent evidence across the common and rare allele frequency spectrum, further highlighting the value of integrating common and rare variant studies for target prioritization.

**Table 1.** Overlap between schizophrenia GWAS index SNPs and SCHEMA 2.0 genes. Index SNPs were deemed to be overlapping SCHEMA genes if they were within 500 kb of a SCHEMA 2.0 FDR 5% gene. GWAS coordinates were converted from GRCh37 to GRCh38.

| ***SCHEMA Gene*** | ***Locus (GRCh38)*** | ***Reference Allele*** | ***Alternate Allele*** | ***GWAS p-value*** | ***GWAS Beta (SE)*** |
|---|---|---|---|---|---|
| *STAG1* | chr3:136434626 | T | C | 7.93e-15 | 0.06 (0.008) |
| *FYN* | chr6:111501486 | A | G | 4.40e-8 | 0.06 (0.01) |
| *SP4* | chr7:21434992 | C | G | 1.22e-9 | 0.05 (0.009) |
| *CHRM4* | chr11:46502463 | T | C | 2.69e-18 | -0.09 (0.01) |
| *LRP1* | chr12:57289173 | C | T | 1.53e-14 | 0.12 (0.02) |
| *SCAF1* | chr19:49659652 | C | G | 3.79e-13 | 0.06 (0.008) |

Although *FURIN* did not meet the SCHEMA 2.0 FDR threshold, it represents a compelling candidate for convergence. Previous GWAS eQTL mapping implicated *FURIN* as the sole gene at its associated locus, with the schizophrenia risk allele associated with reduced expression^19^. Consistent with this biology, rare protein-truncating and damaging missense variants in *FURIN* showed a substantial effect size in SCHEMA 2.0 (OR = 4.39, *P* = 1.46e-4), suggesting that larger rare variant studies may identify *FURIN* as an exome-wide significant schizophrenia risk gene.

We used permutation to determine whether this observed overlap exceeded chance expectations. In each of 100,000 permutations, we randomly sampled 40 genes from the protein-coding genome and counted the number of genes located near a GWAS SNP (Supplementary Figure 23B). An empirical p-value was calculated as the proportion of permutations in which the number of overlapping genes was greater than or equal to the observed overlap of six genes. Under this null model, the observed overlap was not greater than expected by chance (empirical p = 0.41), indicating that SCHEMA FDR 5% genes do not show significant enrichment near GWAS index loci.

Although genes identified by RVAS were not enriched near GWAS index loci, schizophrenia cases in SCHEMA 2.0 showed significant enrichment of damaging URVs within fine-mapped GWAS genes. Among the 64 fine-mapped protein coding genes^18^, the strongest enrichment was observed with damaging missense variants (OR = 1.28, p-value = 6.45e-6), followed by the combined signal of PTVs and damaging missense variants (p-value = 5.74e-4, OR = 1.11) (Supplementary Table 9, Supplementary Figure 23C). These results indicate that despite limited gene-level overlap between RVAS genes and GWAS loci, schizophrenia cases are enriched for damaging URVs within GWAS-implicated genes, highlighting convergence across complementary genetic approaches.

### Gene Expression Patterns

Next, we sought to understand where and when SCHEMA genes are expressed in cells and tissues. We first examined tissue-specific expression patterns of the 40 FDR 5% genes using bulk RNA-sequencing data from GTEx in FUMA^20^ (Figure 3A, Supplementary Table 10). Seven brain regions showed significant overexpression, the frontal cortex (BA9), cerebellum, cerebellar hemisphere, general cortex, nucleus accumbens basal ganglia, anterior cingulate cortex (BA24), and caudate basal ganglia. Overall, 11 of the top 12 tissues ranked by expression were brain regions, confirming that these genes exert their primary biological effects within the central nervous system.

**Figure 3.**
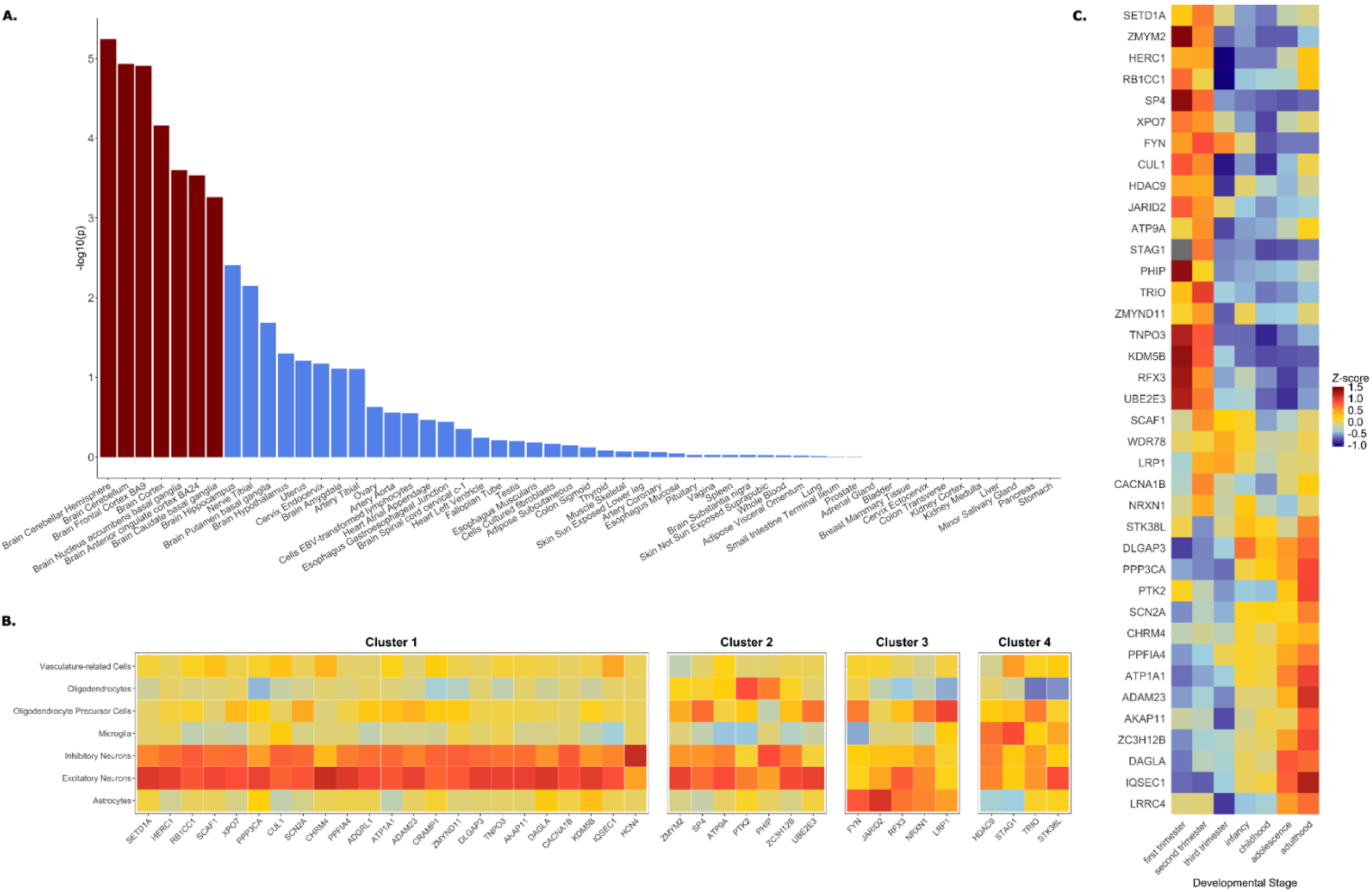
Expression patterns of SCHEMA 2.0 FDR 5% genes. **A.** Enrichment of SCHEMA genes in up-regulated differentially expressed gene (DEG) sets across GTEx tissue types. Bars represent –log₁₀(p-values) from a hypergeometric test assessing whether SCHEMA genes are significantly overrepresented among genes upregulated in each tissue (vs. all others), based on precomputed DEG sets in FUMA. **B.** Cell-type–specific expression of SCHEMA genes in the dorsolateral prefrontal cortex, based on aggregated single-nucleus RNA-sequencing data from 294 donors in the PsychENCODE project. Z-score–scaled expression values are shown across seven broad cell classes. K-means clustering identified four gene expression clusters, with distinct enrichment in neuronal, microglial, astrocytic, and OPC cell types. **C.** Developmental expression trajectories of SCHEMA genes across seven BrainSpan-defined stages. Z-score– scaled expression values were used to compare temporal patterns within each gene. Genes are ordered based on hierarchal clustering.

To resolve expression at the cellular level, we analyzed aggregated single-nucleus RNA-sequencing data from the PsychENCODE project, comprising dorsolateral prefrontal cortex (DLPFC) samples from 294 donors across eight cohorts, including both controls and individuals with psychiatric disorders. The 27 annotated cell types were grouped into seven broader classes, excitatory neurons, inhibitory neurons, astrocytes, microglia, vasculature-related cells, oligodendrocyte precursor cells (OPCs), and oligodendrocytes^21^. For each gene, we z-score scaled expression values to facilitate within-gene comparisons across cell types. All genes demonstrated elevated expression in both excitatory and inhibitory neurons (Figure 3B), a pattern that was consistently replicated at the individual cell type level (Supplementary Figures 24-25, Supplementary Table 11-12). Using k-means clustering, we identified four gene expression clusters: Cluster 1 (23 genes) showed strongest expression in excitatory and inhibitory neurons; Cluster 2 (seven genes) was marked by increased expression in oligodendrocytes and OPCs; Cluster 3 (five genes) included increased astrocyte expression; and Cluster 4 (four genes) comprised genes expressed in microglia. Together, these results suggest that although neuronal expression predominates across SCHEMA 2.0 genes, distinct expression clusters highlight heterogeneity in cellular specificity.

We next assessed gene expression across developmental stages using BrainSpan transcriptomic data^22^. Expression values were z-score scaled within each gene to allow within-gene comparison of expression trajectories over time, spanning seven developmental stages, first, second, and third trimesters; infancy; childhood; adolescence; and adulthood (Figure 3C). Genes were grouped into three developmental clusters based on their temporal expression profiles. The “early” cluster included genes, such as the transcriptional regulators *ZMYM2* and *SP4*, which were most highly expressed during prenatal development. The “late” cluster, which included *AKAP11* and *SCN2A*, showed peak expression during adolescence or adulthood. A smaller “mid” cluster was defined by peak expression during infancy or childhood. While these clusters highlight periods of maximal relative expression, it is important to note that many genes exhibit expression across multiple developmental stages (Supplementary Figure 26), as confirmed by raw TPM values.

### Cellular Functions

Among the exome-wide significant genes, we identified known cellular functions localized to cellular compartments (Figure 4). In the nucleus, several genes are involved in the regulation of gene expression. These include the chromatin modifier *SETD1A*, transcription factors *ZMYM2*, *SP4,* the histone deacetylase, *HDAC9,* and the histone demethylase, *JARID2*^23–29^. Additionally, *SCAF1* facilitates RNA polymerase binding and is predicted to play a role in RNA splicing^30,31^. Finally, *STAG1* functions as a component of the cohesion protein complex during cell division^32^.

**Figure 4.**
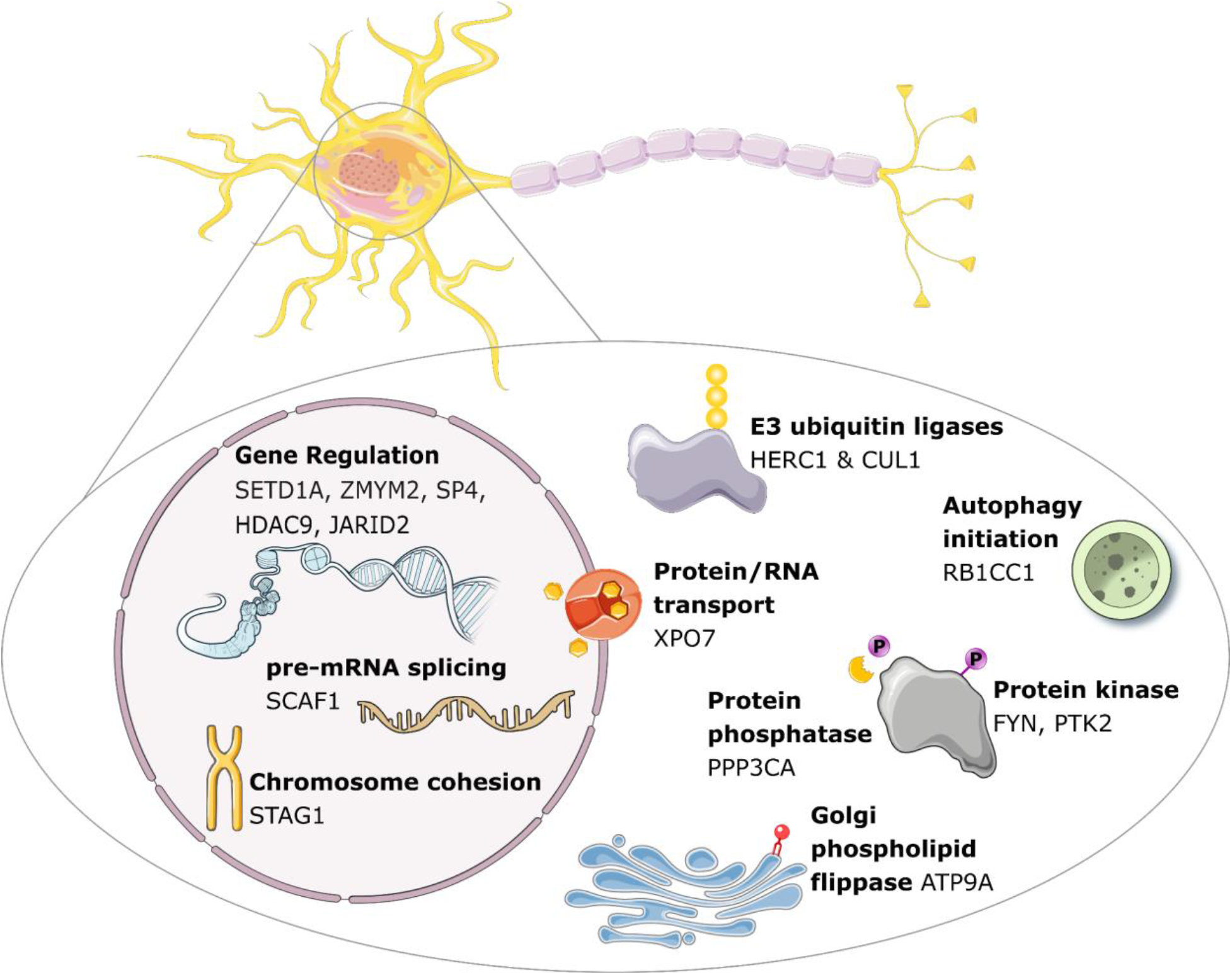
Cellular functions of exome-wide significant SCHEMA genes mapped to major neuronal compartments. In the nucleus, genes like SETD1A, ZMYM2, SP4, CUL1, and SCAF1 regulate chromatin structure, transcription, and RNA splicing. In the cytosol, XPO7, HERC1, and CUL1 are involved in RNA/protein transport and ubiquitin-mediated degradation. At the synapse, CACNA1B and SCN2A encode ion channel subunits, while NRXN1 and MAGI2 contribute to synaptic signaling and scaffolding. Illustrations were made using the NIH BioArt and SmartServier repositories.

In the cytosol, *XPO7* mediates the transport of proteins and large RNAs through nuclear pore receptors^33^, while *HERC1* and *CUL1* function as E3 ubiquitin ligases involved in initiating protein degradation^27,34,35^. *FYN* and *PTK2* function as tyrosine kinases and are currently being investigated as therapeutic targets and prognostic biomarkers across multiple cancer types, suggesting their potential as viable drug targets in other disease contexts^36,37^. Conversely, *PPP3CA* is a subunit of the protein phosphatase, calcineurin, and has been implicated in severe neurodevelopmental disorders and epilepsy^38,39^. *ATP9A* localizes to the Golgi apparatus and endosomes and belongs to a family of ATPases that act as phospholipid flippases between the exoplasm and cytoplasm^40^. Variants in *ATP9A* are catalogues in OMIM (entry: *609126) and have been associated with neurodevelopmental disorders^41^. Finally, *RB1CC1* is an autophagy initiation protein with a role in a variety of biological processes^42^.

We conducted gene ontology (GO) enrichment analysis using g:Profiler, a web-based tool that combines multiple databases^43^. The full set of g:Profiler results, including associations with GO:MF, KEGG, human disease annotations and protein-protein interactions, can be found in Supplementary Table 13. Using the 40 FDR 5% genes as input, we identified five biological processes and 13 cellular components with adjusted p-values < 0.05. The significant terms for biological processes included several signaling related processes, such as trans-synaptic signaling, synaptic signaling, and cell-cell signaling. The other significant processes were nervous system development and system development (Supplementary Figure 27). The top cellular components were the synapse, cell junction, postsynapse, and glutamatergic synapse (Supplementary Figure 27).

### Phenotypic Outcomes of SCHEMA 2.0 PTV Carriers in All of Us

We leveraged data from the All of Us Research Program (AoU) to investigate the phenotypic consequences of carrying variants in SCHEMA 2.0 genes using a phenome-wide association scan (PheWAS). Our analysis included 307,694 individuals from AoU with electronic health records, whole genome sequencing data, and demographic information. We restricted to variants with an AoU-specific MAF <1%, and further excluded variants with a population-maximum allele frequency of >1% in gnomAD v4, to remove variants that may have higher frequencies in ancestries underrepresented in AoU. Among the 40 SCHEMA 2.0 FDR 5% genes, we identified 1,581 PTV carriers in AoU.

In the PheWAS of PTV carriers, we identified 13 phenotypes meeting a Bonferroni correction for the number of phenotypes tested (p < 2.82e-5), including six psychiatric phenotypes. The most significant association was observed with personality disorders (p = 1.25e-7, OR = 2.24), followed by antisocial/borderline personality disorder (p=7.25e-7, OR = 2.60). We also identified associations with bipolar (p = 8.89e-6, OR = 1.55), tobacco use disorder (p = 1.13e-5, OR = 1.32), anxiety disorders (p= 1.60e-5, OR = 1.29), and suicidal ideation (p = 2.46e-5, OR = 1.82). The most significant non-psychiatric phenotypes identified were asthma (p = 8.07e-7, OR = 1.40), type 2 diabetes with neurological manifestations (p = 2.95e-6, OR = 1.64), and candidiasis of skin and nails (p = 4.43e-6, OR = 2.84) (Supplementary Figure 28A, Supplementary Table 14).

After adjusting for lifetime schizophrenia diagnosis, none of the psychiatric phenotypes remained, indicating these associations were driven by an underlying schizophrenia diagnosis. The only significant phenotypes were with type 2 diabetes with neurological manifestations, asthma, candidiasis of skin and nails, and septic shock Supplementary Figure 28B, Supplementary Table 15).

## Discussion

This study represents the largest rare coding variant association analysis of schizophrenia to date, tripling the case sample size from prior efforts and yielding a substantial increase in discovery. We identified 16 genes at exome-wide significance and an additional 24 passing FDR 5% significance, signifying an increased and refined gene list over previous efforts.

These results reinforce and expand upon findings from SCHEMA 1.0. Notably, 7 of the 10 exome-wide significant genes from SCHEMA 1.0 remain significant. The three genes no longer significant, *GRIA3, GRIN2A,* and *CACNA1G* in addition to decreased enrichment of PTVs in DD/ID genes, suggest a shift in schizophrenia case ascertainment away from comorbid DD/ID or milder schizophrenia cases in collections new to SCHEMA 2.0. In particular, null variants in *GRIN2A* are associated with early-onset schizophrenia, intellectual disability, and epilepsy^44^, which likely complicates recruitment of child-onset cases into large-scale schizophrenia cohorts. Alternatively, these differences may reflect technical heterogeneity between datasets, changes in variant calling or annotation, or inherent instability in rare-variant association analyses, including the possibility that some signals in SCHEMA 1.0 were false positives. Rare-variant analyses are particularly sensitive to small numbers of observed events, such that the addition or absence of a small number of variant carriers in expanded datasets can lead to substantial changes in effect estimates. This is a marked contrast to common variant association studies, where large allele counts provide much more statistical stability and reproducibility. While this study expands our understanding of schizophrenia genetic risk, the genes identified continue to be characterized by very low-frequency variants with high effect sizes (OR > 2), a pattern also seen in schizophrenia-associated copy number variants (CNVs). In contrast, common-variant GWAS has uncovered hundreds of common variants with small individual effects. To bridge this gap and uncover the full spectrum of genetic contributors, including identifying genes with intermediate frequency and moderate effects, further increases in sample size remain essential.

*CHRM4* emerged at a 5% FDR threshold, pointing to a muscarinic acetylcholine receptor with clear relevance to current schizophrenia treatment. This finding fits with the broader cholinergic hypothesis, which suggests that disruptions in muscarinic signaling contribute to the neural circuit abnormalities underlying psychosis^45^. The recent success of xanomeline-trospium (Cobenfy), a non-dopaminergic therapy that engages central muscarinic receptors, including M4, highlights the clinical relevance of this pathway^46,47^. At the same time, the Phase 2 failure of emraclidine^48^, a selective M4 agonist, suggests that targeting M4 alone may not be sufficient, and that combined actions at M1 and M4 receptors could be important for therapeutic benefit. Overall, identifying a known drug target among the top RVAS signals supports the ability of RVAS to highlight biologically meaningful and clinically relevant genes, and motivates further work to examine the biological and therapeutic potential of SCHEMA 2.0 findings.

Our findings replicate the substantial genetic overlap between schizophrenia and other neurodevelopmental and psychiatric disorders. Of the 40 SCHEMA genes, half also show associations with DD/ID, ASD, or bipolar disorder, underscoring a shared genetic architecture across these conditions. Notably, six genes, including *SETD1A*, *ZMYM2,* and *SCN2A,* are implicated in schizophrenia, DD/ID, and ASD, highlighting convergent biological pathways. However, a deeper analysis of *SCN2A* illustrates that while the same gene may be involved in multiple disorders, the specific mutation types differ, with schizophrenia enriched for damaging missense variants, and DD/ID enriched for more PTVs. These findings emphasize the importance of considering not only which genes are involved but also how the functional consequences of mutations may shape distinct phenotypic outcomes.

The SCHEMA 2.0 genes are all highly expressed in the brain, particularly within excitatory and inhibitory neurons. These results echo previous neurobiology findings of the alterations of neurons in induced pluripotent stem cells from schizophrenia patients, including decreased expression of postsynaptic density protein and glutamate receptors^49,50^ as well as changes to synaptic structure in postmortem brain tissue of schizophrenia patients^51^. Temporal expression of these genes in the brain find some genes are expressed predominantly in prenatal stages, while others have higher expression in adulthood. Functionally, they implicate diverse biological and cellular mechanisms, including chromatin remodeling (e.g., *SETD1A, SP4*), protein degradation (e.g., *HERC1, CUL1*), synaptic ion channels (e.g., *SCN2A, CACNA1B*), and neuronal receptor signaling (e.g., *NRXN1*). While this molecular diversity underscores the complex and heterogeneous biology underlying schizophrenia, the latest GWAS of schizophrenia established common variant enrichment of synaptic biology, strengthening the convergence of functional annotation between common and rare variant analysis.

Therapeutically, these findings highlight critical considerations for future drug development. First, because nearly all SCHEMA carriers are heterozygous, the implicated mechanisms likely involve partial loss of function, protein knockdown, rather than complete loss of gene activity, protein knockout. This distinction is essential when evaluating potential interventions, as it points toward reduced, but not absent, protein function. Importantly, our analysis does not account for compensatory expression from the reference allele^52^. Moreover, therapeutic strategies must account for developmental timing because genes expressed exclusively during early development are unlikely to be effectively targeted by treatments administered in adulthood. Target tissue specificity is also crucial to minimize off-target effects and maximize therapeutic benefit. Overall, a more complete understanding of expression, timing, and function in the identified genes will help enable therapeutic discovery.

Despite the strengths of this expanded dataset, several limitations must be acknowledged. Ascertainment strategies vary across cohorts, introducing heterogeneity in case definitions and control matching. In many cohorts, diagnostic data were collected decades ago, limiting our ability to assess diagnostic stability and disease trajectory. Integrating electronic health records into future designs will be a useful step for enabling long-term phenotypic follow-up. Additionally, changes in sequencing technology over the lifespan of this project presents further complexity, though we addressed this using harmonized quality control and association testing pipelines.

Finally, while the genes identified here have high odds ratios and robust statistical support, these results do not meet the criteria for application to clinical diagnosis and should not be misinterpreted as diagnostic tools. Many of these genes also harbor rare variant carriers among controls. As such, identifying a mutation in one of these genes is neither sufficient nor necessary to predict schizophrenia risk, with less than 2% of schizophrenia cases harboring a damaging mutation in an identified gene. Use of these findings for individual-level genetic testing would likely do more harm than good, potentially providing false reassurance or undue alarm to patients and families.

In conclusion, this study advances our understanding of schizophrenia genetics by leveraging ultra-rare coding variants, demonstrating overlap with neurodevelopmental pathways, and emphasizing the complex but actionable biology that underpins this disorder. Ongoing efforts to increase sample size, refine phenotyping, and integrate longitudinal data will be essential to elucidate the complex causes of schizophrenia and move towards new therapies.

## Code Availability

SCHEMA 2.0 analysis code can be found here: https://github.com/jsealock1/schema2

## Acknowledgements

We would like to extend gratitude to the patients and their families who made this work possible. We gratefully acknowledge *All of Us* participants for their contributions, without whom this research would not have been possible. We also thank the National Institutes of Health’s *All of Us* Research Program for making available the participant data examined in this study.

## Acknowledgements

J.M.S. This research was funded in part by BD2: Breakthrough Discoveries for thriving with Bipolar Disorder. For the purpose of open access, the author has applied a CC BY 4.0 public copyright license to all Author Accepted Manuscripts arising from this submission. C.A. Swiss National Science Foundation (P500-3_235131). M.A. Stanley Center, NIMH R01MH120642, NIMH U01MH125047. S.G. Stanley Center, NIMH U01MH125047. R.M.M. Stanley Center. C.R.J.C.N. Stanley Center. N.K. Stanley Center, NIMH R01MH120642. Z.Z. Stanley Center. J.K. Stanley Center, NIMH R01MH120642, NIMH U01MH125047. A.S. Stanley Center, NIMH U01MH125045. A.H. The authors would like to acknowledge and thank the study participants and their families. The GEN-SCRIP study is a public-private partnership between the National Institute of Mental Health (NIMH) and the Stanley Center for Psychiatric Research. This work was supported by a grant, R01 MH112904 (J.A.K), from the National Institute of Mental Health (NIMH). A.G. The author acknowledges financial support from the Brazilian Coordination for the Improvement of Higher Education Personnel (CAPES) through a research productivity scholarship. C.E.S.R. Stanley Center. M.A.S.J. Stanley Center. J.F. K08 MH118577. M.C.O.D. MCOD is funded by Medical Research Council Programme Grants (No. MR/Y004094/1 and No. MR/P005748/1). M.J.O. MJO is funded by Medical Research Council Programme Grants (No. MR/Y004094/1 and No. MR/P005748/1) and a Medical Research Council UK Centre grant (No. MR/L010305/1). E.R. ER is funded by a UKRI Future Leader Fellowship (No. MR/Y033922/1), a Medical Research Council Programme grant (No. MR/Y004094/1), The Brain and Genomics Hub of the Mental Health Platform Medical Research Council grant (No. MR/Z503745/1) and The Mental Health Goals Omics Project: A UK Multi-Omics Infrastructure and Resource for Psychosis and Mood Disorders grant (No. UKRI3712). P.F.S. Swedish Research Council (Vetenskapsrådet, award D0886501); NIMH R01 MH077139; NIMH U01 MH109528. J.T.R.W. JTRW is funded by Medical Research Council Programme Grants (No. MR/Y004094/1 and No. MR/P005748/1), The Brain and Genomics Hub of the Mental Health Platform Medical Research Council grant (No. MR/Z503745/1) and The Mental Health Goals Omics Project: A UK Multi-Omics Infrastructure and Resource for Psychosis and Mood Disorders grant (No. UKRI3712). B.C. Stanley Center. M.A. The authors would like to acknowledge and thank the study participants and their families. The GEN-SCRIP study is a public-private partnership between the National Institute of Mental Health (NIMH) and the Stanley Center for Psychiatric Research. This work was supported by a grant, R01 MH112904 (J.A.K), from the National Institute of Mental Health (NIMH). J.A.K. The authors would like to acknowledge and thank the study participants and their families. The GEN-SCRIP study is a public-private partnership between the National Institute of Mental Health (NIMH) and the Stanley Center for Psychiatric Research. This work was supported by a grant, R01 MH112904 (J.A.K), from the National Institute of Mental Health (NIMH). R.S. Stanley Center, NIMH R01MH120642, NIMH U01MH125045. L.A. Stanley Center, NIMH U01MH125047. A.D. Stanley Center, NIMH R01MH120642, NIMH U01MH125047. S.M.K. Stanley Center. K.C.K. Stanley Center, NIMH R01MH120642, NIMH U01MH125045, NIMH U01MH125047. D.J.S. Stanley Center, NIMH R01MH120642. S.T. Stanley Center, NIMH R01MH120642, NIMH U01MH125047. H.H. acknowledges support from Merkin Institute Fellow, the Zhengxu and Ying He Foundation, and the Stanley Center. This research was funded in part by BD2: Breakthrough Discoveries for thriving with Bipolar Disorder. For the purpose of open access, the author has applied a CC BY 4.0 public copyright license to all Author Accepted Manuscripts arising from this submission. B.M.N. This research was funded in part by BD2: Breakthrough Discoveries for thriving with Bipolar Disorder. For the purpose of open access, the author has applied a CC BY 4.0 public copyright license to all Author Accepted Manuscripts arising from this submission. This research was also supported by R01MH101244, R37MH107649, U01MH125047, U24MH140953. The authors note with sadness that D.J.S. (Dan J. Stein) passed away during the preparation of this manuscript.

## Online Methods

### Samples and Quality Control

As part of the Populations Underrepresented in Mental Health Association Studies (PUMAS) and the Stanley Center Global Initiative, we sequenced the exomes of cases and controls from around the globe using the newly developed blended genome exome (BGE) method^53^. The BGE method generates deep whole-exome data (30-40x mean depth) along with low-pass whole genome data (1-4x mean depth) in a single sequencing run. For SCHEMA 2.0, we used only the deep exome component of the BGE data in our analyses.

SCHEMA 2.0 contained samples from 5 different callsets: gnomAD exomes, gnomAD genomes, Cardiff/UCLA, BGE-GATK, and BGE-DRAGEN. Detailed quality control procedures are described in the Supplementary Material. Briefly, we performed within-callset sample quality control for the BGE datasets and the Cardiff callset, while samples from gnomAD were used as provided following gnomAD-standard quality control. We excluded samples with outlier values on standard sample-level metrics, sex discrepancies, high contamination or chimeric read rates, or evidence of relatedness. After combining samples across callsets, we removed related individuals up to the second degree and conducted principal component–based matching of cases and controls within each exome capture platform to control for population structure.

### Variant annotation

To identify damaging variants, we annotated all variants using Ensembl’s Variant Effect Predictor (VEP) v95(ref:^54^). We defined predicted protein-truncating variants (PTVs) as variants with predicted effects of stop-gained, frameshift, splice acceptor, or splice donor variants that also passed LOFTEE high-confidence filters^8^. We defined damaging missense variants by combining annotations from MPC, AlphaMisssense Pathogenicity, MisFit-S, and PopEVE^9–11,55^ by first rank-ordering variants in each annotation from most to least deleterious, calculating the mean rank for each variant, and then using the rank-ordered percentile as the missense mean rank annotation. These missense scoring methods were chosen due to their higher schizophrenia enrichment reported by GeneticsGym^56^. To define a threshold to select damaging missense variants, we evaluated each percentile for enrichment of missense singletons in schizophrenia cases versus controls, using only data not represented in gnomAD to avoid bias from the missense prediction algorithms using gnomAD in their training. Using the enrichment analyses, we selected variants ≥ 93^rd^ percentile for missense mean rank.

### Rare Variant Association Study (RVAS)

Given heterogeneity in ancestry, exome capture platforms, and variant callsets across the SCHEMA 2.0 dataset, we performed rare variant association analyses using a Cochran-Mantel-Haenszel (CMH) framework, stratifying by ancestry and exome capture kit combination. Strata were required to include at least 50 cases and 50 controls to ensure stable estimation. For each gene, we conducted two burden tests: one including protein-truncating variants (PTVs) only and a second including the combined burden of PTVs and damaging missense variants. For each gene, we selected the CMH result with the lowest (i.e., most significant) p-value. Odds ratios were calculated using the Mantel-Haenzel common odds ratio, and confidence intervals were calculated using the Robbins-Breslow-Greenland variance estimator.

Statistical significance was assessed using both Bonferroni and false discovery rate (FDR) correction. Exome-wide Bonferroni correction was defined as 0.05 divided by the total number of gene tests performed across both PTV-only and PTV plus damaging missense analyses. FDR control at 5% was implemented by pooling p-values from both analyses and applying a global FDR procedure. Sensitivity analyses included restricting tests to singleton variants, excluding variants with population maximum allele frequency (popmax) greater than 0.1% in gnomAD, and excluding variants with popmax greater than 0.1% in the Regeneron Genetics Center dataset, where popmax denotes the maximum allele frequency observed within any ancestry group. To assess potential test statistic inflation, we additionally performed CMH burden tests of synonymous singleton variants.

### Overlap with Autism, Developmental Delay/Intellectual Disability, and Bipolar

We curated gene sets associated with developmental delay/intellectual disability (DD/ID), autism spectrum disorder (ASD), and bipolar disorder (BIP) from published large-scale exome sequencing studies. DD/ID genes were obtained from Kapalanis et al. (2020), ASD genes from the most recent ASD exome analysis (Fu & Satterstrom et al.), and bipolar disorder genes from the latest BipEx study (Liao et al.). SCHEMA 2.0 samples were restricted to genes within each disorder-specific gene set, and protein-truncating variant (PTV) singletons were selected.

For each disease bin, we performed logistic regression to test the association between the burden of PTV singletons across all genes in the bin and schizophrenia case-control status. All models included covariates for ancestry group, exome capture, callset, and the top five principal components to control for population structure and technical confounders.

### Expression analysis

Expression analyses were conducted using multiple publicly available transcriptomic resources. Bulk tissue expression in GTEx was assessed using the FUMA GENE2FUNC platform (https://fuma.ctglab.nl/). Single-cell expression data were obtained from the PsychENCODE resource via PsychScreen (https://psychscreen.wenglab.org/psychscreen), and expression values were z-score scaled within each gene across cell types to facilitate relative enrichment comparisons. Developmental expression data were sourced from the BrainSpan atlas; samples were grouped into developmentally relevant age bins, and expression was z-score scaled within each gene across age groups.

### Cellular Functions

Functional enrichment analysis was performed using the g:Profiler web tool (https://biit.cs.ut.ee/gprofiler/gost). Enrichment significance was assessed using the default g:SCS multiple-testing correction. Analyses were first conducted using all 40 SCHEMA genes reaching FDR 5% significance, and subsequently repeated for each gene cluster identified through the PsychENCODE expression–based clustering analysis.

### PheWAS

Analyses were conducted using the All of Us Research Program Controlled Tier Dataset (v8). Protein-truncating variant (PTV) and missense variant carriers with minor allele frequency <1% were identified across the 40 SCHEMA genes using publicly available gnomAD VEP-annotated variant files. Carrier status was then tested in a phenome-wide association study (PheWAS), with models adjusted for sex, median age, ancestry, and the first 10 principal components to account for population structure.

