## Supplemental Methods, Figures for "Analysis of Rare Coding Variation Identifies New Genetic Contributors to Schizophrenia"

### SCHEMA 2.0 Supplementary Methods & Figures

Full SCHEMA Consortium Authorship, Alphabetical ... 47
Cohort Descriptions. ... 55
References ... 68

#### Sequencing Generation

We sought to integrate whole exome and whole genome sequencing across cohorts and sequencing technologies to enable gene discovery in schizophrenia. We obtained individual level sequencing data from five separate callset sources: gnomAD v4.0 exomes, gnomAD v4.0 whole genomes, Cardiff + UCLA, the Blended Genome Exome-GATK (BGE-GATK), and the Blended Genome Exome-DRAGEN (BGE-DRAGEN). All data, except for the Cardiff/UCLA data, were sequenced at the Broad Institute. Sequencing data used in SCHEMA 1.0 were previously described in Singh et al 2020[1].

The Cardiff/UCLA sequencing data were reprocessed at the Broad Institute through joint calling using GATK best practices and aligned onto the human genome reference build 38 (GRCh38). Description of sequencing of the Cardiff dataset was described by Chick et al 2025[2].

DNA samples used in BGE (BGE-GATK and BGE-DRAGEN) were extracted from saliva and whole blood specimens outside of our lab. Following DNA extraction, the BGE protocol was followed, as described by Boltz et al 2025[3]. Briefly, DNA was first normalized to 50 ng/µL, transferred into a 384-well plate, and purified using a 2.75X SPRI clean-up with Ampure XP beads. After quantification by spectrophotometry, DNA was renormalized to 25 ng/µL and 134 ng was used for PCR-free library preparation with custom NEBNext Ultra II FS kits under reduced fragmentation/end-repair/A-tailing conditions (37 °C for 42.57 min, 65 °C for 30 min). Unique dual-indexed adaptors were ligated at 20 °C for 20 min, followed by two SPRI size selections (0.5X and 0.55X). Libraries were quantified by qPCR, normalized to ~0.3–2 nM, pooled, and concentrated. For exome preparation, aliquots of the pooled PCR-free libraries were PCR-amplified (12 cycles, 98 °C/65 °C), purified by 1X SPRI clean-up, normalized to 70 ng/µL, and pooled for hybridization capture with Twist Alliance Clinical Research Exome probes using the xGen capture protocol. PCR-free and exome-captured pools were quantified on the same qPCR run, blended at 67% WGS and 33% WES, and requantified for sequencer loading. The final blended pools of uniquely barcoded samples were sequenced on an Illumina NovaSeq S4 with 2x150 bp runs for BGE-GATK, or Illumina NovaSeqX for BGE-DRAGEN.

#### Variant Joint Calling

Joint calling of SCHEMA samples was performed using either the Genome Analysis Toolkit[4] (GATK) or Illumina DRAGEN[5], for BGE-DRAGEN.

For samples using GATK, it was used to perform local realignment around indels and recalibrate base qualities in each sample BAM. We called each sample using HaplotypeCaller, generating gVCF files containing every position of the genome with likelihoods for variants or the genomic reference. For each GATK callset, (gnomAD exomes, gnomAD genomes, Cardiff/UCLA, and BGE-GATK), we merged samples using CombineVCFs, and joint-called samples using GenotypeVCFs, all using default settings according to the best-practice pipeline. We annotated all variants using the Variant Quality Score Recalibration (VQSR) tool in GATK.

For samples using Illumina DRAGEN, alignment and variant calling were performed with the DRAGEN Bio-IT Platform, which provides hardware-accelerated algorithms for read mapping, duplicate marking, local realignment, and base quality recalibration within a single pipeline. Variants were called per sample in gVCF mode, capturing both variant and non-variant positions with associated likelihoods. For the DRAGEN callset, BGE-DRAGEN, sample gVCFs were merged and joint genotyping was performed using the Genomic Variant Store (GVS) joint-calling workflow under default best-practice parameters.

#### Quality Control of Sequencing Data

We used five different callsets for our analysis: the gnomAD exomes, gnomAD genomes, Cardiff/UCLA, BGE-GATK, and BGE-DRAGEN.

##### Variants

gnomAD Exomes and Genomes

Exome and whole-genome sequencing (WGS) cohorts were extracted from gnomAD version 4.0. The extracted subsets underwent quality control as detailed by gnomAD. For the WGS data, sites were filtered to the gnomAD exome interval list to maintain consistency with the exome data.

After extracting samples of interest, we performed Allele-Specific VQSR on exomes and genomes separately and removed failing variants. We filtered to 'adjusted' genotypes with sequencing depth (DP) ≥ 10, genotype quality (GQ) ≥ 20, and heterozygous allele balance (AB) > 0.20. Finally, we removed variants with call rates <0.90 in each callset (Supplementary Figure 1).

Blended Genome Exome-GATK & Cardiff/UCLA

For the Cardiff/UCLA and BGE data we obtained raw sequencing data rather than a pre-quality-controlled subset. We therefore conducted our own quality control to identify high quality sites, genotypes, and samples. To account for differences in data generation and joint calling, we conducted quality control separately for each callset. We applied the following quality control on variants and genotypes:

At the site level, we removed sites with more than 6 alternate alleles present, outside of the Twist exome capture target intervals or within low complexity regions. We split multi-allelic sites into biallelic sites and removed variants failing Allele-Specific VQSR. We removed invariant sites and calculated variant call rates. Variants within the BGE callset with call rates <0.95 were removed; variants within the Cardiff/UCLA callset with call rates <0.98 were removed (Supplementary Figure 1).

We performed the same genotype quality control on both callsets. We removed genotype calls with genotype quality (GQ) scores < 20 or read depth (DP) < 10. In heterozygous genotype calls, removed calls if the allele balance (AB) < 0.2 or if (allele depth of reference + allele depth of alternate)/total DP < 0.8. In homozygous alternate calls, we removed if AB < 0.8.

Blended Genome Exome-DRAGEN

A newer iteration of the BGE sequencing product included processing through Illumina's DRAGEN software[5] (BGE-DRAGEN). DRAGEN uses multigenome mapping with pangenome references and machine learning-based variant detection with low computational burden. Additionally, DRAGEN data is through the Genomic Variant Store (GVS), rather than the Hail VDS. The change to GVS limits the transferability of the BGE-GATK QC described above due to GVS's lack of read depth information and upper limiting of genotype quality to 50.

First, we used Hail to write the GVS as a Hail matrix table. Next, we removed sites with more than 6 alternate alleles present, outside of the Twist exome capture target intervals or within low complexity regions. We split multi-allelic sites into biallelic sites and removed variants failing VariantExtractTrainScore (VETS) algorithm, which is the functional equivalent of GATK's VQSR for DRAGEN data. We removed invariant sites and removed variants with call rates < 0.95.

We summed the allele depths from the allele balance field for heterozygote and homozygote alternate calls to estimate read depth. In DRAGEN data, homozygote reference calls do not contain allele balance or depth information. Therefore, to perform genotype QC, we required homozygous reference calls to have a genotype quality of at least 25. In heterozygous calls, we removed calls with GQ < 25, estimated DP < 12 or allele balance < 0.25. In homozygous alternate calls, we removed calls with GQ < 25 or estimated DP < 12.

**Supplementary Figure 1**. A) Histograms and B) Cumulative distribution of variant call rates by callset. We applied a call rate filter of $\geq 0.9$ for gnomAD genomes, gnomAD exomes, and BGE callsets, and $\geq 0.98$ for Cardiff/UCLA.

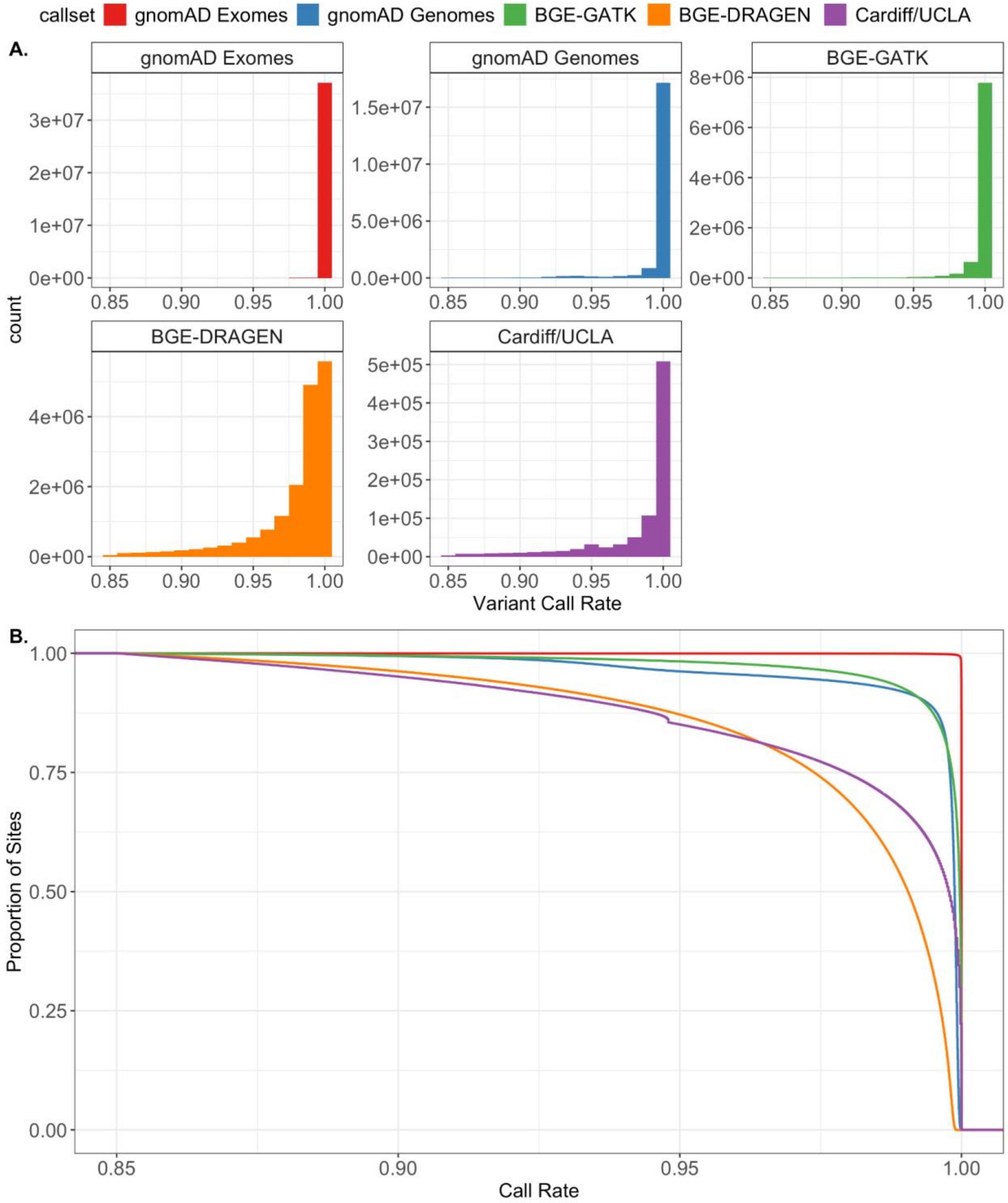

**Samples**

gnomAD Exomes and Genomes

For the exomes and genomes extracted from gnomAD, we used samples labelled as 'high-quality' that passed all of gnomAD's sample quality control. gnomAD's sample QC included removing samples failing hard filters for contamination, number of singletons, heterozygosity ratio, number of bases with high coverage, chimeric read rate, call rate on high quality sites, sex discordance and relatedness. Additionally, outliers for indel ratio, transition to transversion ratio, singleton to transversion ratio, and heterozygosity ratio were removed after stratifying for platform and regressing out the top 20 principal components.

gnomAD provided inferred genetic ancestry by using hwe_normalized_pca() to cluster samples based on genetic similarity over 30 principal components on high-quality sites. The top 20 PCs were used to train a random forest classifier and assign ancestry based on minimum probability from the classifier. We removed samples without a genetic ancestry label.

BGE-GATK

Using the exome portion of the BGE-GATK, we restricted to high-quality samples based on hard filters, ancestry assignment, and outlier filtering. First, we removed samples with greater than 5% chimeric read rate or contamination rates.

Next, we assigned genetic ancestry using a combination of PCA with a random forest algorithm. Given the unique genetic diversity of the BGE data, we used a custom reference panel to infer the genetic ancestry of the BGE-GATK samples. We combined the Human Genome Diversity Project (HGDP) and the 1,000 Genomes Project phase 3 data with the African Wits-INDEPTH Partnership for the Genomic Study of Body Composition and Cardiometabolic Disease Risk (AWI-Gen) and African Genome Variation Project (AGVP). We combined the BGE data with the reference data and filtered for variants with call rate > 99% and minor allele frequency > 0.1%. We LD-pruned the combined data to find independent variants and then calculated the top 10 PCs using Hail's hwe_nomalized_pca() function. Using the top 10 PCs, we trained a random forest model on the custom reference panel samples using continental ancestry as the outcome and applied the model to the BGE samples. We required samples to have a minimum probability of 70% for a single ancestry to be assigned ancestry. Samples failing the 70% threshold for a single ancestry were removed.

On samples with assigned ancestry, we used Hail's sample_qc() function to compute sample quality metrics. Within each cohort and ancestry combination, we removed samples that had values 4 median absolute deviations from the median for the following sample quality metrics: transition to transversion ratio (Ti/Tv), N insertions, N deletions, N transitions, N transversions, heterozygosity ratio, indel ratio, and N SNP. For N singletons, we removed samples with values more than 4 MADs below or 8 MADs above the median, due to the lower bound of 0 for singletons.

To remove samples with discordant sex, we filtered to biallelic variants on chromosome X with call rates > 0.99 and allele frequency > 0.1%. We LD-pruned the remaining variants, and used Hail's impute_sex() function to calculate the F-statistic on the variants outside of the pseudo-autosomal region. Samples with an F-statistic less than 0.6 were assigned female and samples with F-statistic greater than 0.6 were assigned male. Samples with discordant genetic sex and reported gender were removed.

**Supplementary Figure 2.** Sample quality control of BGE-GATK data restricted to passing samples A) PC 1 vs PC2 with reference panel (colors) and BGE-GATK samples (black) stratified by cohort. B) distribution of contamination rates, C) distribution of chimeric read rates, D) concordance between reported gender and imputed genetic sex, and E) Sample quality control metrics.

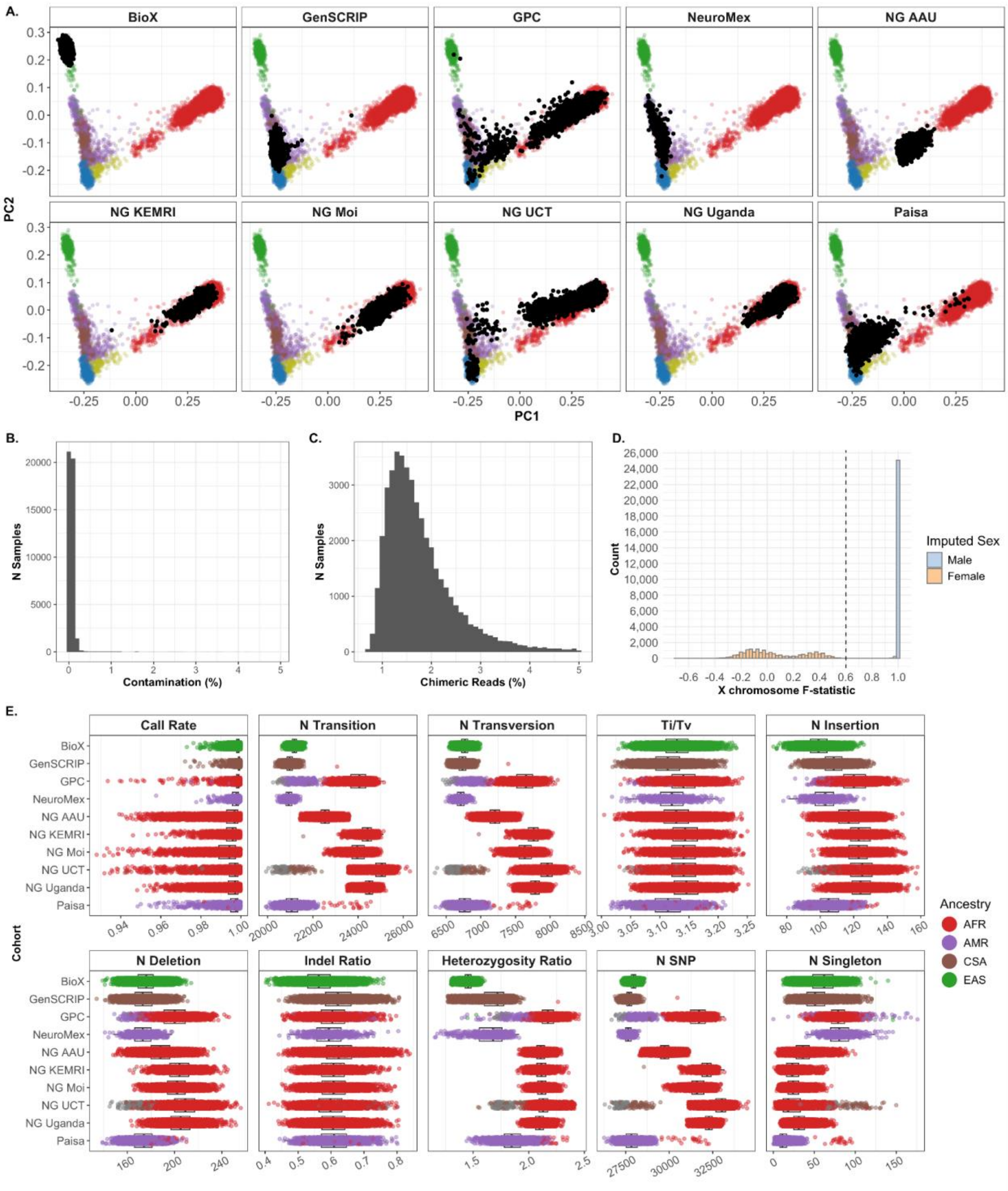

BGE-DRAGEN

We performed similar sample QC methods for the BGE-DRAGEN callset, with slight changes to fit the DRAGEN data format.

First, we removed samples with greater than 5% chimeric read rate or contamination rates. Next, we assigned genetic ancestry with PCA and a random forest algorithm. We combined the BGE data with the HGDP and 1KGP phase 3 reference data and filtered for variants with call rate > 90% and minor allele frequency > 0.5%. We LD-pruned the combined data to find independent variants and then calculated the top 10 PCs using Hail's hwe_nomalized_pca() function. Using the top 10 PCs, we trained a random forest model on the reference panel samples using continental ancestry as the outcome and applied the model to the BGE-DRAGEN samples. We required samples to have a minimum probability of 70% for a single ancestry to be assigned ancestry. Samples failing the 70% threshold for a single ancestry were removed (Supplementary Figure 3A).

On samples with assigned ancestry, we used Hail's sample_qc() function to compute sample quality metrics. Within each cohort and ancestry combination, we removed samples that had values 4 median absolute deviations from the median for the following sample quality metrics: transition to transversion ratio (Ti/Tv), N insertions, N deletions, N transitions, N transversions, heterozygosity ratio, indel ratio, and N SNP. For N singletons, we removed samples with values more than 4 MADs below or 8 MADs above (Supplementary Figure 3D).

Due to the structure of the DRAGEN data, we could not use the standard Hail impute_sex() function to find genetic sex. Instead, we calculated the number of chrY genotype calls for each sample. Samples with < 2,000 genotype calls were labeled female, and samples with > 9,000 genotype calls were labeled males (Supplementary Figure 3C). Samples with discordant genetic sex and reported gender were removed.

**Supplementary Figure 3.** Sample quality control of BGE-DRAGEN data restricted to passing samples A) PC 1 vs PC2 with reference panel (colors) and BGE-DRAGEN samples (black) stratified by cohort. B) distribution of contamination values C) distribution of chimeric read rates, D) imputed genetic sex using number of chrY calls, and E) Sample quality control metrics.

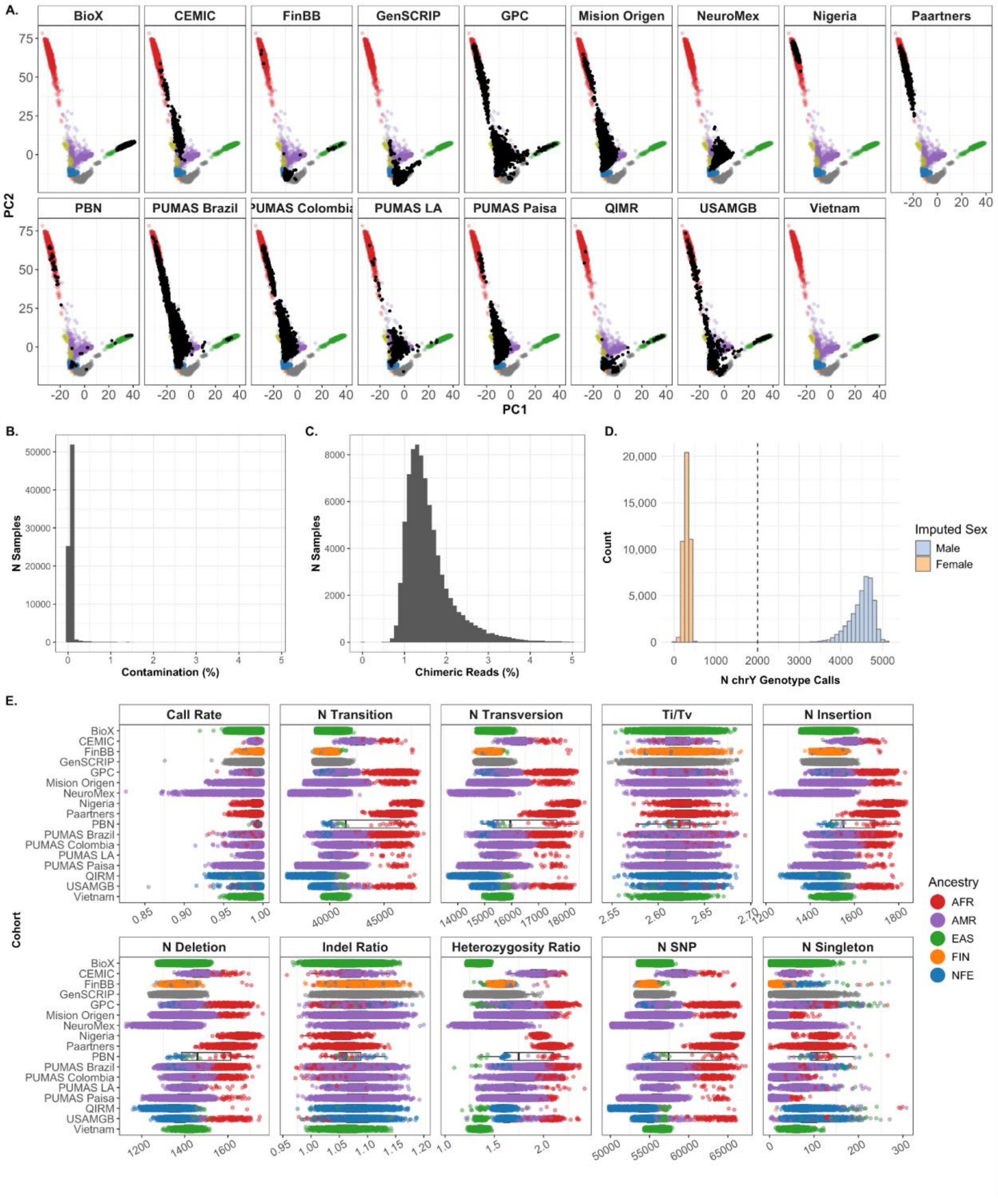

Cardiff/UCLA

In the Cardiff/UCLA callset, we completed sample QC similar to the BGE data. Due to the samples in this callset being externally sequenced, we did not have contamination or chimeric read rates available for these samples. Instead, we used CHARR. We removed samples with CHARR values greater than 5%.

To assign ancestry, we combined the Cardiff/UCLA callset with the HGDP and 1KGP reference dataset and used the gnomAD v3.1 PCA loadings to define common and independent variants. We further filtered to variants with a call rate > 98% and minor allele frequency > 0.1% and calculated the top 10 PCs using Hail's hwe_nomalized_pca() function. We trained a random forest model on the reference dataset samples and then applied the model to the Cardiff/UCLA samples. We required samples to have a minimum probability of 70% for a single ancestry assignment. Samples with probability < 70% were removed.

As described in the BGE data, we used Hail's sample_qc() function to compute sample quality metrics on the Cardiff/UCLA samples. Within each cohort and ancestry combination, we removed samples that had values 4 median absolute deviations from the median for the following sample quality metrics: transition to transversion ratio (Ti/Tv), N insertions, N deletions, N transitions, N transversions, heterozygosity ratio, indel ratio, and N SNP. For N singletons, we removed samples with values more than 4 MADs below or 8 MADs above the median.

To remove samples with discordant sex, we filtered to biallelic variants on chromosome X with allele frequency > 1%. We LD-pruned the remaining variants, and used Hail's impute_sex() function to calculate the F-statistic on the variants outside of the pseudo-autosomal region. Samples with an F-statistic less than 0.6 were assigned female and samples with F-statistic greater than 0.6 were assigned male. Samples with discordant genetic sex and reported gender were removed.

**Supplementary Figure 4.** Sample QC of the Cardiff & UCLA callset. A) Histogram of CHARR contamination values less than 5%, B) PC1 vs PC2 with reference panel in colors and samples in black, C) sex and gender concordance, and D) sample quality metric distributions by cohort and ancestry.

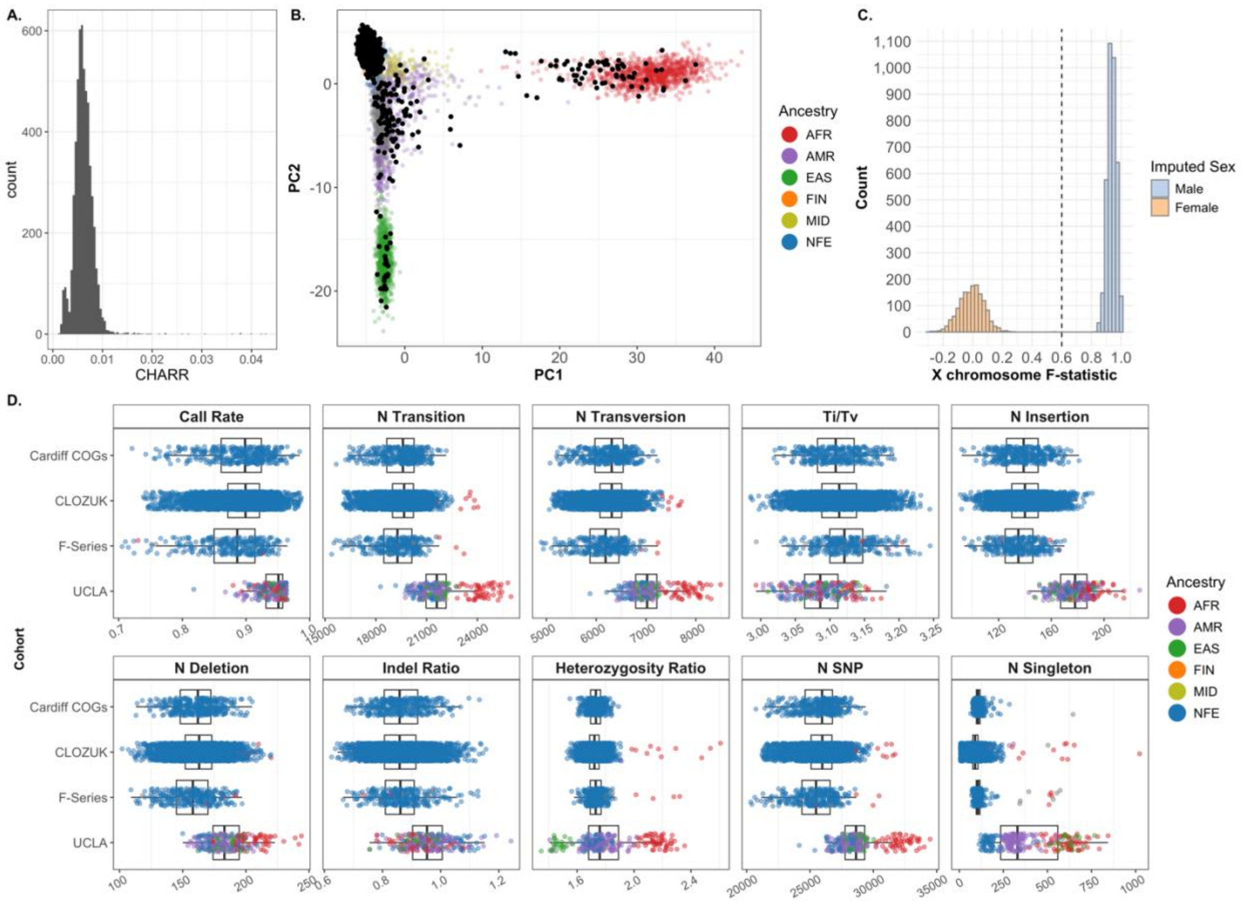

Combined Sample QC

Following callset-specific quality control, we merged all passing samples from each callset into a single Hail matrix table, comprising 68,924 cases and 128,123 controls. We then assessed the quality of the samples within this combined matrix table. Given that each callset was joint-called separately, the resulting matrix displayed patterns of missingness across variants, which could introduce bias into the sample QC metrics and complicate cross-callset comparisons. To mitigate this, we filtered the data to retain only high-quality variants, defined as variants with a call rate > 0.99, minor allele frequency > 0.01%, and a Hardy-Weinberg equilibrium p-value > 1e-8, yielding 115,638 variants. We used these high-quality variants to calculate sample quality metrics via Hail's sample_qc() function. We visually inspected these metrics to identify any sample outliers or issues related to sample QC. Based on this visual inspection, we removed samples with a call rate < 0.9 (94 samples from gnomAD exomes callset). The passing samples and their QC metrics are presented in Supplementary Figure 5.

**Supplementary Figure 5.** Sample quality control metrics by callset and ancestry calculated on high-quality sites.

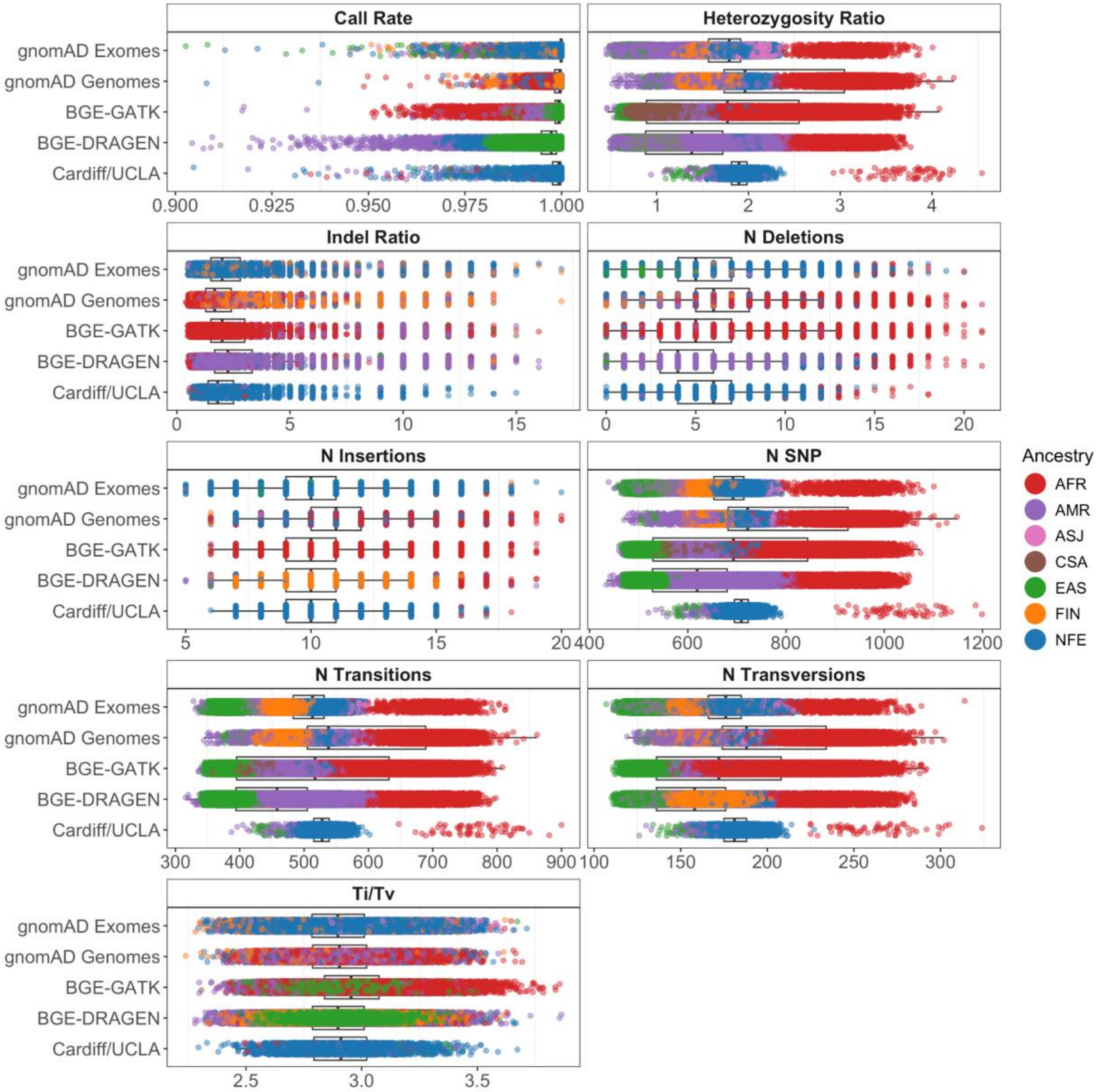

Relatedness

Inclusion of duplicate, or first- or second-degree relatives in a population-based RVAS can inflate rare variant counts and provide inaccurate reflections of population allele frequencies. Prior to defining rare variants and conducting the RVAS, we filtered the SCHEMA 2 dataset for related samples across all callsets. On the combined dataset, we filtered to biallelic variants with minor allele frequency > 0.5% and call rate > 90%. Next, we filtered to LD-pruned variants in the gnomAD v4.0 PCA loadings. We exported the matrix table as PLINK binary files and used PLINK 2.0 to run KING to calculate pairwise relatedness across all samples (Supplementary Figure 6).

We filtered to pairs with a kinship of > 0.0884, representing duplicates, first-degree, or second-degree relatives. From each related pair, we removed one sample using Hail's maximal_independent_set function.

**Supplementary Figure 6.** Histogram of pairwise kinship coefficients for pairs with kinship > 0.05.

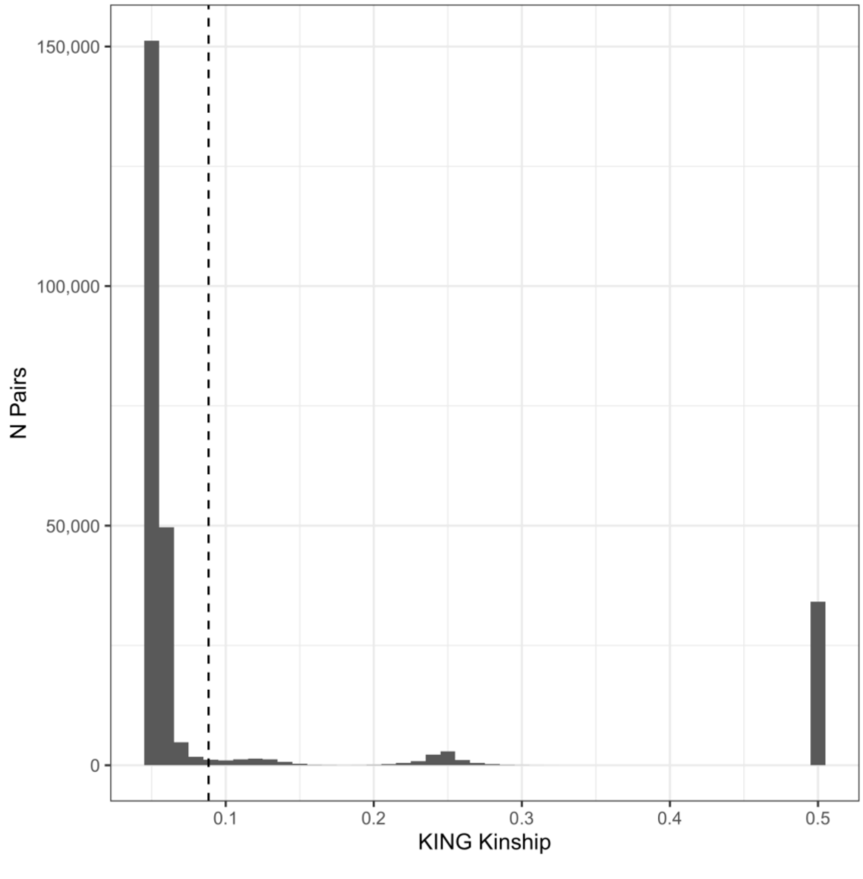

Ancestry Matching

We sought to match cases and controls using principal components (PCs) to ensure ancestry-matched samples for analysis. Using the unrelated samples, we filtered to biallelic variants with MAF > 0.5% and call rate >99%. Next, we filtered to LD-pruned variants in the gnomAD v4.0 PCA loadings, leaving 8,051 variants. Using the hwe_normalized_pca function in Hail, we calculated the top 10 PCs across all samples (Supplementary Figure 7).

Within each capture, we required each ancestry group to have at least 50 cases and 50 controls. Because the Cardiff/UCLA samples were sequenced using the Nextera capture kit, we combined these samples with Nextera samples from the gnomAD exomes callset. Prior to filtering, the dataset contained 82,591 cases and 141,160 controls. After filtering, we retained 82,485 cases and 138,979 controls (Supplementary Figure 8).

**Supplementary Figure 7.** PC1 vs PC2 of all SCHEMA 2 samples.

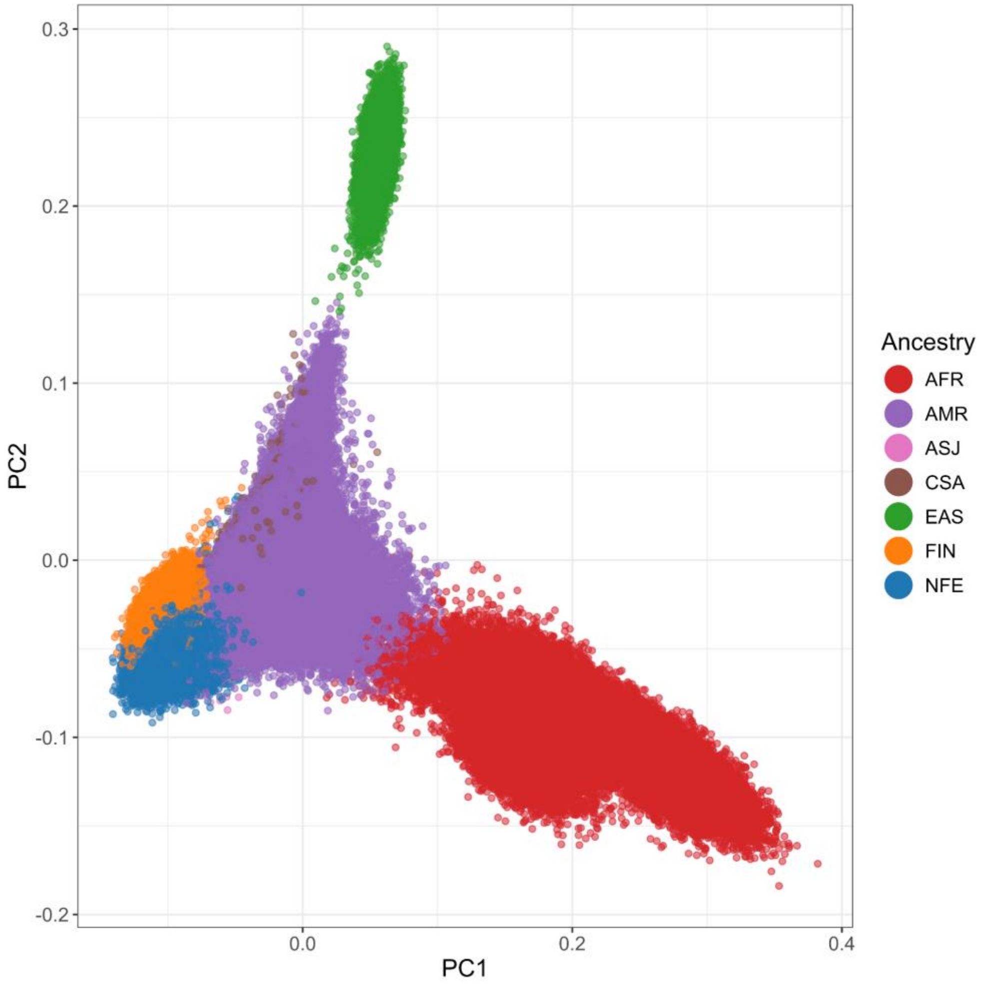

**Supplementary Figure 8.** PC1 vs PC2 of samples A) before and B) after ancestry matching within capture.

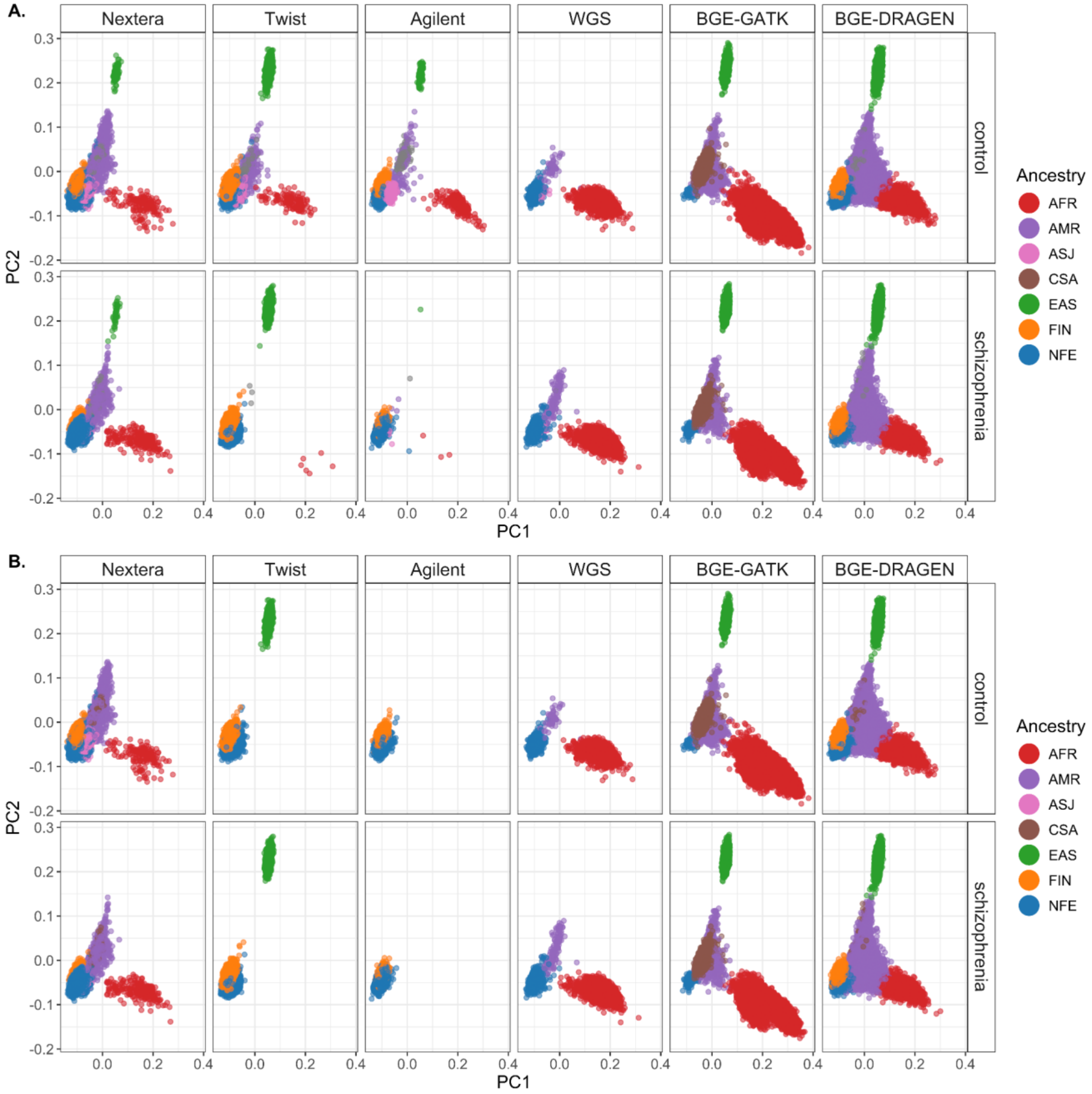

#### Variant Annotation

We used the Ensembl Variant Effect Predictor (VEP) v95 to annotate all variants with minor allele count ≤ 10. All predicted variant consequences were defined using the GENCODE canonical transcript.

##### Protein-truncating variants (PTVs)

We considered variants with 'transcript ablation', splice acceptor, splice donor, stop-gained, and frameshift variants as protein truncating variants (PTVs). To determine high-confidence protein-truncating variants, we implemented the LOFTEE annotation with minor adaptations[7]. After filtering to variants deemed high-confidence PTVs by LOFTEE, we removed variants with flags that indicated a single exon gene or variants where no exon number is indicated due to overlapping an intron. Additionally, due to missingness considerations with GERP scores in GRCh38 LOFTEE, we removed PTVs that fail the 50 base pair rule and have a GERP distance of less than zero, indicating variants that are the end of a transcript and likely not protein truncating that also have low conservation scores.

##### Missense variants

To classify the deleteriousness of missense variants, we used multiple scoring methods, including MPC, AlphaMissense Pathogenicity, PopEVE, and MisFit-S. While these scores were all developed using similar datasets (e.g., gnomAD, UK Biobank), no single method consistently outperforms the others across all variants. To integrate information across these methods, we created a missense mean rank score by ranking each missense variant from most to least deleterious within each method, then averaging the ranks across all available scores.

Next, we sought to identify the optimal threshold of missense mean rank to use in the rare variant analyses. Because each of the missense scoring methods used gnomAD to generate the scores, we focused on samples not included in gnomAD, using the BGE-GATK and BGE-DRAGEN datasets as independent testing sets. For each percentile of the missense mean rank score, we assessed the enrichment of singleton missense variants in schizophrenia cases versus controls (Supplementary Figure 9A). We also performed enrichment tests across binned percentiles: 1–90 (as a negative control), and ≥90, ≥91, ≥93, and ≥94 (Supplementary Figure 9B). Based on these results, we selected the ≥93rd percentile as the threshold for defining damaging missense variants in the rare variant analysis.

**Supplementary Figure 9.** Enrichment testing of missense mean ranking percentiles in BGE data. We evaluated the enrichment of singleton missense variants in schizophrenia cases versus controls in BGE data A. within each missense mean rank percentile and B. binned by percentiles. The dashed line in panel A represents the 93rd percentile.

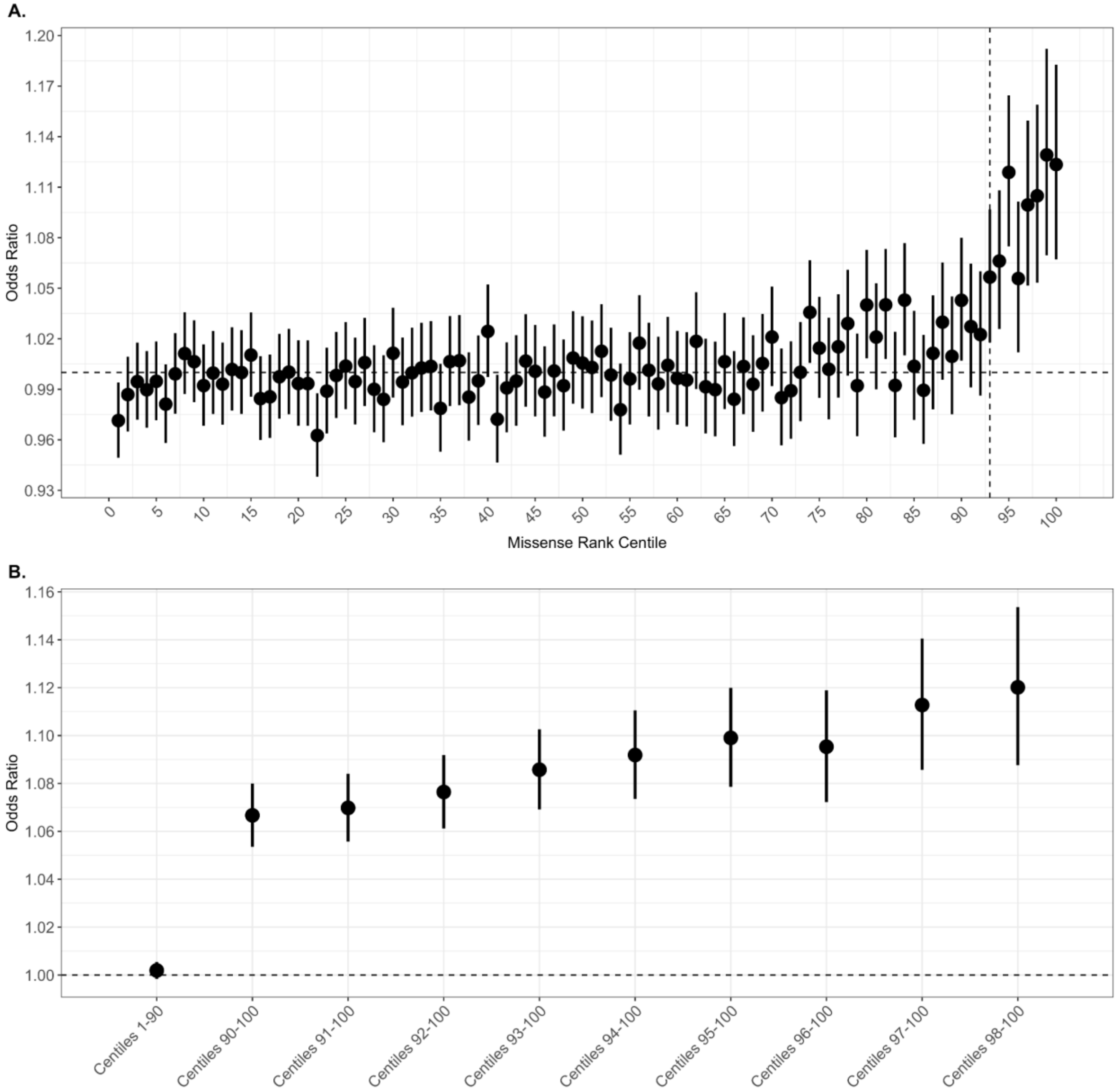

**Genotype Quality Control for RVAS**

While the genotyping QC described above is effective for general QC purposes, we sought to ensure that the rare variants included in our analysis were genuine rather than sequencing artifacts. To achieve this, we applied highly stringent genotyping filtering criteria prior to rare variant analysis.

To identify the correct genotyping filtering criteria, we focused on high-confidence pLoF variants with a minor allele count ≤ 15 in constrained genes to ensure the filters were relevant to the variants targeted in RVAS. We visualized the genotype quality (GQ) and allele balance (AB) for each callset in the combined matrix table as well as within each capture in the gnomAD exomes data (Supplementary Figure 10).

Most genotypes exhibited GQ ≥ 50 and AB ≥ 0.25, with those having lower GQ also showing lower AB. By design, DRAGEN calls were limited to an upper GQ threshold of 50, while callsets from GATK (all others) were limited to an upper GQ threshold of 100.

Based on these observations, we restricted our analysis to genotypes with GQ ≥ 25 and AB ≥ 0.25 for all rare variant analyses.

**Supplementary Figure 10.** Histograms of allele balance binned by genotype quality (GQ) for high-confidence pLoF variants in constrained genes stratified by A) callset and B) capture kit within the gnomAD Exomes callset.

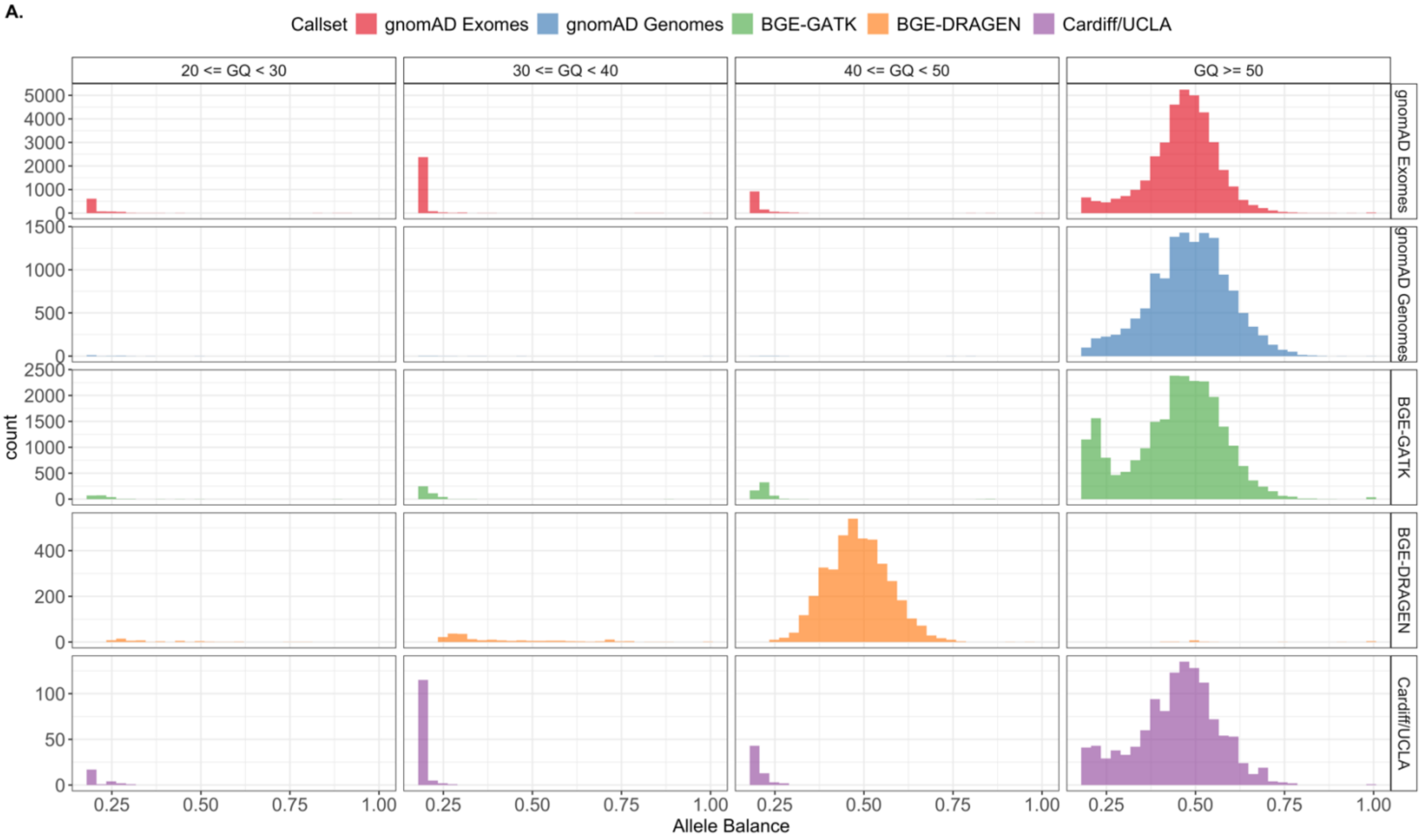

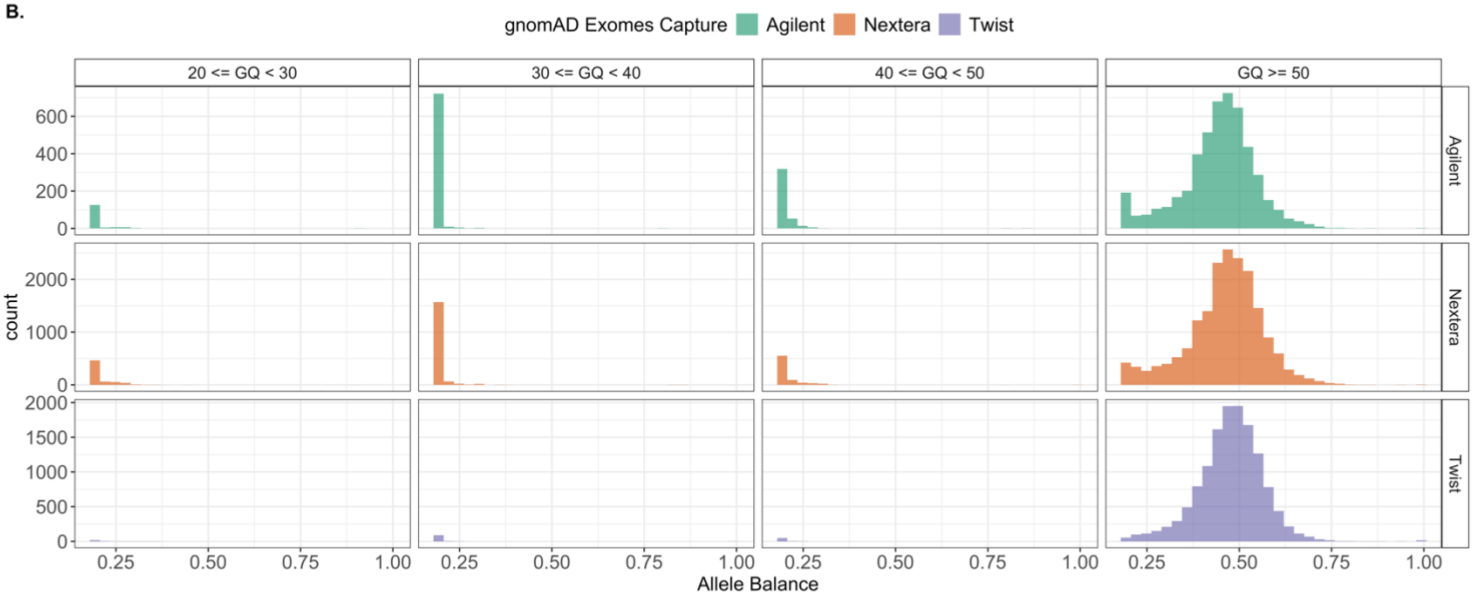

#### Enrichment of Damaging URVs

Previous studies have shown that schizophrenia cases are enriched for damaging coding variants in constrained genes compared to controls[1,18]. To inform our choice of minor allele count threshold for the RVAS, we compared enrichment of damaging URVs in schizophrenia cases versus controls across MAC groups.

First, we investigated the distribution of MAC across damaging variants. From the combined matrix table, we filtered to pLoF and damaging variants with MAC ≤ 10 and plotted the distributions in Supplementary Figure 11. Most variants were singletons (67.6% for PTV, and 73.3% for damaging missense). As the MAC increases, the number of variants within the bin decreases, with only 6.5% of PTV and 3.7% of missense variants with 6 ≤ MAC ≤ 15.

Next, we stratified the counts by MAC bin and number of ancestries (Supplementary Figure 12) to determine if any URVs are overrepresented in a single ancestry. Overall, most variants tend to be shared across ancestries. In MAC = 5, over half (51%) of the variants are in more than one ancestry. This proportion increases in 60% and 65% of variants in MAC = 10 and MAC = 15, respectively. Additionally, we calculated the population-specific allele frequencies of all variants with MAC < 100 in our dataset (Supplementary Figure 13). Other than ASJ, as noted in the main text, no ancestry exhibited variants with frequencies > 0.1%, confirming we are evaluating variants that appear to be rare across all ancestries.

From the combined matrix table, we restricted to constrained genes, defined as genes with pLI > 0.9 from the gnomAD v4.0 constraint calculations. Next, we binned variants by MAC grouping: singletons, variants with 2 ≤ MAC ≤ 5, variants with 6 ≤ MAC ≤ 10, and variants with 11 ≤ MAC ≤ 15. We grouped variants into three annotation categories: PTVs, damaging missense with missense mean rank >= 93%, and synonymous. Within each MAC bin, we summed the number of URVs present per sample per annotation group. Additionally, we calculated the total number of singletons per individual across the entire genome, not restricted to constrained genes, to control for genome-wide mutation rate.

For each ancestry group, we projected PCs from the gnomAD v4.0 PCA loadings to use as covariates in the enrichment models.

Within each ancestry group and capture, we tested for enrichment in each annotation group and MAC bin by regressing schizophrenia status on the total number of variants of interest, controlling for the total number of synonymous singletons across the genome and the top five principal components. We combined results across captures within the same ancestry and MAC group using a fixed effects inverse-weighted meta-analysis, giving ancestry-statified results. Finally, we combined results within the same MAC bin and across ancestries using a fixed effects inverse-weighted meta-analysis, resulting in a single estimate per annotation per MAC bin (Supplementary Figure 14A). To illustrate the ancestry-agnostic burden of damaging URVs in schizophrenia cases, we plotted the enrichment of singletons in schizophrenia cases stratified by ancestry in Supplementary Figure 14B.

**Supplementary Figure 11**. Histograms of minor allele count for A) PTV and B) damaging missense variants.

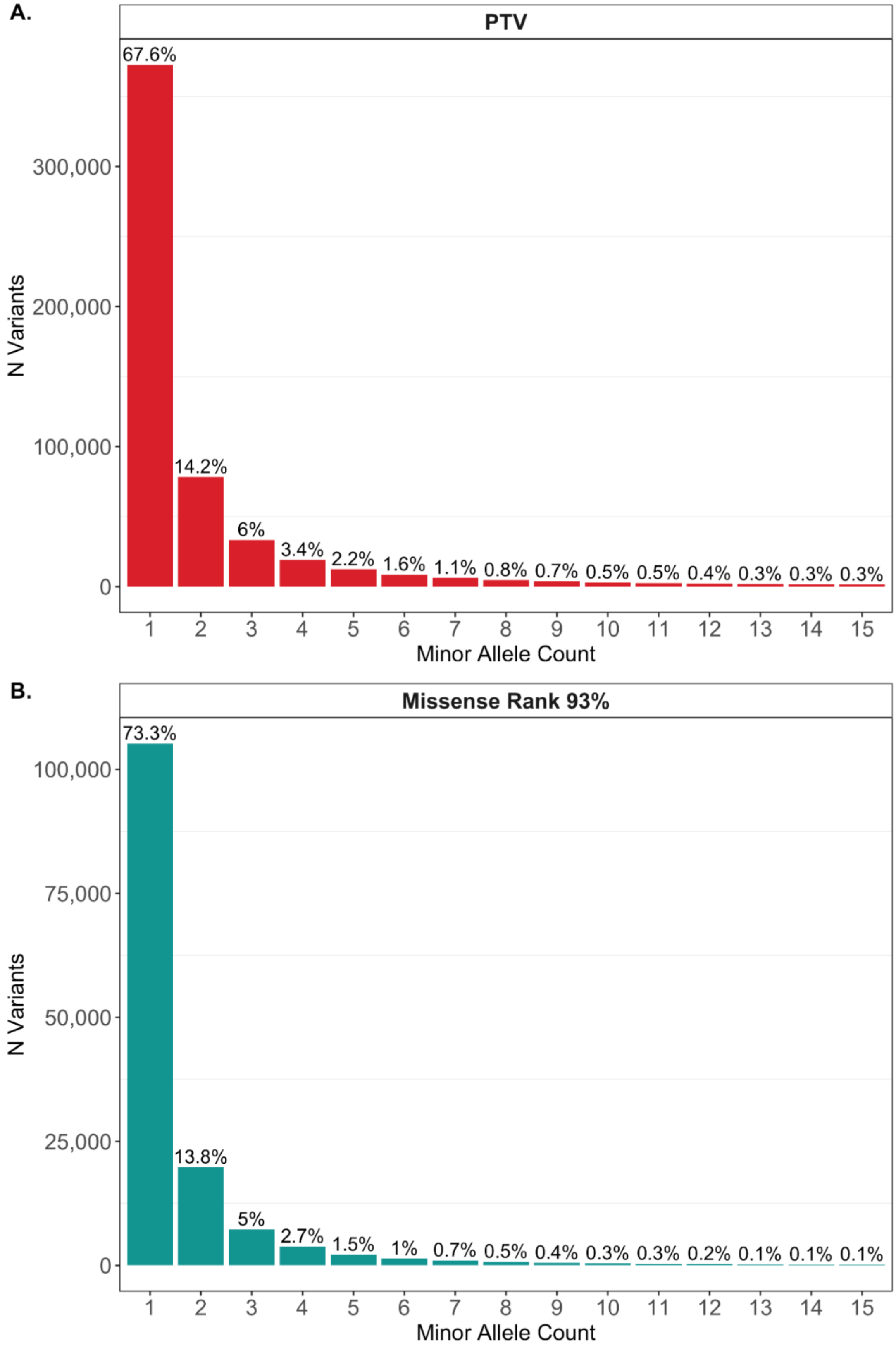

**Supplementary Figure 12.** Number of ancestries per MAC bin with carriers in A) PTVs and B) damaging missense variants.

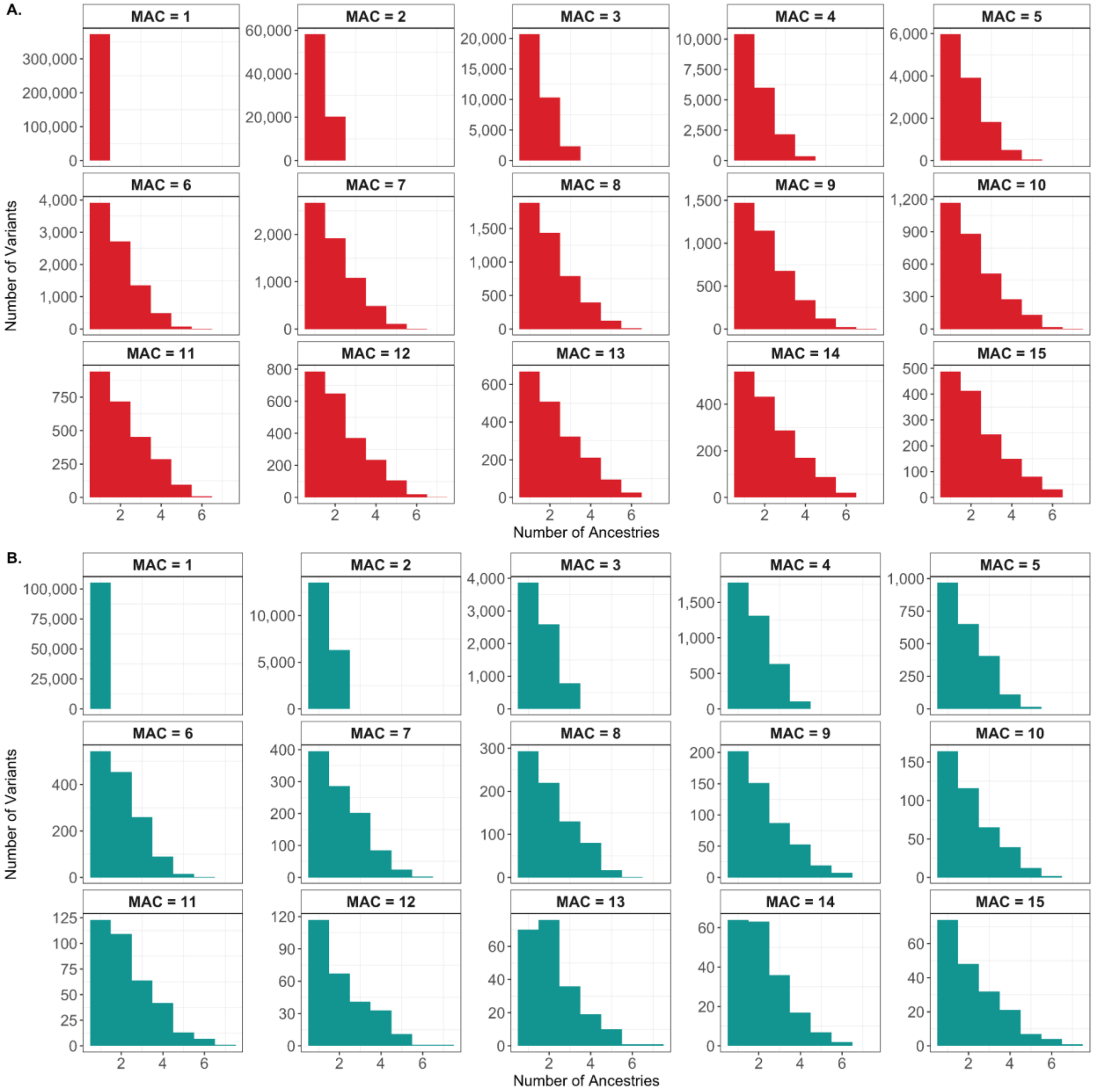

**Supplementary Figure 13.** Histograms of ancestry-stratified minor allele frequencies for PTVs with MAC < 100. The dashed lines indicate a MAF of 0.1% within each ancestry.

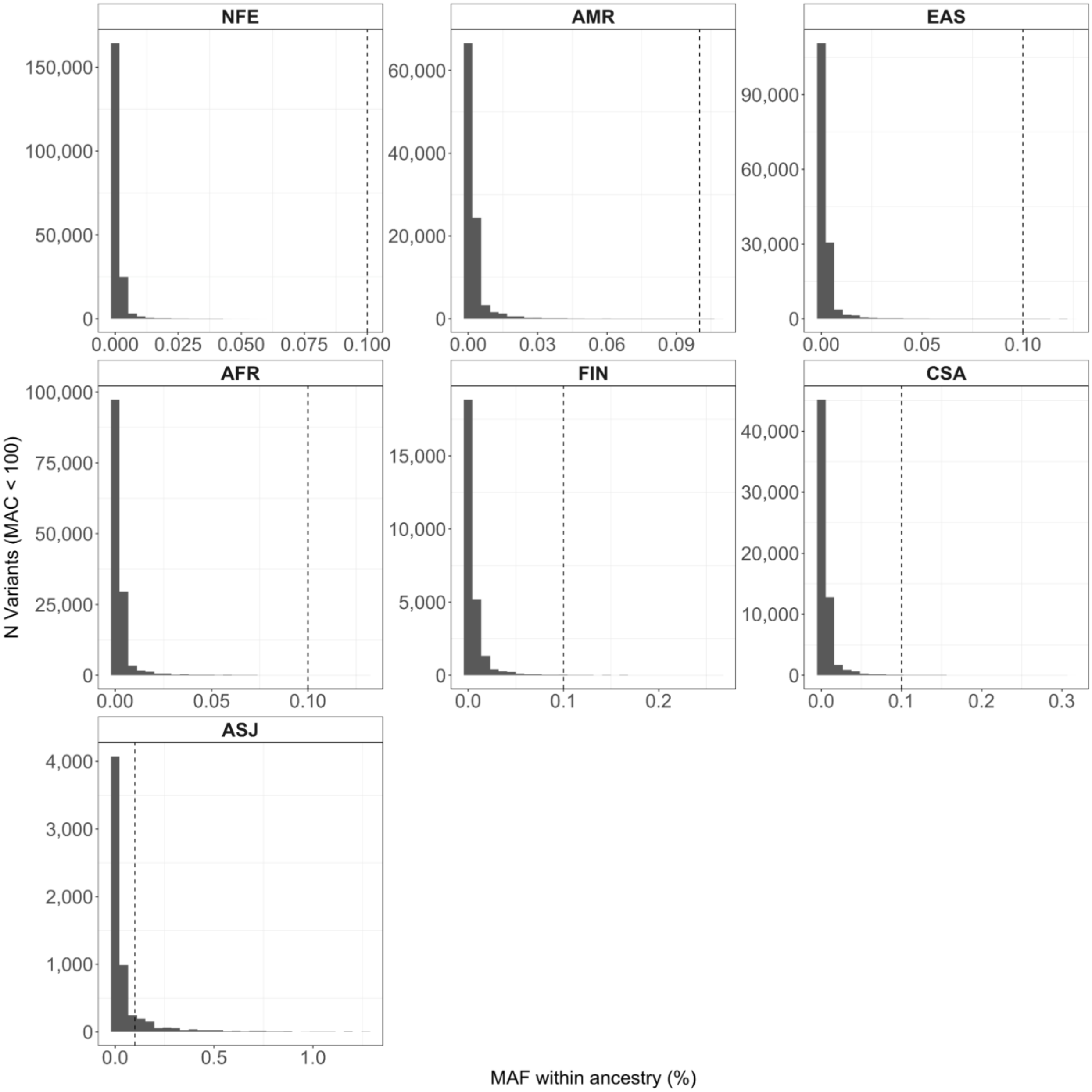

**Supplementary Figure 14.** Enrichment of rare, damaging mutations. A) enrichment of ultra-rare variants stratified by minor allele count (MAC) bin and annotation. B) enrichment of singletons by annotation and ancestry. Ancestries are ordered from largest case sample size to smallest.

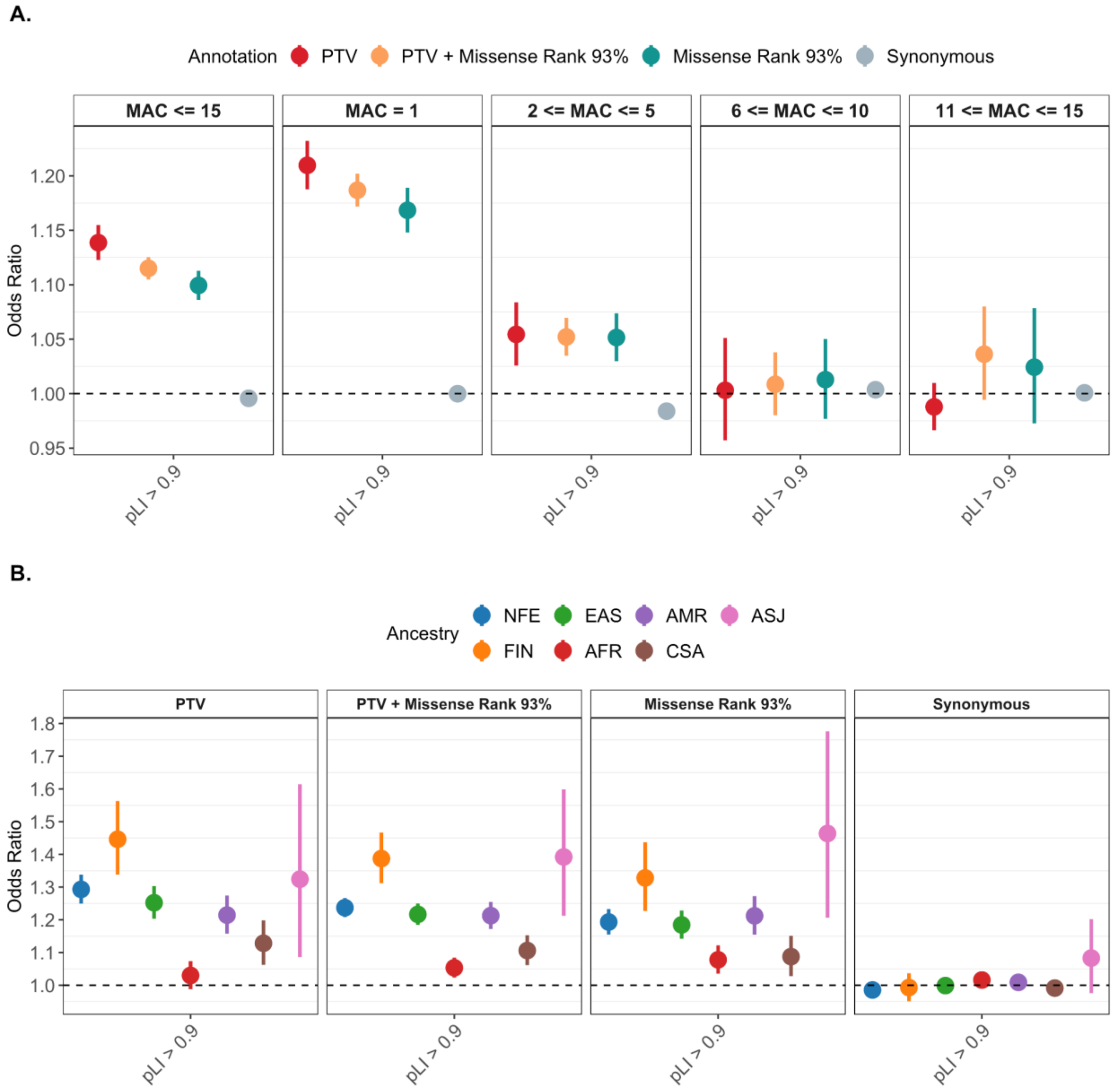

#### Rare Variant Association Testing

##### Case-control association testing

Two cohorts from SCHEMA 1.0, iPSYCH and UK10K, are no longer available to use with individual level sequences. To maintain the data from these two cohorts for SCHEMA 2.0, we extracted the variant counts for these two cohorts from SCHEMA 1.0 to include in the RVAS. We lifted over the coordinates from GRCh37 to GRCh38 and annotated as described for the remaining case-control data.

Using samples matched on ancestry, we first conducted the case-control association testing with CMH tests on variants with MAC ≤ 15 using ancestry and capture as strata. We conducted CMH tests for each annotation separately: PTV only, PTV + damaging missense variants, and synonymous. We visually inspected quantile-quantile plots of the CMH p-values for each annotation (Supplementary Figure 14). As expected, the majority of the signal comes from the PTV variants. However, the combined analysis of PTV + damaging missense variants results in a noticeable boost in power. For each gene in the case-control analyses, we conducted a maximum of two association tests, PTV + missense or PTV only. For each gene, we selected the most significant p-value between these two annotations.

Overall, we tested 18,864 genes in the PTV only tests and 19,389 genes in the PTV + missense tests. We set Bonferroni significance at $1.31 \times 10^{-6}$ (0.05 / 18,864 + 19,389). We determined the FDR 5% significance threshold using a distribution of all tests ($p = 6.87 \times 10^{-5}$).

##### Incorporating *de novo* data

Updating *De Novo* Variant Coordinates and Annotations

SCHEMA 1.0 included the incorporation of *de novo* mutations from 10 published schizophrenia trio studies of 3,402 parent-proband trios[8–17]. Using the *de novo* mutations identified in SCHEMA 1.0, we updated the variant information and annotation to match our current analyses. First, we lifted over the variant coordinates from GRCh37 to GRCh38. Next, we updated variant annotations to match the case-control analysis by running Ensembl VEP v95, and annotating using LOFTEE and missense mean rank scores.

##### Modeling *De Novo* Variants

In alignment with SCHEMA 1.0, we modeled *de novo* mutations as the Poisson probability of observing >= N *de novo* mutations in a gene given the baseline gene and annotation-specific mutation rate from gnomAD v2. We calculated Poisson p-values for pLoF variants only, damaging missense missense variants only (mean rank percentile ≥ 93) and the combination of pLoFs and damaging missense variants. Among the maximum of three tests per gene, we selected the minimum p-value to carry forward into meta-analysis with the case-control data.

##### Meta-analysis with *de novo* data

In SCHEMA 1.0, a substantial proportion of the association signal was driven by *de novo* results from 3,402 trios. With a nearly fourfold increase in case numbers in SCHEMA 2.0, the incremental contribution of the trio data is comparatively small. For this reason, we did not include *de novo* results in the primary analyses; instead, we incorporated them into a sensitivity analysis by meta-analyzing case-control and *de novo* p-value using a weighted Stouffer's method. In SCHEMA 1.0, the case-control p-values were assigned a weight of 2. For SCHEMA

2.0, we evaluated multiple weighting schemes and selected the weight at which the number of significant genes plateaued (weight = 10).

**Supplementary Figure 15**. QQ plots for each meta-analysis step by annotation. The PTV panel represents the results for the PTV only CMH test and PTV + Missense are the results from the PTV and damaging missense CMH test. The Lowest Pvalue panel is the most significant pvalue between the PTV only and PTV + damaging missense tests. The De Novo Meta-analysis panel is after conducting a weighted meta-analysis between the most significant case-control pvalue and the *de novo* pvalues.

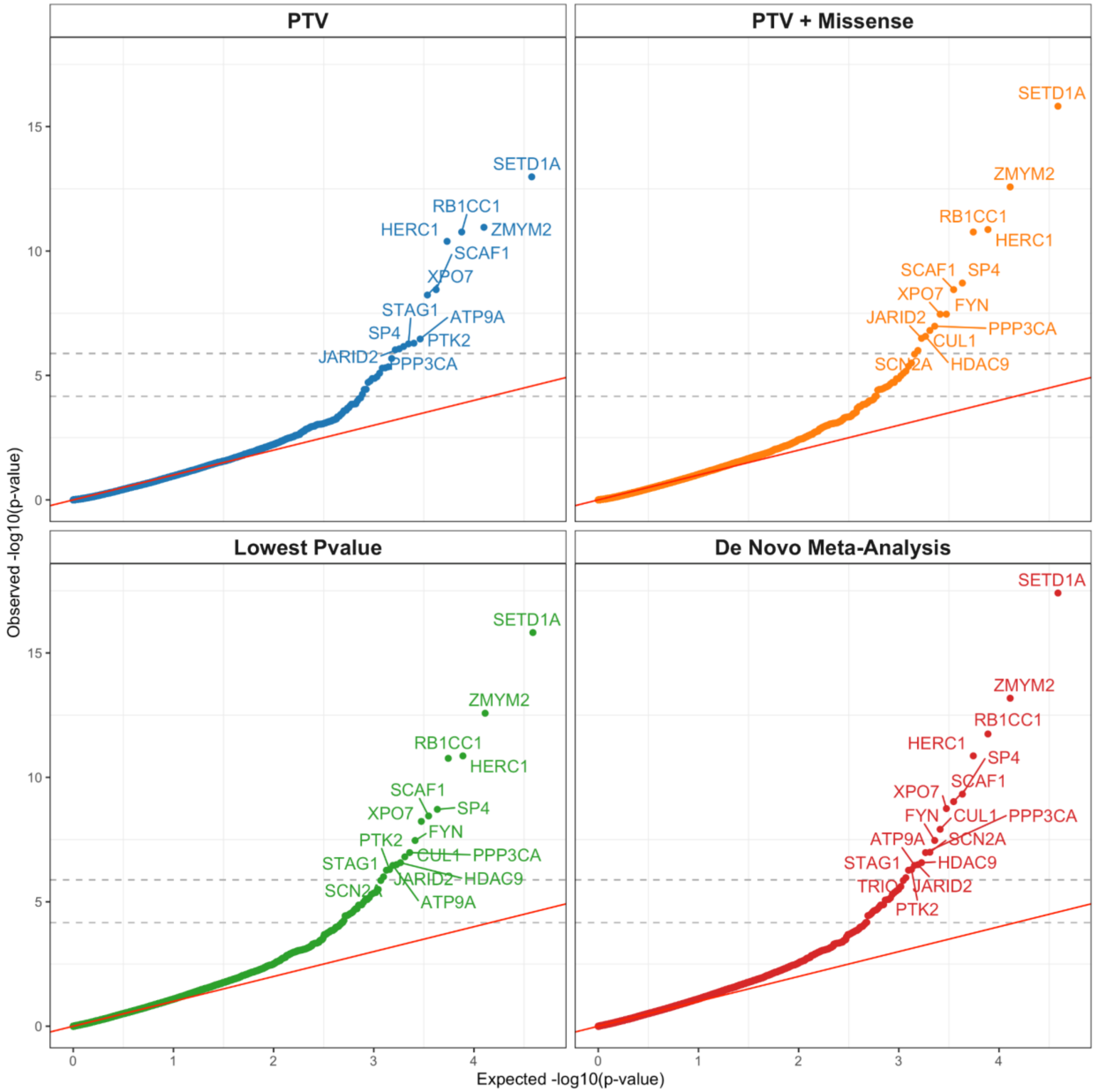

#### Sensitivity Analyses

To evaluate the robustness of the RVAS results and assess potential sources of bias, we conducted several sensitivity analyses on the case-control data.

##### Synonymous Singleton RVAS

As a negative control to verify data quality and model calibration, we performed a CMH test on synonymous singleton variants. Because synonymous variants are not expected to influence schizophrenia risk, this analysis should yield no association signal if quality control procedures and confounder adjustment are adequate. To avoid instability driven by genes with very small carrier counts, we further filtered genes based on the number of synonymous carriers and PTV carriers per gene, applying minimum carrier thresholds of 1, 20, 50, and 100. Across all thresholds, we observed no evidence of systematic inflation in the synonymous singleton results, supporting the adequacy of sample and variant quality control and indicating that the CMH framework was well controlled for potential confounders (Supplementary Figure 16).

##### Singleton RVAS Comparison

Given that the enrichment analyses indicated that the majority of the rare variant association signal was driven by singleton variants, we assessed the concordance of gene-level association results between singleton-only analyses and analyses including variants with minor allele count (MAC) ≤ 15. We repeated the rare variant association study (RVAS) using only singleton PTVs and damaging missense variants and compared the resulting gene-level p-values to those obtained from the MAC ≤ 15 analysis (Supplementary Figure 17). Rank-based correlation analysis demonstrated strong overall concordance between the two approaches (Spearman's $\rho = 0.39$, $p < 2.2 \times 10^{-16}$). Concordance was further increased when restricting to genes reaching a false discovery rate (FDR) of 5% ($\rho = 0.67$, $p = 2.68 \times 10^{-4}$). Notably, 8 of the 40 genes significant at FDR 5% exhibited more significant p-values in the singleton-only analysis than in the MAC ≤ 15 analysis, most prominently *LRP1* (MAC ≤ 15 p-value = $2.79 \times 10^{-5}$; singleton p-value = $1.80 \times 10^{-8}$). Overall, however, most genes demonstrated increased statistical power when including additional rare variants up to MAC ≤ 15.

##### Filtering by Population Frequency

To assess whether variants with elevated population-specific allele frequencies influenced the association results, we performed additional analyses excluding variants with a population maximum (popmax) allele frequency greater than 0.1% in either gnomAD v4 or the Regeneron Genetic Center (RGC) Million Exome Variant Browser. RVAS results obtained after applying each filter were highly consistent with the unfiltered analyses, indicating that variants with higher population frequencies were not driving the observed associations (Supplementary Figure 18).

**Supplementary Figure 16.** QQ plots for a CMH test of synonymous singletons after filtering genes based on A. number of synonymous variant carriers within a gene and B. number of PTV carriers within a gene.

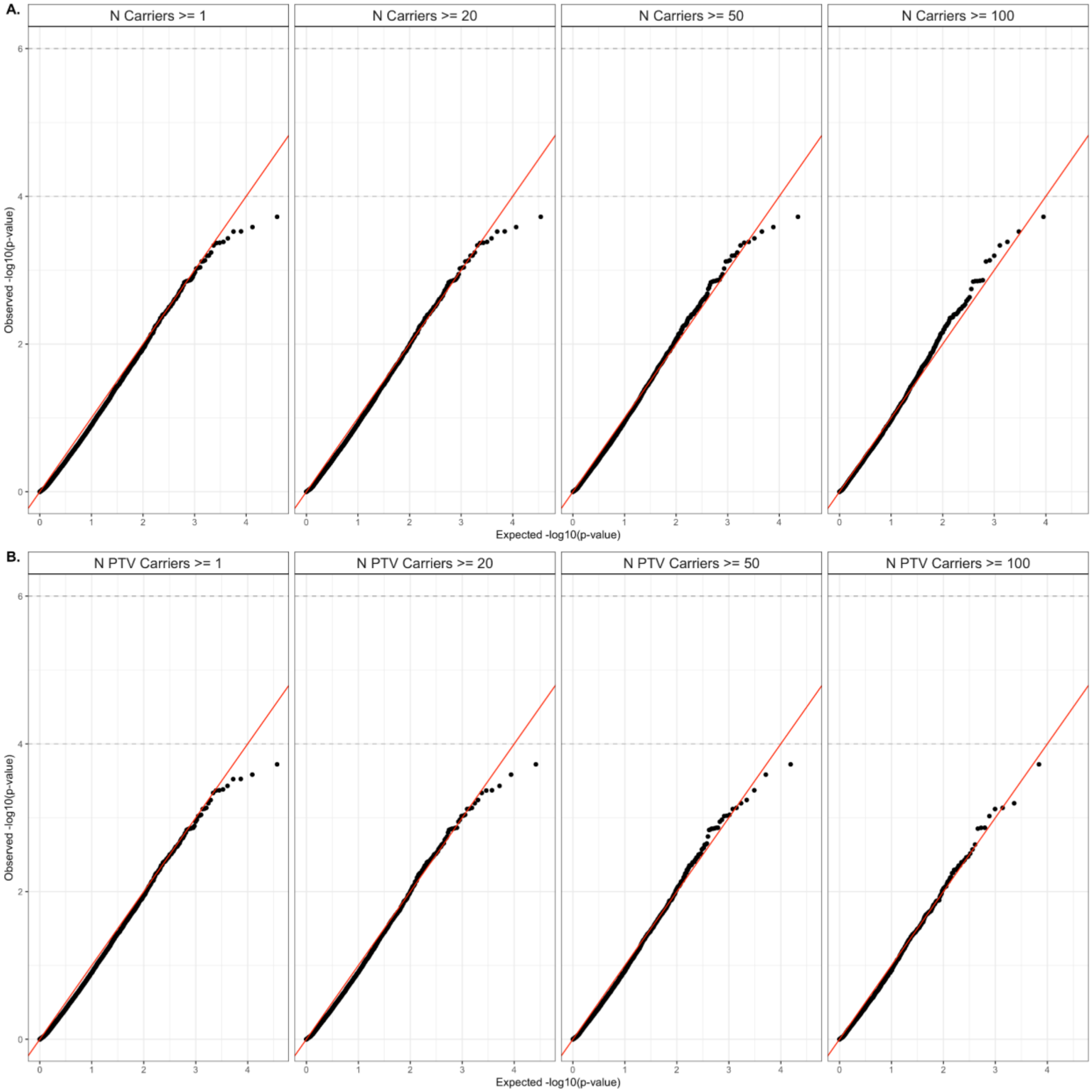

**Supplementary Figure 17.** A. Comparison of p-values from MAC <= 15 (x-axis) and singletons (y-axis). The red line indicates y=x, and the dashed lines indicate FDR 5% significance. Genes considered FDR 5% significant are labelled. B. QQ-plots of MAC <= 15 and singleton pvalues.

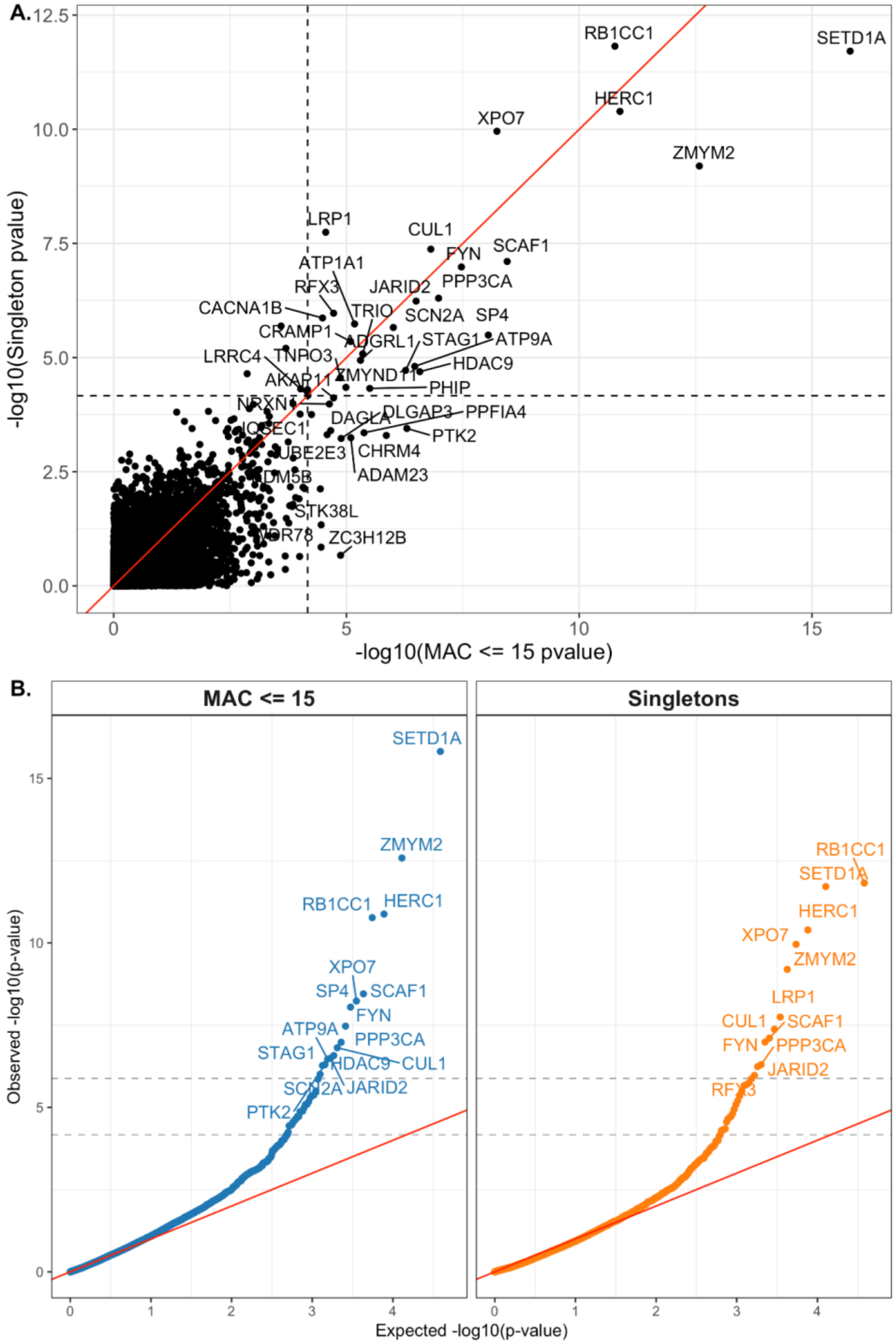

**Supplementary Figure 18.** Case-control RVAS after removing variants with >0.1% population maximum (popmax) allele frequency in RGC or gnomAD v4. The exome wide significant genes remained the same.

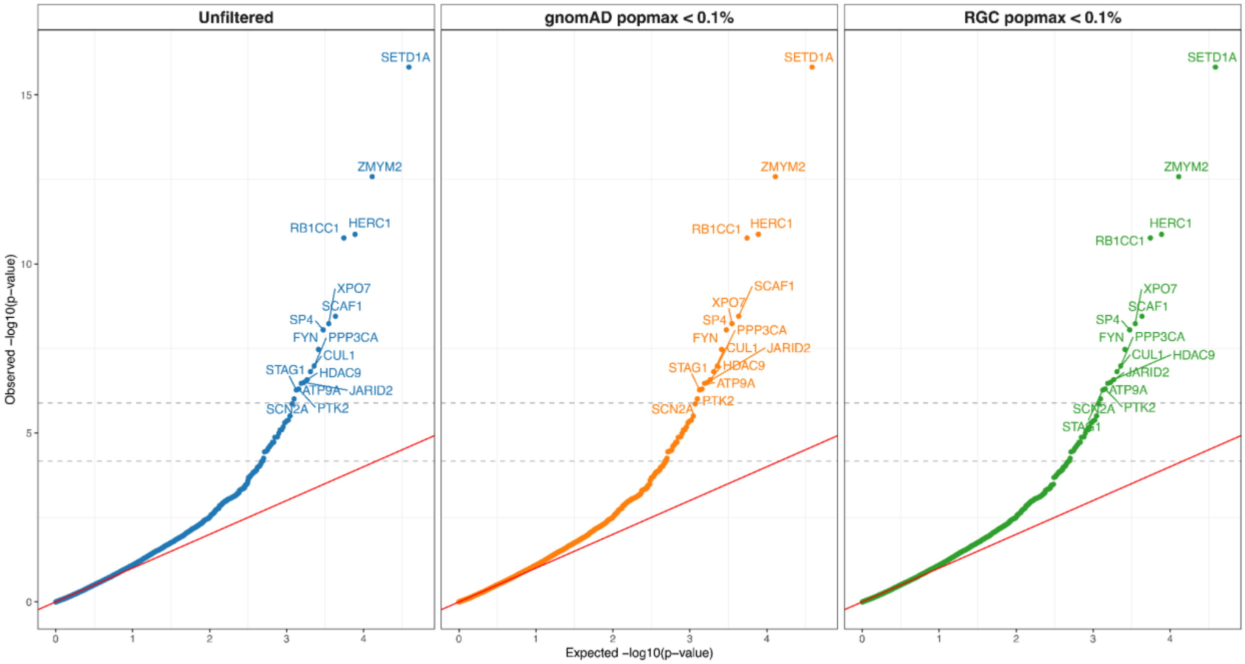

#### Comparing to SCHEMA 1.0 results

**Supplementary Figure 19.** Manhattan plot of SCHEMA 1.0 Bonferroni significant genes in the updated SCHEMA 2.0 results.

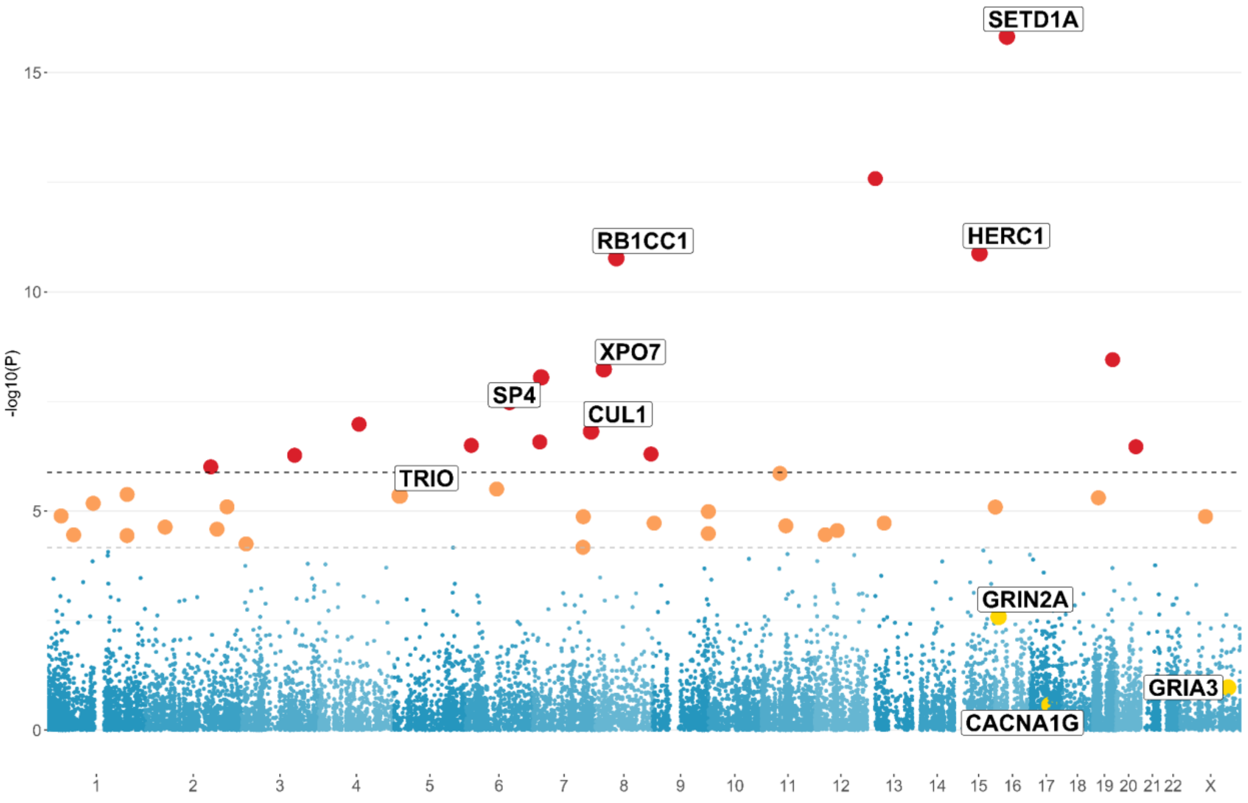

**Comparison of Missense Definitions**

In SCHEMA 1.0, damaging missense variants were defined using the missense badness, PolyPhen-2, and constraint (MPC) score, with variants stratified into Class 2 ($2 < MPC \leq 3$) and Class 1 ($MPC > 3$). Because Class 2 variants showed minimal evidence of enrichment, we focused on Class 1 variants ($MPC > 3$). To assess the impact of missense annotation strategy on gene discovery in SCHEMA 2.0, we repeated the rare variant association study (RVAS) using $MPC > 3$ in place of the $MisRank \geq 93$ threshold. For each gene, we selected the minimum p-value across tests of PTVs alone and the combined test of PTVs and damaging missense variants (PTVs + $MPC > 3$). These results were then compared with those obtained using the $MisRank \geq 93$ approach. Overall, the MisRank-based analysis identified a larger number of associated genes at a false discovery rate (FDR) of 5% (40 genes) compared with the $MPC > 3$ analysis (25 genes), indicating that the MisRank framework captures additional association signal by integrating information across multiple missense annotations rather than relying on a single score (Supplementary Figure 20).

**Supplementary Figure 20.** A. QQ plots of the minimum p-value between PTV | PTV + MisMeanRank ≥ 93 in red and PTV | PTV + MPC > 3 in blue. The dashed line indicates exome-wide significance. B. Comparison of -log10(pvalues) between PTV | PTV + MisMeanRank ≥ 93 (y-axis) and PTV | PTV + MPC > 3 (x-axis). The dashed lines indicate exome-wide significance, and FDR 5% significance. Genes nominated at FDR 5% significance in SCHEMA 2.0 are labelled.

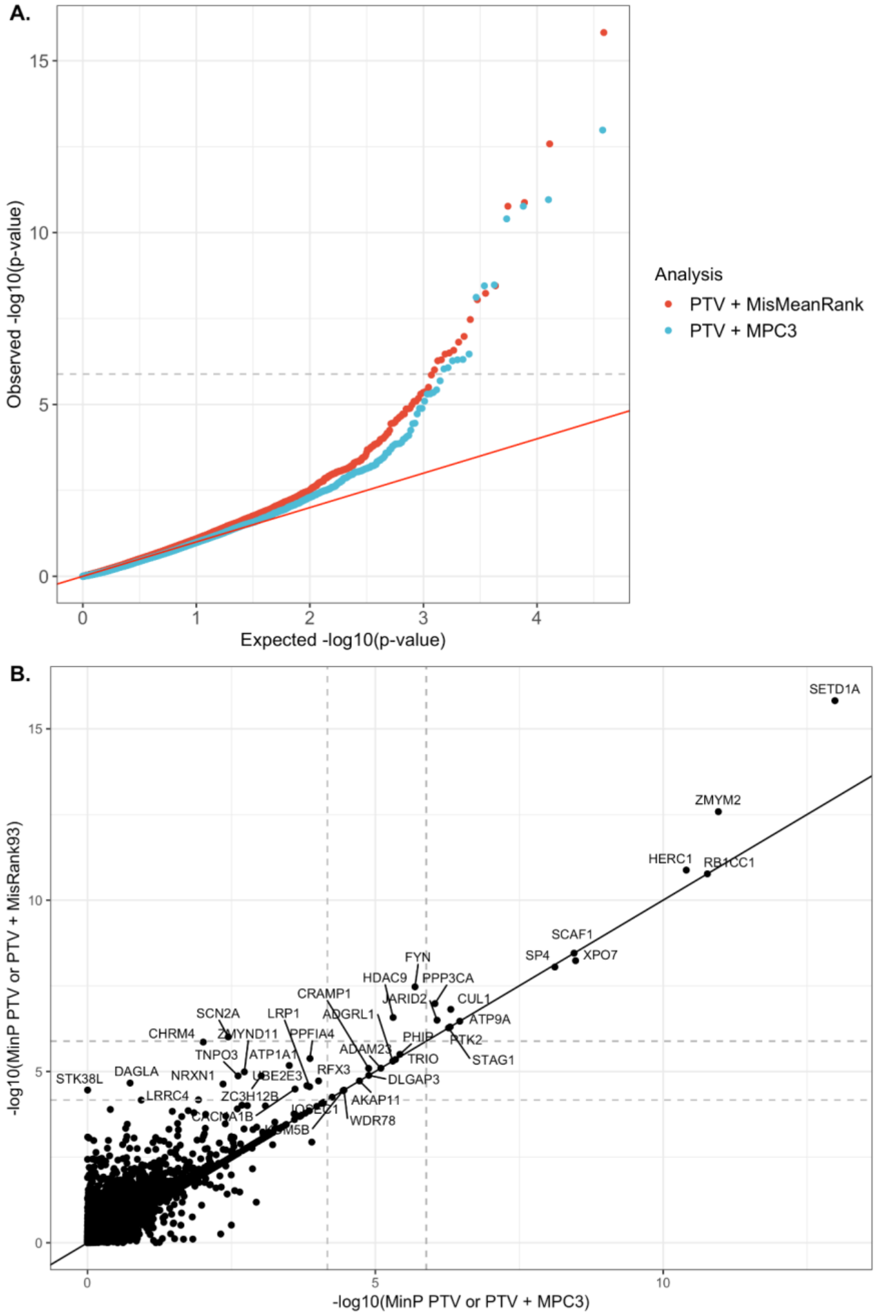

**Supplementary Figure 21.** Forest plot of log-transformed PTV odds ratios for SCHEMA 1.0 exome-wide significant genes compared to SCHEMA 2.0. OR and upper 95% confidence interval estimates were capped at OR = 100. Arrows indicate capped values for SETD1A, CUL1, XPO7, SP4, GRIA3, and GRIN2A.

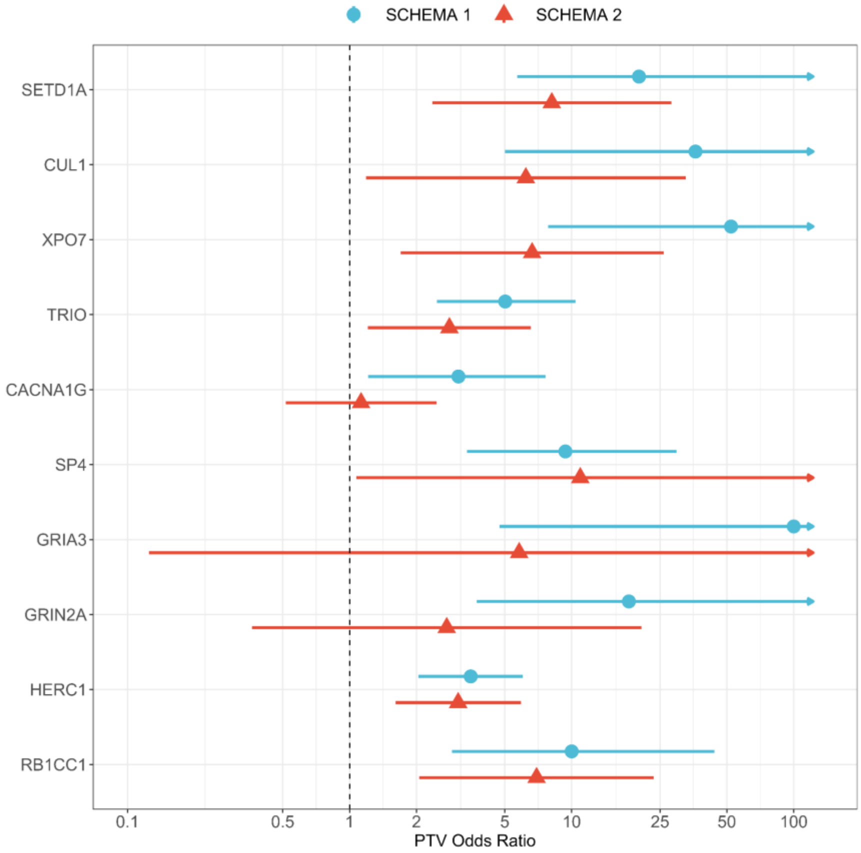

**Enrichment of PTVs in DD/ID Genes by Callset**

Given the lack of replication of three DD/ID related genes from SCHEMA 1.0 to SCHEMA 2.0 (*CACNA1B, GRIA3,* and *GRIN2A),* we sought to determine if datasets new to SCHEMA 2.0 were depleted for DD/ID cases. Because we do not have access to the full medical record, we used genetics as a proxy. We extracted PTV MAC <= 15 in genes implicated in DD/ID from SCHEMA 2.0 cases and controls. Next, we calculated carrier frequencies between cases and controls within each combination of callset and capture kit (Supplementary Figure 20A). We separated out cohorts from SCHEMA 1.0 into the 'gnomAD Exomes (SCHEMA1)' grouping seen in the figure. Next, we used the carrier frequencies to calculate a rate ratio, as the case carrier proportion divided by the control carrier proportion for each grouping (Supplementary Figure 20B).

This analysis demonstrated that the new datasets have decreased case carrier rates and rate ratios for PTV variants in DD/ID genes compared to SCHEMA 1.0 cohorts, signaling a potential recruitment bias that would reduce our ability to identify genes overlapping with DD/ID.

**Supplementary Figure 22.** A. Carrier frequency of PTV MAC <= 15 variants in genes associated with developmental delay or intellectual delay (DD/ID) stratified by cases (solid line) and controls (dashed lines), callset, and capture kit. B. Enrichment of PTV MAC <= 15 within DD/ID genes stratified by callset and capture kit. In both plots, cohorts present in SCHEMA 1.0 are separated from new cohorts in the gnomAD Exomes data.

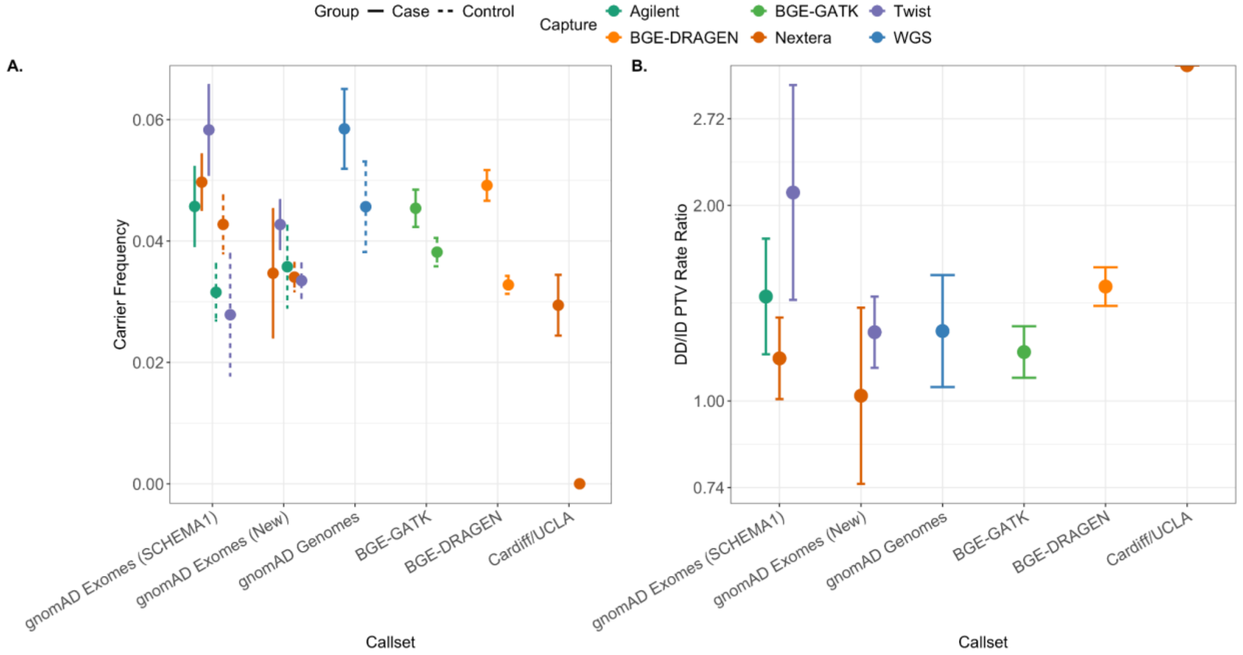

**GWAS Overlap**

To assess the overlap between common variant associations and genes identified through rare variant analysis, we utilized summary statistics from the most recent schizophrenia genome-wide association study (GWAS) conducted by the Psychiatric Genomics Consortium (Trubetskoy et al., 2022). This GWAS employed a multi-stage meta-analytic and fine-mapping approach to identify risk loci. For the most comprehensive set of common variant associations, we used the 287 loci reported in the "extended GWAS," as listed in Supplementary Table 3 of the original study. Because the GWAS summary statistics were based on the GRCh37 human genome reference and our RVAS was conducted using GRCh38, we converted GWAS loci to GRCh38. Gene coordinates were obtained from the RefSeq annotation track via the UCSC Genome Browser.

To define overlap, we identified index SNPs located within ±0.5 megabase of any SCHEMA 2.0 gene reaching an FDR threshold of 5%, giving 6 index SNPs in proximity to 6 SCHEMA 2.0 genes (Table 1, Supplementary Figure 21A). To evaluate whether this observed overlap exceeded random expectation, we conducted a permutation-based analysis. We randomly selected 10 genes from the genome and calculated the number of overlaps with GWAS index SNPs within ±0.5 megabase, repeating this process 100,000 times. The resulting empirical p-value was 0.41 (Supplementary Figure 21B), indicating that the observed overlap was not significantly greater than expected by chance.

**Supplementary Figure 23**. A. Overlap between GWAS index SNPs and SCHEMA 2.0 FDR 5% genes. Orange points represent the linkage disequilibrium block associated with the index SNP. B. Results of simulated overlap of 6 genes with GWAS index SNPs, repeated 10,000 times. C. Enrichment of MAC <= 15 variants by annotation within 64 fine-mapped protein coding genes.

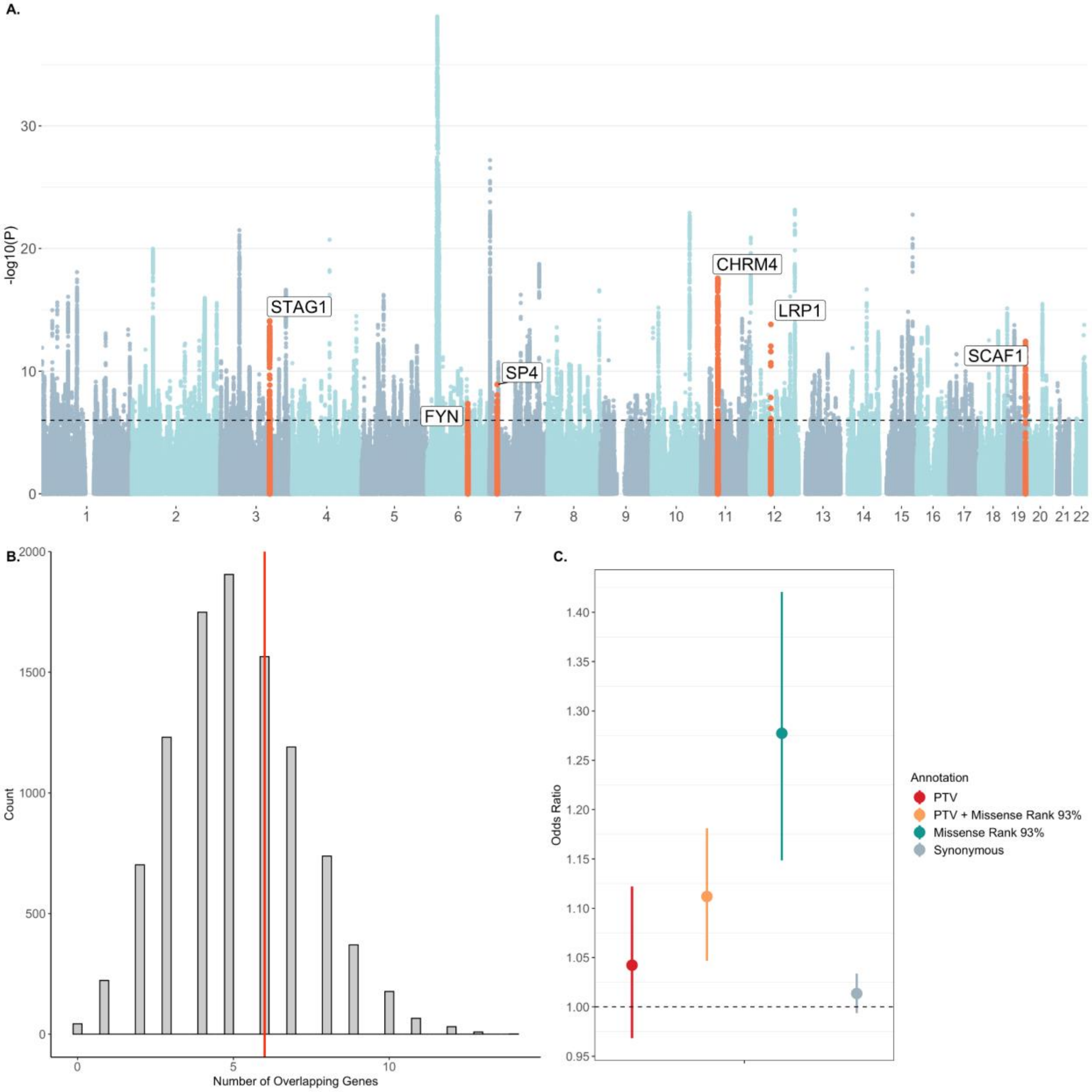

#### Integrating Expression Data

##### PsychEncode

We obtained single-cell gene expression data for the 36 SCHEMA FDR 5% genes from the PsychScreen portal (https://psychscreen.wenglab.org/psychscreen/gene). The dataset includes dorsolateral prefrontal cortex (DLPFC) expression profiles from 294 donors across eight cohorts, encompassing 27 annotated cell types. Expression values were provided both at the level of individual cell types and grouped into broader categories. We first averaged gene expression across cohorts within each broad cell type. For each gene, we then z-score normalized expression values across the broad cell types to enable within-gene comparisons (Supplementary Figure 22A). This process was repeated for the 27 individual cell types to generate gene-specific z-score profiles at higher resolution (Supplementary Figure 22B). To identify patterns of gene co-expression, we focused on the broad cell type groupings and performed k-means clustering with $k = 4$. The choice of $k$ was guided by visual inspection of the within-cluster sum of squares using the elbow method (Supplementary Figure 23). Each gene was assigned to one of the four clusters based on its expression profile, and expression patterns were visualized by cluster assignment (Figure 3B).

**Supplementary Figure 24.** A. Grouped single cell expression of SCHEMA 2.0 genes from PsychEncode across 7 groups. B.Single-cell expression of SCHEMA 2.0 genes from PsychEncode across 27 cell types. Expression is standardized across cell types within each gene.

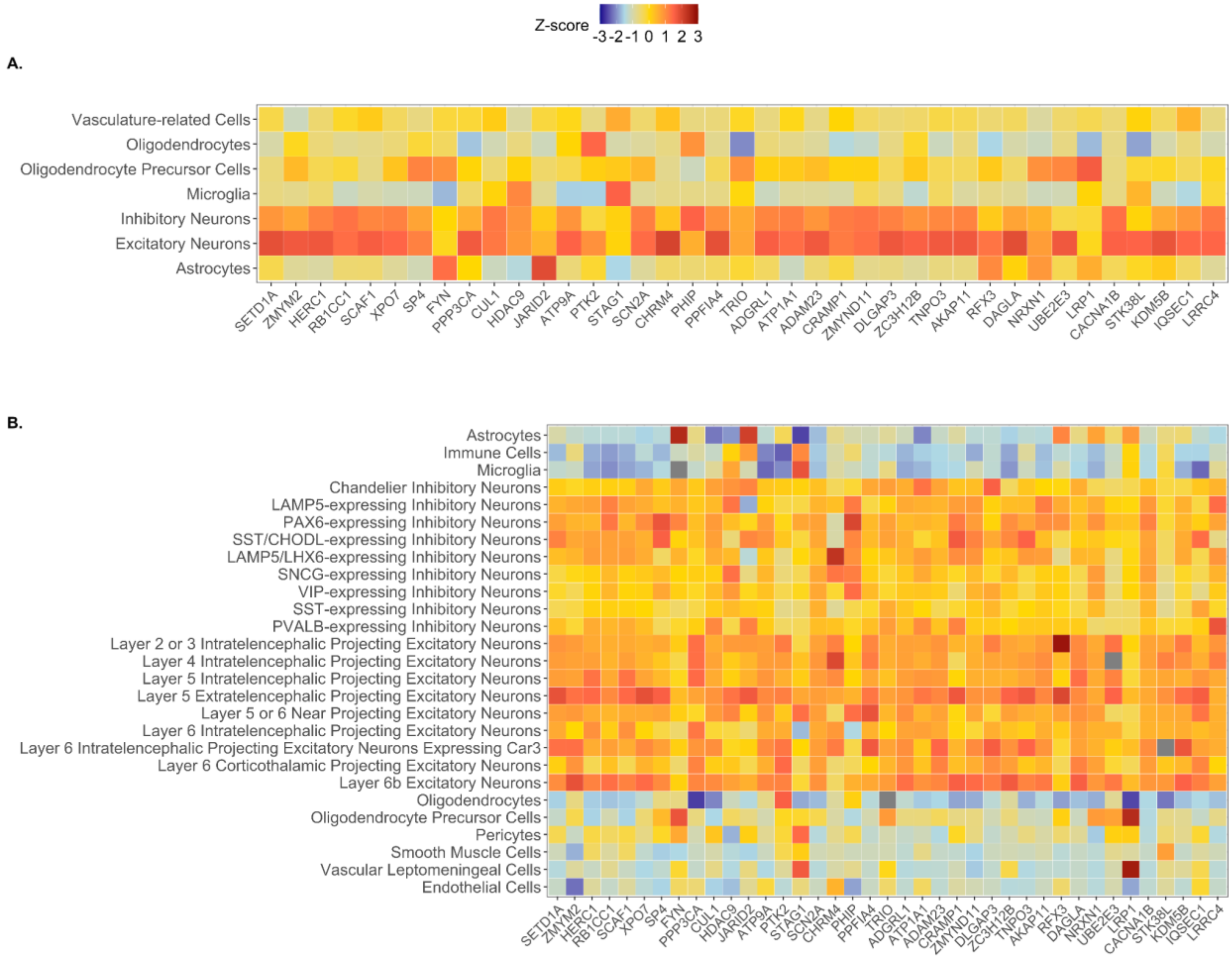

**Supplementary Figure 25.** Elbow method for determining optimal k in the k-means clustering analysis of broad cell type expression.

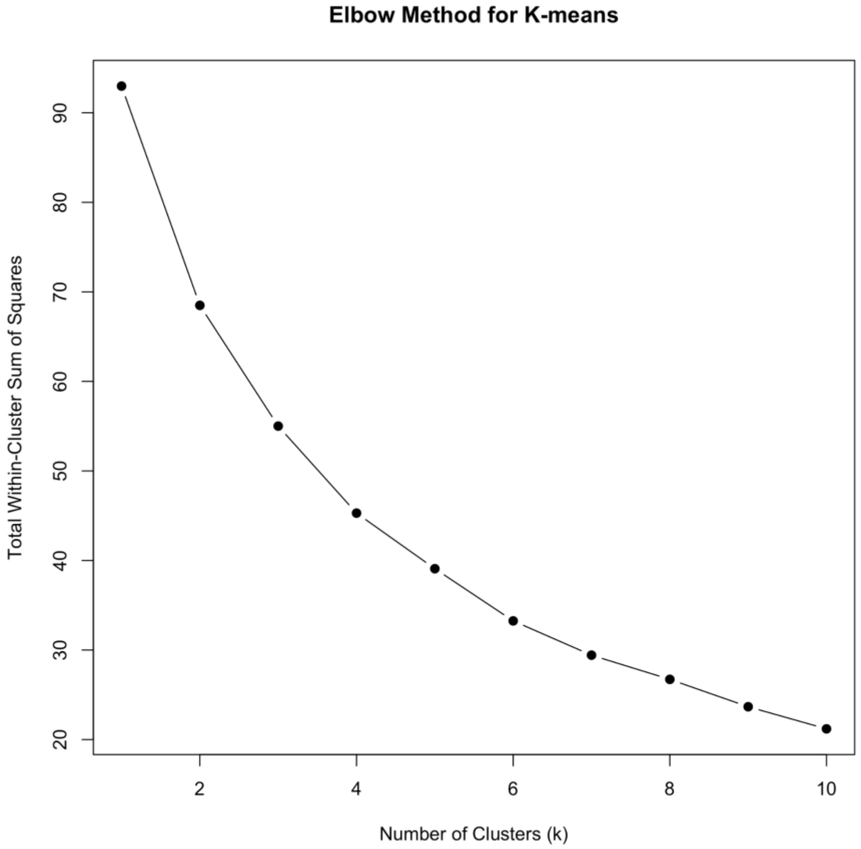

**BrainSpan**

We obtained gene expression data from the BrainSpan Atlas, which includes transcriptomic profiles from 43 human donors across 27 brain structures spanning developmental time points from 8 weeks post-conception to 40 years of age. Developmental stages were grouped into seven age bins: first trimester (<13 weeks post-conception), second trimester (13–26 weeks), third trimester (27–40 weeks), infancy (birth to <1 year), childhood (1–12 years), adolescence (13–20 years), and adulthood (>20 years).

For each SCHEMA 2.0 FDR 5% gene, we z-score scaled transcripts per million (TPM) values across all samples and brain structures within that gene to enable within-gene comparisons of developmental expression patterns. For each gene and developmental stage, we averaged the z-score scaled TPM values across all donors and brain structures to generate a heatmap of developmental expression patterns (Figure 3C).

To visualize absolute expression levels, we first averaged TPM values across all brain structures for each donor to obtain a brain-wide expression estimate. We then plotted the expression across developmental stages for each gene, as shown in Supplementary Figure 24.

**Supplementary Figure 26.** Average brain-wide expression values binned by age group for each SCHEMA 2.0 FDR 5% gene.

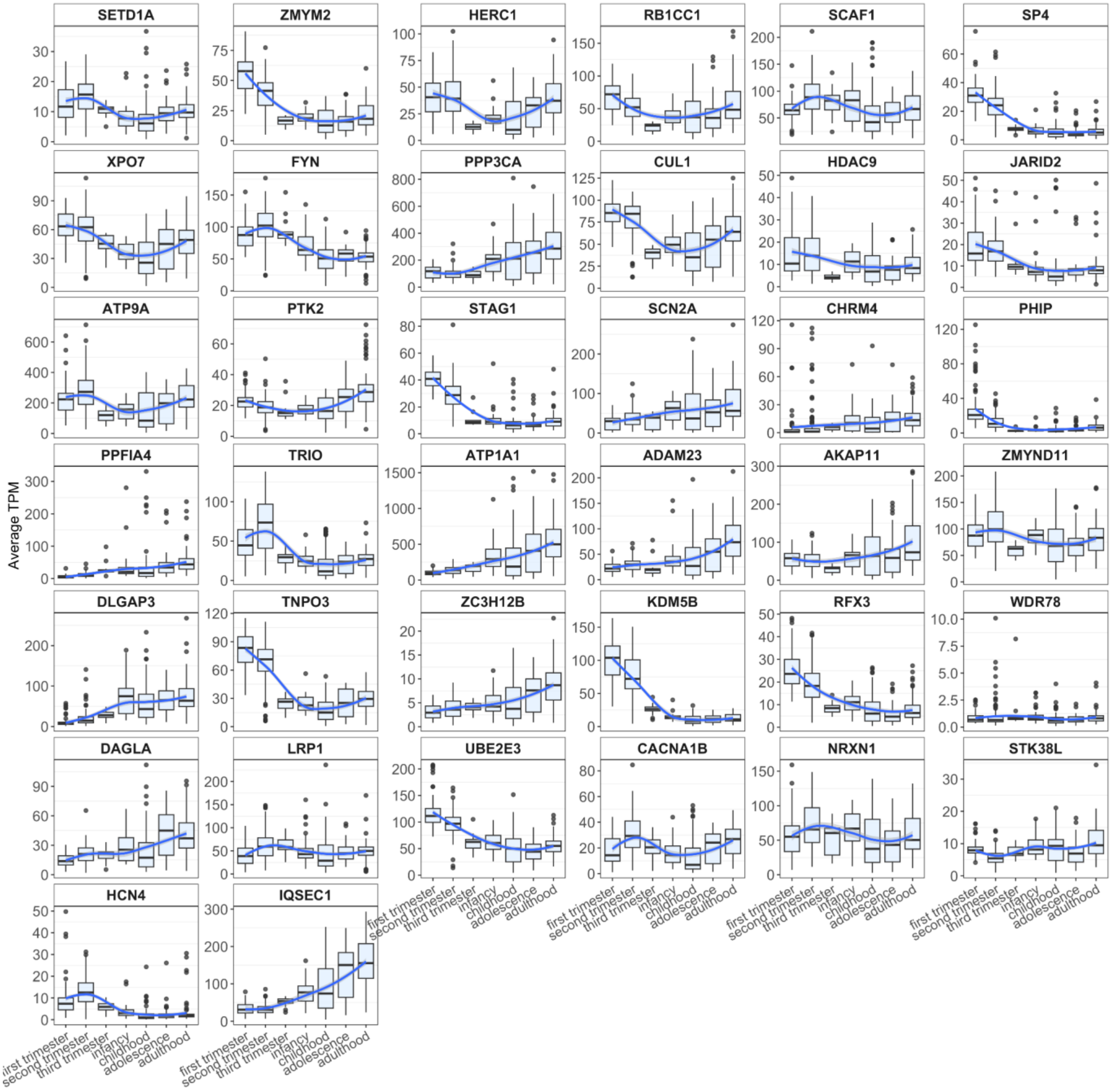

#### Gene Ontology

We used the web tool g:Profiler to conduct functional enrichment analysis of the SCHEMA 2.0 genes. g:Profiler integrates multiple databases, including Gene Ontology, KEGG, and TRANSFAC to provide a comprehensive evaluation of gene lists. First, we analyzed the 36 FDR 5% genes (plotted as red dots in Supplementary Figure 21) for biological processes and cellular components. Across all genes, we identified enrichment in ion transport processes and synaptic components. Next, we repeated the analysis stratifying to each cluster identified using PsychEncode expression data (Figure 3B).

**Supplementary Figure 27.** Gene ontology of A. Biological Processes and B. Cellular Components from g:Profiler. The red dots indicate the results from the 40 SCHEMA 2.0 FDR 5% genes, and the blue, green, purple, and orange dots represent the ontology results stratified by expression clusters. The size of the dot is proportional to the number of genes.

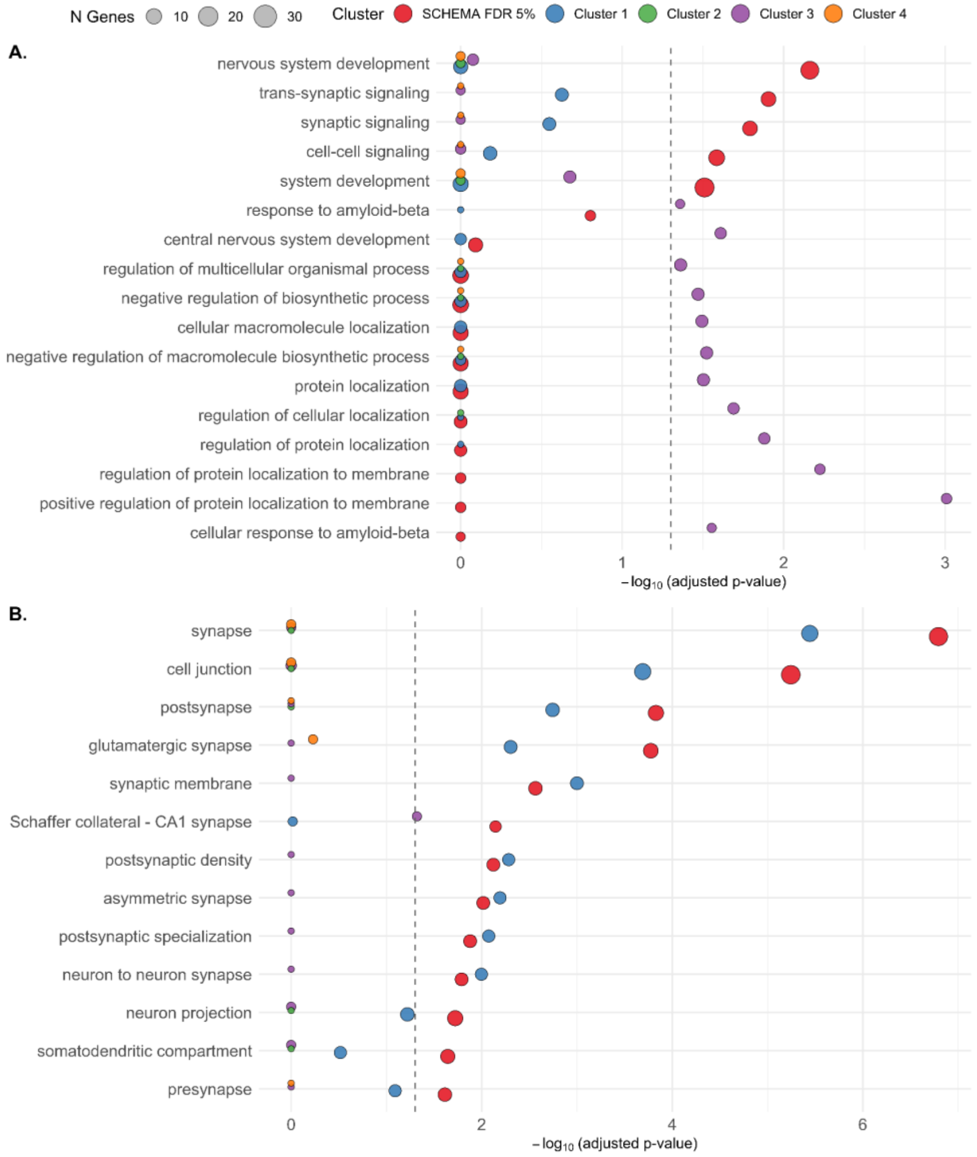

#### PheWAS of Carrier Status in the All of Us Research Program

We utilized data from the All of Us Research Program Curated Data Repository (CDR) version 8(ref:[19]), which includes electronic health records (EHRs) and whole genome sequencing (WGS) data from hundreds of thousands of participants. We analyzed a subset of 307,694 participants with available EHR data, short-read WGS, inferred genetic ancestry, and reported male or female sex at birth. Phenotypes were defined using the PheWAS R package, mapping ICD-9/10 codes to phecodes, with cases requiring two or more code instances; phecodes with fewer than 100 cases or controls were excluded. Carrier status for deleterious variants in 40 SCHEMA 2.0 schizophrenia-associated genes was determined using Hail, based on rare protein-truncating variants (PTVs), annotated with Ensembl VEP and gnomAD v4.1 data. A total of 1,581 PTV carriers met inclusion criteria. Logistic regression was used to assess associations between carrier status and phecodes, adjusting for age, sex at birth, and the first 10 genetic principal components Supplementary Figure 26A. In a sensitivity analysis, we added an additional covariate for lifetime schizophrenia diagnosis using phecode 295.1 (Supplementary Figure 26B).

**Supplementary Figure 28.** PheWAS of A. PTV carriers controlling for age, sex, and top 10 principal components, and B. controlling for lifetime schizophrenia diagnosis in the All of Us Research Program. The red line indicates Bonferroni correction (p = 2.82e-5).

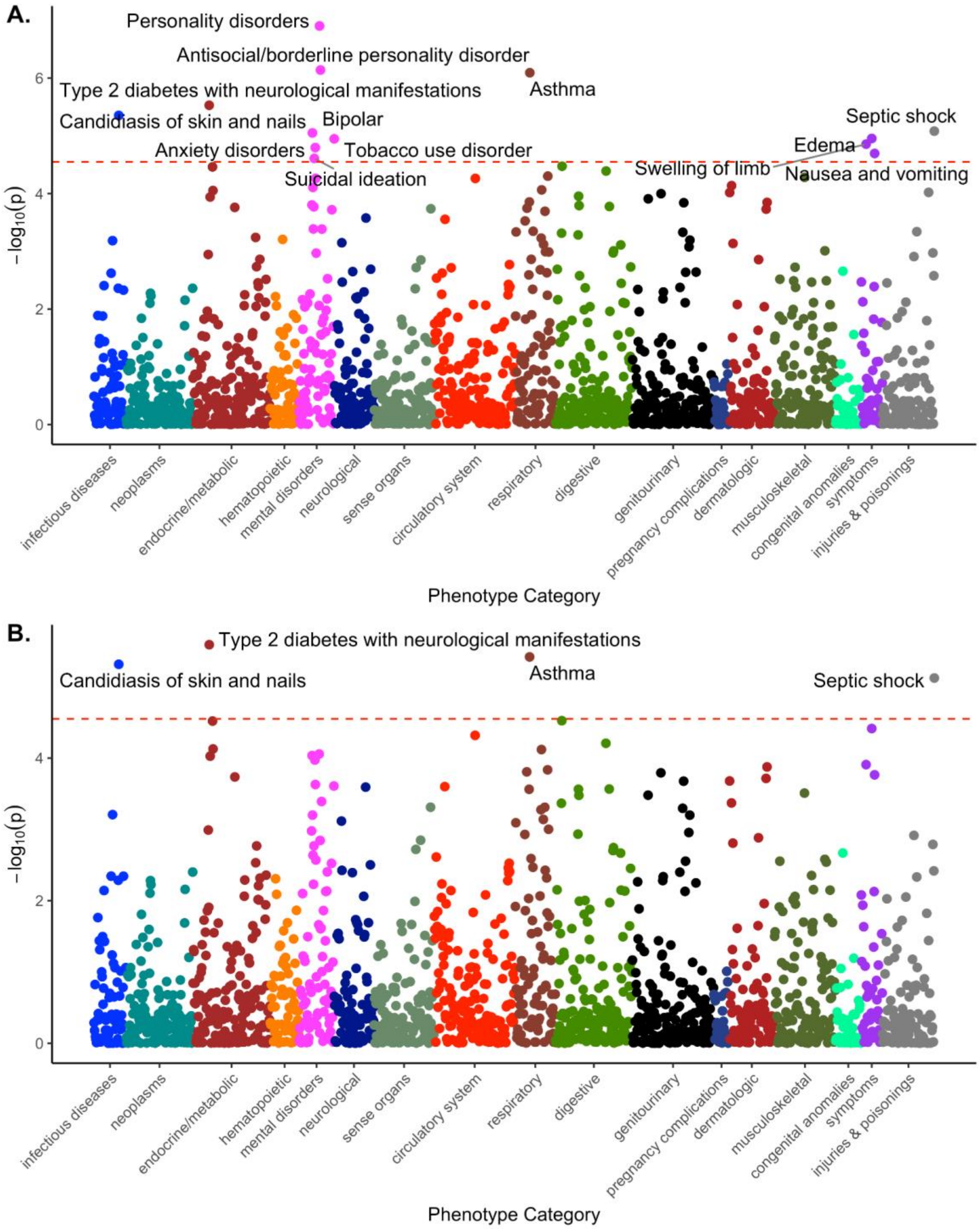

**Full SCHEMA Consortium Authorship, Alphabetical**

Adeel[1], Ahmer[2], Ambreen[3], Irfan[4], Lutfullah[5], Nasir[3], Sheeba[1], Tayyaba[6], Zahoor[1], Nathalie Acevedo[7], Benjamin O Adegoke[8], Aderopo Adelola[9], Elizabeth O Aderinmola[10], Ismail O Adesina[10], Hassan Adeyemi[9], Rolf Adolfsson[11], Adil Afridi[12], Jalil Afridi[13], Bashir Ahmad[14], Hussain Ahmad[15], Shams Uddin Ahmad[16], Zaheer Ahmad[17], Noman Ahmed Gill[5], Javed Akhtar[18], Syed Mubin Akhtar[19], Kazufumi Akiyama[20], Aftab Alam[21], Melkam Alemayehu[22], Gohar Ali[23], Hazrat Ali[24], Jawad Ali[25], Mian Nizam Ali[26], Muhammad Ali[1], Simon G Anderson[27], Moin Ahmed Ansari[1], Aqsa Anwar[6], Sibtain Anwar[4], Makoto Arai[28], Naohiro Arai[29,30], Tetsuaki Arai[31], Celso Arango López[32,33], Alejandro Arias[34], Robert Asarnow[35], Lukoye Atwoli[36,37], Summaira Awan[1], Nora Ayola Serrano[38], Ayoyinka Ayorinde[9], Muhammad Ayub[39], Mian Mukhtar Ul Haq Azeemi[12], Xuefei Bai[40], Nasir Baig[6], Malek Bajbouj[41], Saqib Bajwa[5], Kathleen C Barnes[42], Nick Bass[43], Zahida Batool[1], Carrie E Bearden[44,45], Sintia I Belangero[46,47], Kiramat Ullah Bettani[15], Moti Ram Bhatia[48], Tim B Bigdeli[49], Douglas H Blackwood[50], Marco P Boks[51], Shuken Boku[52], Toni Boltz[53,54], Harrison Brand[55,56,57,58], Alice Braun[59,41], Gerome Breen[60], Hung Luu Bui[61], Shogyoku Bun[62], Ammara Butt[3], Jonas Bybjerg-Grauholm[63,64,] Anders D Børglum[65,66,67], Lucas C. Quarantini[68], Camarena Beatriz[69], Monica Campbell[42], Truong Xuan Cao[70], Zhongyu Cao[40], Luis Caraballo[7], Mauricio Castaño[71,72], Saulo G Castor Albuquerque[73], Felecia Cerrato[74], Chiao-Erh Chang[75,76], Parveen Channar[1], Katherine Chao[77,78,54], Sinéad B Chapman[53,54,79], Hsi-Chung Chen[80,81,82], Luan Chen[40,83], Ruoyu Chen[40], Ahsan Ul Haq Chishti[84], Abdul Rashid Choudhary[85], Shahzad Tahir Choudhary[86], Yunpeng Chu[40], Claire Churchhouse[87,74], Aiden P Corvin[88], Nicholas Craddock[89], Nicolas Crossley[90], Álvaro Augusto Souza da Cruz Filho[91], David Curtis[92], Caroline Cusick[74], Mobolaji U Dada[8], Shahnawaz Dal[1], Mark J Daly93,54,78,77, Candace Machado de Andrade94, Juan F De la Hoz44,95,96, Andre L de Souza Rodrigues[97,98], Ana M Diaz-Zuluaga[44], Akena Dickens[99], Allah Din[1], Mateus Diniz[100], Quang Hong Doan[101], Sheila Dodge[102], Farasat Ali Dogar[3], Imtiaz Ahmad Dogar[6], Connor Dowd[87,74], Huihui Du[40], Olaniyi O Duduyemi[103], Onyedikachi Eguzoro[104], Eija Hämäläinen[93], Oladoyin Esan[9], Michael A Escamilla[105], Javier I Escobar[106], Tõnu Esko[107], Oluwatoyin A Fasesan[104], Xiaoqi Feng[40], CAMILA ALEXANDRINA VIANA DE FIGUEIREDO FONTANA[94], Jennifer Forsyth[108], Nelson Freimer[44], Thiago H Freitas[109,110], Clara Frydman-Gani[44], Jack Fu[111], Kumiko Fujii[112], Stacey B Gabriel[102], Ary Gadelha[113,72], Greta Gerdes[44], Stella Gichuru[114], Shamshad Ahmed Gill[115], Stephen J Glatt[116], Juliana Gomez-Makhinson[117], Riley Grant[118], Michael F Green[35], Jakob Grove[65,66,67], Zhenglin Guo[74], Eric Hahn[119], Arsalan Hassan[87,74], Kotaro Hattori[120], Lin He[40], Teruhiko Higuchi[121], Akitoyo Hishimoto[122], Dung Ho[123], Nam Giang Ho[124], Yasue Horiuchi[125], Daniel Howrigan[87,74], Cong Huai[40], Hailiang Huang[54,126], Ming-Chyi Huang[127,128], Christina Hultman[129], Ashfaq Hussain[130], Asif Hussain[13], Fahad Hussain[131], Mian Iftikhar Hussain[132], Neilofar Hussain[1], Olanrewaju I Ibigbami[9], Masayuki Ide[31], Muhammad Idrees[133], Osemhen Ighedosa[103], Parveen Ijaz[1], Masashi Ikeda[134], Muhammad Ilyas[24], Sadia Iqbal[17], Hiroki Ishiguro[135], Sayuri Ishiwata[120], Masanari Itokawa[28], Nakao Iwata[136], Shaili Jha[137,74], Bixuan Jiang[40], Joanna Jiménez-Pavón[138], Deborah Jonker[139], Kyebuzibwa Joseph[99], Temitope A Joseph[103], Jukka T. Koskela[93], Jamil Junejo[1], René S Kahn[140], Shigenobu Kanba[141], Konrad J Karczewski[87,118], Symon M Kariuki[142], Takahiro A Kato[143], Kenneth S Kendler[144], Tanveer Khalid[145], Ahmad Khan[146], Anis Khan[146], Fahadullah Khan[131], Inzemam khan[131], Kamran Khan[147], Muhammad Firaz khan[148], Muslim Khan[149], Bakht khizar[131], Hassan T Kilani[103], Soyeon Kim[150], Kimmo Kontula[151], Makoto Kinoshita[152], George Kirov[89], Toshifumi Kishimoto[153], James Knowles[154], Karestan C Koenen[137,74,155,156], Abdulghaffar O Kolade[103], Alex Kopelowicz[117], Julia Kraft[59,41], Hiroshi Kunugi[157], Po-Hsiu Kuo[158], Itaru Kushima[159], Chooni Lal[160], Minh Tuong Lam[161], Mikael Landen[129], Abisola Lawal[10], Linh Thuy Le[162], Manh Tien Le[163], Ngoc Minh Le[164], Tan Bat Le[165], Calwing Liao[87,118,74], Yunxiao Lin[40], Penelope A Lind[166], Qinyi Liu[40], Esteban A Lopera-Maya[44], Carlos Lopez-Jaramillo[167,90], Pedro G Lorencetti[113,72], Regina Mahmood[5], Dana Manoach[168], Niaz Maqsood[169], Alicia R Martin[170,171], Martti Farkkila[172], Odunayo K Mayowa[104], Andrew McIntosh[50], Andrew McQuillin[173], Sarah Medland[166], Muntazir Mehdi[174], Khalid Mahmood[175], Nasir Mehmood[2], Zainab Mehmood[3], Raza Memon[1], Mikko Hiltunen[176], Masaru Mimura[62], Jun Miyata[177], Pamela Morales-Cedillo[69], Derek W Morris[178], Ole Mors[179,66], Preben Bo Mortensen[180], Ali Ahsan Mufti[181], Khalid Mufti[181], Shah Muhammad[4], ToshiyaMurai[182], Ali Burhan Mustafa[183], Rehema Mwende[184], Nazish Naqvi[1], Zahid Nazr[12], Benjamin Neale[87,118,74,171], Charles Newton[185], Chuong Anh Nguyen[186], Hung Van Nguyen[187], Huu Tu Nguyen[188], Khoa Dang Nguyen[189], Nam Hoai Nguyen[190], Phat Manh Nguyen[191], Phuong Doan Nguyen[192], Thang Huu Nguyen[193], Tinh Van Nguyen[194], Tuan Van Nguyen[195], Van Phi Nguyen[196,197], Hideto Niimura[62], Asad Tamizuddin Nizami[198], Keith Nuechterlein[35], Shusuke Numata[152], Joshua E Nwokike[103],

Michael C O'Donovan[89], Adetunji Obadeji[8], Olakunle Oginni[199,200], Omotola Ogunjobi[103], Tetsuro Ohmori[201], Hideyuki Okano[202], Adeniran Okewole[203], Maryam G Okorejior[8], Adesola Olalekan[204], Loes Olde Loohuis[44,205,206], Ana Maria Olivares[74], Nayana Holanda de Oliveira[207], Dost Ongur[208], Nnena C Onuorah[10], Roel A Ophoff[209], Tolulope A Oso[103], Ikuo Otsuka[122], Willem Ouwehand[210], Andrew M Owadokun[10], Michael J Owen[89], Norio Ozaki[159], Yuji Ozeki[112], Juan David Palacio-Ortiz[90,167], Aarno Palotie[93,54,211,212,78,77], Carlos N Pato[213,171], Michele T Pato[213,171], Nancy Pedersen[214], Kieu Duy Pham[215], Gabriela Pimentel Pinheiro das Chagas[91], Olayiwola A Popoola[204], Danielle Posthuma[216], Ann E Pulver[217], Syed Qalb-i-Hyder[1], Shengying Qin[40], Fazal Rabbani[12], Ayanjide L Raji[10], Aatir Rajput[1], Ana M Ramirez-Diaz[44], Muhammad Azeem Rao[5], Qazi Humayun Rashid[1], Saleem Rashid[218], Ghulam Rasool[219], Huma Rasool[1], Mohsin Raza[1], Elliott Rees[89], Andreas Reif[220], Victor I Reus[221,222], Qasim Riaz[13], Stephan Ripke[170,41], Angela Rose[223], Päivi Saavalainen[224], Chiara Sabatti[225], Marco Antonio Sanabrais-Jiménez[69], Cinthia Vila Nova Santana[226], Marcos L Santoro[227,72], Akira Sawa[228], Julia Sealock[87,74], Susan Service[44,229], Sadia Shafiq[13], Abdul Shakoor[230], Shabbir Sheikh[1], Lu Shen[40], Kazutaka Shimoda[231], Hatilla dos Santos Silva[94], Tarjinder Singh[232], Jordan W Smoller[233,234], Matthew Solomonson[118], Ichiro Sora[122], Carlo Esteban Sotelo-Ramírez[69], Oladipo A Sowunmi[103], Dan J Stein (In memoriam)[235], Christine R Stevens[77,78,54], Rocky Stroud II[137,74], Patrick F Sullivan[129,236], Sayed Muhammad Sultan[237], Jing Sun[40], Yidan Sun[40], Michio Suzuki[238], Thi Minh Tam Ta[119], Momina Tahir[6], Rizwan Taj[239], Atsushi Takata[240], Yoichiro Takayanagi[238], Minoru Takebayashi[241], Michael E Talkowski[242,57,58,56,243], Jie Tan[40], Bansi Lal Tanwani[1], Muhammad Tariq[13], Solomon Teferra[22], Terri Teshiba[44], Zizhao Tian[40,244], Nicholas Timpson[245], Michihiro Toritsuka[246], Tomoko Toyota[121], Ngoc Xuan Tran[247], Ngoc Nguyen Tran[248], Nhan Ngoc Tran[249], Quang Trong Tran[250], Vinh Van Tran[251], Tuan Xuan Trinh[252], Ming Tsuang (In memoriam)[253], Muhammad Umar[39], Raza ur Rehman[254], Johanna Valencia-Echeverry[90], Marquis P Vawter[255], Biju Viswanath[256], Sinh Canh Vo[257], Annabel Vreeker[258], Hanh Minh Vu[259], Uy Ngoc Vu[260], James Walters[261], Cong Wang[40], Yongzhi Wang[40], Yuqi Wei[40], Thomas M Werge[262,263], Michael Wilson[118], Chenyu Wu[40], Hao Wu[40], Qiange Xiao[40], Kun Yang[264], Jiajie Ye[40], Robert Ye[74], Robert Yolken[265], Yujiro Yoshihara[182], Takeo Yoshikawa[121], Khwaja Younas[266], Adnan Yousaf[5], Muhammad Yousuf[1], Peter P Zandi[267], Na Zhang[40], Yingtian Zhang[40], Xianglong Zhao[40], Chenxi Zhou[40], Wei Zhou[40,83], Jinhang Zhu[40], Carolina Ziebold[113], Zukiswa Zingela[268], Ali Zulqarnain[269].

1. Sir Cowasjee Jehangir Institute of Psychiatric and Behavioral Sciences, Hyderabad, Pakistan
2. Karwan e Hayat, Karachi, Pakistan
3. Sir Ganga Ram Hospital, Lahore, Pakistan
4. Sibtain Anwar Psychiatric Hospital, Mardan, Pakistan
5. Punjab Institute of Mental Health, Lahore, Pakistan
6. District Headquarter Hospital, Faisalabad, Pakistan
7. Institute for Immunological Research, University of Cartagena, Cartagena de Indias, Colombia
8. Ekiti State University Teaching Hospital, Ado Ekiti, Ekiti State, Nigeria
9. Obafemi Awolowo University Teaching Hospital, Ile-Ife, Osun State, Nigeria
10. Federal Neuropsychiatric Hospital Yaba, Lagos State, Nigeria
11. Department of Clinical Sciences, Psychiatry, Umea University, Sweden
12. Lady Reading Hospital, Peshawar, Pakistan
13. Shafique Psychiatric Hospital, Peshawar, Pakistan
14. Bashir Psychiatric Hospital, Peshawar, Pakistan
15. Mufti Mehmood Memorial Teaching Hospital, Dera Ismail Khan, Pakistan
16. Al-baari Clinic Sargodha
17. Allama Iqbal Teaching Hospital, Dera Ghazi Khan, Pakistan
18. Dr. Javed Akhtar Psychiatric Clinic, Bannu, Pakistan
19. Karachi Psychiatric Hospital, Karachi, Pakistan
20. Department of Biological Psychiatry and Neuroscience, Dokkyo Medical University, School of Medicine, Japan
21. Ayub Medical Complex, Abbottabad, Pakistan
22. Addis Ababa Univeriesty, Ethiopia
23. Saidu Teaching Hospital, Swat, Pakistan

24. Balochistan Institute of Psychiatry and Behavioral Sciences, Quetta, Pakistan
25. Government Mental Hospital, Mansehra, Pakistan
26. Swat Institute of Medical Sciences, Swat, Pakistan
27. Caribbean Institute of Health Research, The University of the West Indies
28. Department of Psychiatry and Behavioral Sciences, Tokyo Metropolitan Institute of Medical Science, Japan
29. Department of Neuropsychiatry, Keio University School of Medicine, Tokyo, Japan
30. Centre for Addiction and Mental Health, Toronto, Ontario, Canada
31. Department of Psychiatry, Division of Clinical Medicine, Faculty of Medicine, University of Tsukuba, Japan
32. Department of Child and Adolescent Psychiatry, Institute of Psychiatry and Mental Health, Hospital
33. General Universitario Gregorio Marañón (IiSGM), Madrid, Spain
34. Alejandro AriasAlejandro
35. University of California, Los Angeles
36. Department of Medicine, the Aga Khan University Medical College East Africa, Kenya
37. Brain and Mind Institute, the Aga Khan University
38. Clínica Psiquiátrica CEMIC, Cartagena, Colombia
39. Division of Psychiatry, University College London, London
40. Bio-X Institutes, Key Laboratory for the Genetics of Developmental and Neuropsychiatric Disorders (Ministry of Education), Shanghai Jiao Tong University, Shanghai, China
41. Charité Universitätsmedizin
42. Galatea Bio, Inc., Miami, FL USA
43. Division of Psychiatry, University College London, London, UK
44. Center for Neurobehavioral Genetics, Semel Institute for Neuroscience and Human Behavior, David Geffen School of Medicine, University of California Los Angeles, Los Angeles, California, USA
45. Department of Psychology, University of California Los Angeles, Los Angeles, California, USA
46. Psychiatry Department, Universidade Federal de São Paulo, São Paulo, SP, Brazil
47. Graduate Program of Functional and Structural Biology, Universidade Federal de São Paulo, São Paulo, SP, Brazil
48. Peoples Medical College Hospital, Nawabshah, Pakistan
49. SUNY Downstate Health Sciences University
50. University of Edinburgh
51. Department of Psychiatry, Amsterdam University Medical Center, University of Amsterdam, Amsterdam, Netherlands
52. Department of Psychiatry, Niigata University, Japan
53. Stanley Center for Psychiatric Research, Broad Institute of MIT and Harvard, Cambridge, MA, USA
54. Analytic and Translational Genetics Unit, Department of Medicine, Massachusetts General Hospital, Boston, MA, USA
55. Pediatric Surgical Research Laboratories, Department of Surgery, Massachusetts General Hospital, Boston, MA 02114, USA
56. Center for Genomic Medicine, Massachusetts General Hospital, Boston, MA 02114, USA
57. Program in Medical and Population Genetics, Broad Institute of MIT and Harvard, Cambridge, MA 02142, USA
58. Department of Neurology, Massachusetts General Hospital and Harvard Medical School, Boston, MA 02114, USA
59. Department of Psychiatry and Psychotherapy, Charité – Universitätsmedizin Berlin, Charitéplatz 1, 10117 Berlin, Germany
60. Social Genetic and Developmental Psychiatry, Institute of Psychiatry, Psychology and Neuroscience, King's College London, SE5 8AF, UK
61. Mai Huong Day Psychiatric Hospital
62. Department of Neuropsychiatry, Keio University School of Medicine, Japan
63. Center for Neonatal Screening, Department for Congenital Disorders, Statens Serum Institut, Copenhagen, Denmark

64. The Lundbeck Foundation Initiative for Integrative Psychiatric Research, iPSYCH, Copenha, Denmark
65. Department of Biomedicine, Aarhus University, Aarhus, Denmark
66. The Lundbeck Foundation Initiative for Integrative Psychiatric Research, iPSYCH, Aarhus, Denmark
67. Center for Genomics and Personalized Medicine, Aarhus, Denmark
68. Department of Neurology and Psychiatry, Faculdade de Medicina da Bahia, Universidade Federal da Bahia
69. Instituto Nacional de Psiquiatria Ramon de la Fuente Muñiz, Mexico city, Mexico
70. My Duc Psychiatric Hospital
71. Department of Mental Health and Human Behavior, Universidad de Caldas, Colombia
72. Interdisciplinary Laboratory of Clinical Neurosciences (LINC), Department of Psychiatry, Paulista School of Medicine, Federal Univetsity of São Paulo
73. Hospital Mental Frota Pinto, Brazil
74. Stanley Center for Psychiatric Research, The Broad Institute of MIT and Harvard, Cambridge, Massachusetts, USA
75. Institute of Epidemiology and Preventive Medicine, National Taiwan University, Taipei, Taiwan
76. Stanley Center for Psychiatric Research, Broad Institute, Cambridge, MA, USA
77. Program in Medical and Population Genetics, The The Broad Institute of MIT and Harvard, Cambridge, MA, USA
78. Stanley Center for Psychiatric Research, The The Broad Institute of MIT and Harvard, Cambridge, MA, USA
79. Program in Medical and Population Genetics, Broad Institute of MIT and Harvard, Cambridge, MA, USA, BD2: Breakthrough Discoveries for thriving with Bipolar Disorder, Santa Monica, CA, USA
80. School of Medicine, National Taiwan University College of Medicine, Taipei, Taiwan
81. Department of Psychiatry, National Taiwan University Hospital, Taipei, Taiwan
82. Center of Sleep Disorders, National Taiwan University Hospital, Taipei, Taiwan
83. Ministry of Education - Shanghai Key Laboratory of Children's Environmental Health & Department of Developmental and Behavioural Paediatric & Child Primary Care, Xinhua Hospital Affiliated to Shanghai Jiao Tong University School of Medicine, 1665 Kongjiang Road, Shanghai 200092, China
84. Recovery Rehab Centre, Lahore, Pakistan
85. New Millat Brain Center, Sahiwal, Pakistan
86. Shahzad Tahir Choudhary
87. Analytic and Translational Genetics Unit, Department of Medicine, Massachusetts General Hospital, Boston, Massachusetts, USA
88. Trinity College Dublin, Dublin, Ireland
89. Cardiff University
90. Research Group in Psychiatry, Department of Psychiatry, School of Medicine, Universidad de Antioquia, Medellin, Colombia
91. Federal University of Bahia and PROAR Foudation, Salvador, Bahia, Brazil
92. UCL Genetics Institute, University College London
93. Institute for Molecular Medicine Finland, FIMM, HiLIFE, University of Helsinki, Helsinki, Finland
94. Institute of Health Sciences (ICS), Federal University of Bahia (UFBA), Salvador, Bahia, Brazil
95. Center for Precision Psychiatry, Massachusetts General Hospital, Harvard Medical School, Boston, MA, USA
96. Center for Genomic Medicine, Massachusetts General Hospital, Harvard Medical School, Boston, MA, USA
97. Genetics and Molecular Biology graduate program, Federal University of Pará, Belém, Pará, Brazil
98. Departament of specialized health, State University of Pará
99. Makerere University, Uganda
100. PAX Institute of Psychiatry, Brazil
101. Hai Phong Mental Health Hospital
102. Genomics Platform, The The Broad Institute of MIT and Harvard, Cambridge, MA, USA
103. Neuropsychiatric Hospital Aro, Abeokuta, Ogun State, Nigeria

104. Babcock University Teaching Hospital, Ilisan Remo, Ogun State, Nigeria
105. Texas Tech University Health Sciences Center El Paso
106. Professor Emeritus, Rutgers University, New Brunswick, NJ, USA
107. University of Tartu
108. Department of Psychology, University of Washington, Seattle, Washington
109. School of Medicine, Universidade de Fortaleza (UNIFOR)
110. Postdoctoral Research Fellow, Department of Psychiatry, Escola Paulista de Medicina, Universidade Federal de São Paulo
111. Beam Therapeutics
112. Department of Psychiatry, Shiga University of Medical Science, Japan
113. Department of Psychiatry, Escola Paulista de Medicina, Universidade Federal de São Paulo
114. Medical College East Africa, the Aga Khan Univeristy, Kenya
115. The Brain Clinic, Sialkot, Pakistan
116. Upstate Medical University, Syracuse New York
117. Department of Psychiatry and Biobehavioral Sciences, David Geffen School of Medicine at UCLA, Los Angeles, CA, USA
118. Program in Medical and Population Genetics, The Broad Institute of MIT and Harvard, Cambridge, Massachusetts, USA
119. Charité Universitätsmedizin, Berlin, Germany, Hanoi Medical University, Hanoi Vietnam
120. National Institute of Neuroscience, National Center of Neurology and Psychiatry, Japan
121. RIKEN Center for Brain Science, Japan
122. Department of Psychiatry, Kobe University Graduate School of Medicine, Japan
123. Hue Psychiatric Hospital
124. Nghe An Psychiatric Hospital
125. Shizuoka Graduate University of Public Health
126. Stanley Center for Psychiatric Research, The The Broad Institute of MIT and Harvard, Cambridge, MA, USA, BD2: Breakthrough Discoveries for thriving with Bipolar Disorder, Santa Monica, CA, USA
127. Department of Psychiatry, Taipei City Psychiatric Center, Taipei City Hospital, Taipei, Taiwan
128. Department of Psychiatry, School of Medicine, College of Medicine, Taipei Medical University, Taipei, Taiwan
129. Karolinska Institute, Sweden
130. Swat Medical Complex Teaching Hospital, Swat, Pakistan
131. Institute of Omics & Health Research, Lahore, Pakistan
132. Mian Iftikhar Psychiatry Hospital, Peshawar, Pakistan
133. Dr Idrees Private Clinic, Akbar Medical Centre, Peshawar, Pakistan
134. Nagoya University Graduate School of Medicine
135. Department of Neuropsychiatry and Clinical Ethics, Graduate School of Medical Science, University of Yamanashi, Yamanashi, Japan
136. Department of Psychiatry, Fujita Health University School of Medicine, Japan
137. Department of Epidemiology, Harvard T. H. Chan School of Public Health, Boston, MA, USA
138. Instituto Nacional de Psiquiatria Ramon de la Fuente Muñiz, Universidad Nacional Autónoma de México, Mexico city, Mexico
139. Dept of Psychiatry & Neuroscience Institute, University of Cape Town, South Africa
140. Department of Psychiatry, Icahn School of Medicine at Mount Sinai, New York, NY 10029, United States
141. Department of Neuropsychiatry, Graduate School of Medical Sciences, Kyushu University, Japan
142. African Population and Health Research Center, Nairobi, Kenya
143. Department of Psychiatry, Hokkaido University, Japan
144. Virginia Commonwealth University

145. Institute of Omics and Health Research, Lahore, Pakistan
146. Nai Zindagi Hospital, Multan, Pakistan
147. Dr Younas Khan Clinic, Charsadda, Pakistan
148. Khyber Medical University, Peshawar, Pakistan
149. Sarhad Mental Hospital Peshawar, Pakistan
150. Department of Medicine, Kyung Hee University College of Medicine, Seoul, Republic of Korea
151. Department of Medicine, Helsinki University Hospital, and Research Program for Clinical and Molecular Metabolism, University of Helsinki, Finland
152. Department of Psychiatry, Institute of Biomedical Sciences, Tokushima University Graduate School, Japan
153. Department of Psychiatry, Nara Medical University School of Medicine, Nara, Japan
154. Rutgers University, New Jersey, USA
155. Department of Social and Behavioral Sciences, Harvard T.H. Chan School of Public Health, Boston, MA, USA
156. Psychiatric & Neurodevelopmental Genetics Unit, Department of Psychiatry, Massachusetts General Hospital, Boston, MA, USA, BD2: Breakthrough Discoveries for thriving with Bipolar Disorder, Santa Monica, CA, USA
157. Department of Psychiatry, Teikyo University School of Medicine, Japan
158. Department of Public Health & Institute of Epidemiology and Preventive Medicine, National Taiwan University, Taipei, Taiwan
159. Department of Psychiatry, Nagoya University Graduate School of Medicine, Nagoya, Japan
160. Jinnah Postgraduate Medical Centre, Karachi, Pakistan
161. Kien Giang Psychiatric Hospital
162. DRPHI mental Clinic
163. Phu Tho Psychiatric Hospital
164. Thai Binh Mental Health Hospital
165. Thanh Hoa Psychiatric Hospital
166. Queensland Institute of Medical Research
167. Deparment of Psychiatry, School of Medicine, Universidad de Antioquia, Medellin, Colombia
168. Department of Psychiatry, Massachusetts General Hospital
169. Quaid e Azam Medical College, Bahawalpur, Pakistan
170. Stanley Center for Psychiatric Research, The Broad Institute of MIT and Harvard, Cambridge, MA, USA
171. BD2: Breakthrough Discoveries for thriving with Bipolar Disorder, Santa Monica, CA, USA
172. Helsinki University Central Hospital, Helsinki, Finland
173. University College London
174. Haji Abdul Qayyum Hospital, Sahiwal, Pakistan
175. Arrahma Hospital for Mental Health, Multan, Pakistan
176. Institute of Biomedicine, University of Eastern Finland, Kuopio, Finland
177. Aichi Medical University
178. University of Galway, Ireland
179. Psychosis Research Unit, Aarhus University Hospital-Psychiatry, Aarhus, Denmark
180. Aarhus University
181. Ibadat Hospital, Peshawar, Pakistan
182. Department of Psychiatry, Kyoto University Graduate School of Medicine, Japan
183. Sheikh Zayed Medical College and Hospital, Rahim Yar Khan, Pakistan
184. Brain and Mind Institute, the Aga Khan University, Kenya
185. KEMRI, Kenya
186. Khanh Hoa Psychiatric Hospital
187. Mental Health center, Lao Cai Province General Hospital No. 1
188. Hanoi Medical University, Vietnam

189. Ho Chi Minh Psychiatric Hospital
190. Vinh Phuc Psychiatric Hospital
191. National Psychiatric Hospital No. 1
192. Department of Mental Health, Medical and pharmaceutical University Hospital, Vietnam National University, Hanoi
193. National Psychiatric Hospital No. 2
194. Hung Yen Mental Health Hospital
195. Hanoi Medical University, National Institut of Mental Health, Vietnam
196. Hanoi Medical University, Hanoi Vietnam
197. Hanoi Medical University, Vietnam, National Geriatric Hospital
198. Institute of Psychiatry, Rawalpindi Medical University, Rawalpindi, Pakistan
199. Obafemi Awolowo University Teaching Hospitals Complex, Ile-Ife, Osun State, Nigeria
200. The Wolfson Centre for Young People's Mental Health, Cardiff University, United Kingdom
201. Department of Psychiatry, Institute of Biomedical Sciences, Tokushima University Graduate School, Japan
202. Department of Physiology, Keio University School of Medicine, Japan
203. Department of Psychiatry, University of Cambridge
204. Department of Medical Laboratory Science, College of Medicine, University of Lagos, Nigeria
205. Department of Computational Medicine, David Geffen School of Medicine, UCLA, Los Angeles, CA, USA
206. Department of Human Genetics, David Geffen School of Medicine, UCLA, Los Angeles, CA, USA
    1. Hospital de Saúde Mental Professor Frota Pinto, Fortaleza, Ceará, Brazil 2. Department of Pharmacology, Federal University of Ceará, Fortaleza, Ceará, Brazil
207. McLean Hospital, Harvard Medical School, Belmont, MA, USA
208. Center for Neurobehavioral Genetics, Semel Institute for Neuroscience and Human Behavior, David Geffen School of Medicine, University of California Los Angeles, Los Angeles, CA, USA
209. University of Cambridge, Cambridge, UK
210. Department of Neurology, Massachusetts General Hospital, Boston, MA, USA
211. Department of Psychiatry, Massachusetts General Hospital, Boston, MA, USA
212. Rutgers University, New Brunswick, NJ, USA
213. Department of Medical Epidemiology and Biostatistics, Karolinska Institutet, Sweden
214. Nam Dinh Psychiatric Hospital
215. VU University Amsterdam
216. School of Medicine, Johns Hopkins University, Baltimore, MD, USA
217. Hyderabad Consulting Chambers, Hyderabad, Pakistan
218. Akram Hospital, Quetta, Pakistan
219. Department of Psychiatry, Universitats Klinikum Frankfurt, Germany
220. Department of Psychiatry and Behavioral Sciences, School of Medicine, University of California, San Francisco, San Francisco, California, USA
221. Laboratory of Neuropsychopharmacology, Psychiatry Service, Universidade Federal da Bahia, Salvador, BA, Brazil
222. The University of the West Indies, Cave Hill Campus, Barbados, West Indies
223. Folkhälsan Research Center and Translational Immunology Research Program, University of Helsinki, Finland
224. Department of Biomedical Data Science, Stanford, CA, USA
225. Escola Bahiana de Medicina de Saúde Pública and PROAR Foundation, Salvador, Bahia, Brazil
226. Disciplina de Biologia Molecular, Universidade Federal de São Paulo, São Paulo, SP, Brazil
227. Johns Hopkins Schizophrenia Center and Johns Hopkins iMIND, Dept. of Psychiatry, Neuroscience, Biomedical Engineering, Pharmacology, Genetic Medicine, and Mental Health, Johns Hopkins University School of Medicine and Bloomberg School of Public Health

228. Department of Human Genetics, UCLA
229. Shakoor Mind Care Institute, Bahawalpur, Pakistan
230. Department of Psychiatry, Dokkyo Medical University School of Medicine, Japan
231. Columbia University: New York, NY, US
232. Psychiatric and Neurodevelopmental Genetics Unit, Massachusetts General Hospital, Boston, MA, USA
233. Department of Psychiatry, Harvard Medical School, Boston, MA, USA
234. SAMRC Unit on Risk & Resilience in Mental Disorders, Dept of Psychiatry & Neuroscience Institute, University of Cape Town, South Africa
235. Univeristy of North Carolina, Chapel Hill, North Carolina, USA
236. Sayed Psychiatric Hospital, Peshawar, Pakistan
237. Department of Neuropsychiatry, University of Toyama Graduate School of Medicine and Pharmaceutical Sciences, Japan
238. Pakistan Institute of Medical Sciences, Islamabad, Pakistan
239. RIKEN Center for Brain Science, Japan
240. Department of Neuropsychiatry, Faculty of Life Sciences, Kumamoto University, Japan
241. Stanley Center for Psychiatric Research, Broad Institute of MIT and Harvard, Cambridge, MA 02142, USA
242. Program in Bioinformatics and Integrative Genomics, Harvard Medical School, Boston, MA 02115, USA
243. City university of hong kong, Department of Biomedical Sciences, Hong Kong, China
244. MRC Integrative Epidemiology Unit at the University of Bristol
245. Department of Psychiatry, Kumamoto University, Japan
246. Bac Ninh Provincial Mental Health Hospital No. 2
247. Da Nang Psychiatric Hospital
248. Ben Tre Psychiatric Hospital
249. Ha Nam Psychiatric Hospital
250. Dong Thap Psychiatric Hospital
251. Bac Ninh Provincial Mental Health Hospital No. 1
252. University of California, San Diego, USA
253. Dow University of Health Sciences, Karachi, Pakistan
254. University of California, Irvine, USA
255. National Institute of Mental Health and Neurosciences, Bangalore, India
256. Can Tho Psychiatric Hospital
257. Department of Psychology, Education & Child Studies, Erasmus School of Social and Behavioural Sciences, Erasmus University Rotterdam, The Netherlands, Department of Child and Adolescent Psychiatry/Psychology, Erasmus University Medical Center, Rotterdam, The Netherlands
258. Quang Ninh Mental Health Hospital
259. Hanoi Psychiatric Hospital
260. MRC Centre for Neuropsychiatric Genetics and Genomics, Cardiff University, UK
261. Mental Health Centre Copenhagen, Capital Region of Denmark, Copenhagen University Hospital, Copenhagen, Denmark
262. The Lundbeck Foundation Initiative for Integrative Psychiatric Research, iPSYCH, Copenhagen, Denmark
263. Department of Psychiatry, Johns Hopkins University School of Medicine
264. Stanley Division of Developmental Neurovirology, Johns Hopkins University, Maryland, USA
265. Khushal Medical Centre Peshawar, Pakistan
266. Department of Psychiatry and Behavioral Sciences, Johns Hopkins School of Medicine, Baltimore, Maryland, USA
267. Nelson Mandela University, South Africa
268. Alee’z Neuro Psychiatric Centre, Sargodha, Pakistan

**Cohort Descriptions.**

The GEN-SCRIP Study (GENetics of SChizophRenia in Pakistan)
**Consortium/Collaboration:** Pakistan Alliance on genetic RisK factors for Health (PARKH)
**Investigator:** James Knowles
**Country:** Pakistan
**Sample size:** 7647 cases, 7648 controls
**Description:** The GenSCRIP study will ascertain and genotype large cohort of Pakistani individuals with (n=8,000) and without (n=9,000) schizophrenia (SCZ). Added to our existing Pakistani samples, a total of 10,000 cases and 10,000 controls will be included in genome-wide association studies and these data will be contributed to larger meta-analyses.
**Ethics/IRB approval:** Approved by Rutgers University IRB and the National Bioethics Committee of Pakistan
**Acknowledgements:** The authors would like to acknowledge and thank the study participants and their families. The GEN-SCRIP study is a public-private partnership between the National Institute of Mental Health (NIMH) and the Stanley Center for Psychiatric Research. This work was supported by a grant, R01 MH112904 (J.A.K), from the National Institute of Mental Health (NIMH)
**Data availability:** NIMH Data ARchive (NDA)

Asian Bipolar Genetics Network (A-BIG-NET)
**Consortium/Collaboration:** ABIGNET Consortium
**Investigator:** Hailiang Huang
**Country:** Taiwan, Singapore, Korea, India, and Vietnam
**Description:** A-BIG-NET is a large multi-site research study, planning to collect over 30,000 bipolar cases and controls in five Asia countries - Taiwan, Singapore, Korea, India, and Vietnam. The study will collect a rich array of phenotyping information in Asian population.

Neuropsychiatric Genetics of African Populations (NeuroGAP)-Psychosis
**Consortium/Collaboration:** African Collections
**Investigator:** Karestan Koenen
**Country:** Ethiopia, Kenya, South Africa, and Uganda
**Sample size:** 9546 cases, 16406 controls
**Description:** Neuropsychiatric Genetics of African Populations (NeuroGAP)-Psychosis is an ongoing case-control study of participants recruited across multiple sites within Ethiopia, Kenya, South Africa, and Uganda. All participants provided psychiatric, behavioral, and demographic data, as well as a range of phenotypic data including, but not limited to, substance use information and history of traumatic events. Saliva was also collected for DNA extraction.
**Ethics/IRB approval:** Ethical clearances to conduct this study have been obtained from all participating sites, including: • Ethiopia: Addis Ababa University College of Health Sciences (#014/17/Psy) and the Ministry of Science and Technology. National Research Ethics Review Committee (#3.10/14/2018). Kenya: Moi University College of Health Sciences/Moi Teaching and Referral Hospital Institutional Research and Ethics. Committee (#IREC/2016/145, approval number: IREC 1727), Kenya National Council of Science and Technology (#NACOSTI/P/17/56302/19576) KEMRI Centre Scientific Committee (CSC# KEMRI/CGMRC/CSC/070/2016), KEMRI Scientific and Ethics Review Unit (SERU#KEMRI/SERU/CGMR-C/070/3575). South Africa: The University of Cape Town Human Research Ethics Committee (#466/2016) and Walter Sisulu University Research and Ethics Committee (# 051/2016). Uganda: The Makerere University School of Medicine Research and Ethics Committee (SOMREC #REC REF 2016-057) and the Uganda National Council for Science and Technology (UNCST #HS14ES). USA: The Harvard T.H. Chan School of Public Health (#IRB17-0822).
**Acknowledgements:** We would like to acknowledge the data managers, clinicians, research assistants, and project managers who have worked on this study from Addis Ababa University: Seble Abate, Lidia Abebaye, Tarikua Abera, Beakal Amare, Biruh Alemayehu, Melkam Assefa, Habtamu Assegid, Mihret Daniel, Samrawit Daniel, Wubit Demeke, Harun Esmael, Tolessa Fanta, Sintayehu Gurmessa, Belete Habtewold, Yodit Habtamu, Kelemua Haile, Mena Hibist, Desalegn Kefiyebel, Mohammed Negussie, Agitu Tadesse, Mickiyas Tilahun, Mikiyas Tullu, and Degu Zenab; from KEMRI-Wellcome Trust: Phanice Amukhale, Mary Bitta, Patrick Tsuma Idd, Moses Mangi, Eric Mwajombo, Sylvia Mwamba, Paul Mwangi, Amina Mwinyi, Musa Mzee, Mercy Mzungu, Hamisi Rashid, Branis Widole; from Makerere University: Adiru Tamali, Apio Racheal, Clare Samba Nalwoga, Francis Ojara, Julius Okura, Naome Nyinomugisha, Samalie Nsangi, Stanley Baniyo, and Stella Anena; from Moi University/Moi

Teaching and Referral Hospital: Mohamed Aden, Sarah Busienei, Eunice Jeptanui, Kimutai Katwa, Wilberforce Ndenga, and Fredrick Ochieng; from Harvard/Broad Institute: Iman Ali, Mark Baker, Justin McMahon, and Sophie Greenebaum; from the University of Cape Town: Bronwyn Malagas, Bukeka Sawula, Deborah Jonker, Linda Ngqengelele, Michaela De Wet, Nabila Ebrahim, Ncumisa Nzenze, Onke Maniwe, Phelisa Bashman, Sibonile Mqulwana, Sibulelo Mollie, Renier Swart, Roxanne James, Tyler Linnen, and Xolisa Sigenu. We would also like to acknowledge the participants who shared their time and their experiences with us. Without them, this work would not be possible.

BRIDGES
**Consortium/Collaboration:** African Collections
**Investigator:** Niran Okewole
**Country:** Nigeria
**Sample size:** 197 cases, 140 controls
**Description:** Participants were recruited at five sites in southwest Nigeria (Lagos, Abeokuta, Ile-Ife, Ado-Ekiti and Ilisan-Remo). Ethics approvals were obtained from each study site and from the National Health Research Ethics Committee. Subjects with schizophrenia were interviewed with the PANSS, CGI, GAF and WHODAS 2.0. Control participants completed the PSQ and Kessler Scale. All participants completed a sociodemographic questionnaire, CIDI, CTQ, LEC, ASSIST, morphometry and vitals. All participants provided saliva samples using standard DNA kits for subsequent DNA analysis.
**Ethics/IRB approval:** National ethics (NHREC) Approval Number NHREC/01/01/2007- 20/11/2022
Project Among African-Americans to Explore Risks for Schizophrenia (PAARTNERS)
**Consortium/Collaboration:** African Collections
**Investigator:** Rodney CP Go
**Country:** USA
**Sample size:** 22 cases, 760 controls
**Description:** The Project among African-Americans to explore risks for schizophrenia (PAARTNERS) included probands meeting DSM-IV criteria for schizophrenia or schizoaffective disorder, depressed type, their relatives (as trios, affected sibling pairs, or multiplex pedigrees), and community controls (14,15). Participants were recruited in the Southeastern and Eastern USA, self-identified as African American, and were assessed using the Diagnostic Interview for Genetic Studies, Family Interview for Genetic Studies, the Penn-CNB, and medical records review. Blood or buccal cells were collected for genetic analyses. Probands and ancestry-matched, psychiatrically unscreened controls from MiGEN 16 were genotyped on the MEGA.

ClozaGene study
**Consortium/Collaboration:** Australian Collections
**Investigator:** Sarah Medland
**Country:** Australia
**Sample size:** 600 cases, 3265 controls
**Description:** The ClozaGene study recruited individuals who had been prescribed Clozapine for treatment resistant schizophrenia from across Australia (DOI: 10.1093/schbul/sbae065). Controls were drawn from among the participants of the QSkin study (DOI: 10.1093/ije/dys107). The samples were sent to the Broad Institute for Blended Genome Exome sequence generation and included in this meta-analysis.
**Ethics/IRB approval:** Ethics approval was obtained from the QIMR Berghofer Human Research Ethics Committee (P3556; P2034;P3434)
**Acknowledgements:** We thank the participants for giving their time and support for this project. This work was supported by the Australian National Health and Medical Research Council Grants (APP1138514, APP1194635, APP1172917 and APP2025674).
**Data availability:** Anonymized survey data may be shared for collaborative projects, subject to a data transfer agreement and governance and ethics approval.
Precision medicine study of psychiatric disease in Chinese population (PMPDC)
**Consortium/Collaboration:** BioX Collection
**Investigator:** Shengying Qin
**Country:** China
**Sample size:** 13068 cases, 12418 controls
**Description:** This study contains 8732 Chinese schizophrenia cases and 8660 controls

Finnish Collection - SUPER
**Consortium/Collaboration:** Finnish Collections
**Investigator:** Aarno Palotie
**Country:** Finland
**Sample size:** 5270 cases
**Description:** The Finnish SUPER study on genetic mechanisms of psychotic disorders is a part of the international Stanley Global Neuropsychiatric Genomics Initiative (The Stanley Center for Psychiatric Research at the Broad Institute of MIT and Harvard) The objective of the study is to better understand the genetic and biological background of psychotic disorders in order to provide more accurate information for the development of new therapeutic interventions. The national study was launched in early 2016 and the sample collection was completed by the end of 2018. During this time, SUPER study recruited 10,470 individuals. Inclusion criteria was at least one psychotic episode during life-time. Patients were contacted during normal course of treatment at hospitals, nursing facilities and health care centers. Five university hospital districts were involved in sample collection. The study was coordinated by Institute for Molecular Medicine Finland (FIMM), University of Helsinki and the Finnish Institute for Health and Welfare (THL).

Finnish Collection - Finnish Biobank
**Consortium/Collaboration:** Finnish Collections
**Investigator:** Aarno Palotie
**Country:** FInland
**Sample size:** 1627 cases, 1668 controls
**Description:** The mission of Finnish Biobank Cooperative – FINBB is to enhance the competitiveness of Finnish health and biomedical research by providing researchers a centralized access to collections and services of the Finnish biobanks and their background organizations. FINBB's activities create value for its members and owners by providing services at the national level.

Japan
**Consortium/Collaboration:** Japanese Collections
**Investigator:** Akira Sawa
**Country:** Japan
**Sample size:** 3917 cases, 5133 controls
**Description:** The Japanese cohorts were onboarded and collected as part of the Stanley Center's Asia Initiative to increase global collections. The Japanese Exome study expands the sample cohorts across East Asia to help with fine mapping and ; 1) complete the allelic spectrum of schizophrenia 2) confirm that genetic effects are shared across ancestries and 3) fine-map the causal variant. Yasue Horiuchi at Shizuoka Graduate University of Public Health and Akira Sawa at Johns Hopkins University led efforts across 15 sites to recruit schizophrenia cases and matching controls.

UCLA External Data
**Consortium/Collaboration:** North American Collaborations
**Investigator:** Multiple
**Country:** USA
**Sample size:** 175 cases, 26 controls
**Description:** These samples are from a set of studies listed here: Transmission of Vulnerability Factors for Schizophrenia, Center Aftercare – II, Genetic Risk Factors, CT&E R01, Familial Psychiatric Disorders and Attention in Schizophrenia, COGS all conducted at the University of California Los Angeles. These data were sequenced externally, and reprocessed for inclusion in larger SCHEMA analyses

MGB Biobank
**Consortium/Collaboration:** North American Collaborations
**Investigator:** Elizabeth Karlson
**Country:** United States
**Sample size:** 155 cases, 2711 controls
**Description:** The MGB (formerly Partners) Biobank (https://biobank.massgeneralbrigham.org/), launched in 2010, is a biorepository of consented patients samples at Mass General Brigham (parent organization of Massachusetts General Hospital and Brigham and Women's Hospital). The Biobank has enrolled >100K individuals to study how

genes, lifestyle, and other factors affect people's health and contribute to disease. As part of the NHGRI's Centers for Common Disease Genomics, Broad Institute of MIT and Harvard generated genetic data for ~13,500 individuals from the MGB Biobank. Data is available by application to dbGaP/AnVIL under accession number phs0002018.
**Acknowledgements:** MGB Biobank: We gratefully acknowledge the participants and leadership team of the MGB Biobank, funding support from the NHGRI CCDG (UM1HG008895), and generation of new whole exome sequencing data by the Broad Genomics Platform.
**Data availability:** Mass General Brigham Biobank Portal: Qualified investigators can access these integrated resources via the web-based biobank portal subject to institutional approvals.

PUMAS - LATAM
**Consortium/Collaboration:** PUMAS Consortium
**Investigator:** Nelson Freimer
**Country:** Brazil
**Sample size:** 2624 cases, 5023 controls
**Description:** The PUMAS-LATAM cohort is a case-control study of 14,638 participants recruited from five major cities from different regions in Brazil (São Paulo, Fortaleza, Salvador, Goiânia, Belém), and one coastal city in Colombia (Cartagena). Participants included individuals who were at least 18 years old at the time of recruitment with a clinical diagnosis of SZA, SCZ, and BP, and matched controls. Potential cases and controls were screened with the GPC screener. All cases were assessed using the DI-PAD interview, with psychiatric diagnoses made based on an overall assessment by the recruiting clinician. Item-level phenotypes were also derived from this instrument.

PUMAS - Mision Origen
**Consortium/Collaboration:** PUMAS Consortium
**Investigator:** Carlos Lopez-Jaramill
**Country:** Colombia
**Sample size:** 2391 cases, 14354 controls
**Description:** Through Misión Origen, we are creating a biobank for Severe Mental Illness, consisting of 50,000 participants with severe mental illness and 50,000 population controls from the Paisa region of Colombia. All phenotyping will be through electronic medical records.Co-led by Loes Olde Loohuis, Carlos Lopez Jaramillo, and Nelson Freimer. Funded by NIMH. For some of our ongoing work in this population, see: Service et al. 2020, Lancet Psychiatry and Mooney et al. 2018, American Journal of Human Genetics, Service et al. 2023, Song et al. 2024, De la Hoz et al 2022.

PUMAS - NeuroMex
**Consortium/Collaboration:** PUMAS Consortium
**Investigator:** Beatriz Camarena
**Country:** Mexico
**Sample size:** 1823 cases, 2529 controls
**Description:** The NeuroMex study is a collaboration between the Broad Institute, Harvard T.H. Chan School of Public Health and the Ramón de la Fuente Muñiz National Institute of Psychiatry (INPRFM) in Mexico. The study aims to expand knowledge of the genetic and environmental risk factors for neuropsychiatric disorders in Mexico through large-scale sample collection and analysis, so that future advances in science and therapeutics can account for and be applicable to Mexican populations.

PUMAS - Paisa
**Consortium/Collaboration:** PUMAS Consortium
**Investigator:** Nelson Freimer
**Country:** Colombia
**Sample size:** 1376 cases, 7474 controls
**Description:** The Paisa Project is a case-control study of 9,105 participants recruited from the Paisa genetic isolate of Colombia. Participants included individuals from the Paisa region who were at least 18 years old at the time of recruitment with a clinical diagnosis of SZA, SCZ, BP, or severe MDD, and matched controls. Cases were recruited from two large psychiatric hospitals in the region. Potential controls were screened using the overview module of the online version of the SCID-5 (NetSCID-5, Spanish Version)28. Psychiatric diagnoses were determined through the NetSCID interview. Item-level symptoms were extracted from the i) NetSCID interview, and ii) Supplemental

Symptoms Questionnaire (SSQ), an 8-item questionnaire designed to assess transdiagnostic symptoms that the NetSCID's branching logic did not capture.

PUMAS - Colombia
**Consortium/Collaboration:** PUMAS Consortium
**Investigator:** Nelson Freimer
**Country:** Colombia
**Sample size:** 695 cases, 1951 controls
**Description:** The PUMAS-LATAM cohort is a case-control study of 14,638 participants recruited from five major cities from different regions in Brazil (São Paulo, Fortaleza, Salvador, Goiânia, Belém), and one coastal city in Colombia (Cartagena). Participants included individuals who were at least 18 years old at the time of recruitment with a clinical diagnosis of SZA, SCZ, and BP, and matched controls. Potential cases and controls were screened with the GPC screener. All cases were assessed using the DI-PAD interview, with psychiatric diagnoses made based on an overall assessment by the recruiting clinician. Item-level phenotypes were also derived from this instrument.

PUMAS - LA
**Consortium/Collaboration:** PUMAS Consortium
**Investigator:** Nelson Freimer
**Country:** USA
**Sample size:** 1033 cases, 0 controls

Cardiff
**Consortium/Collaboration:** SCHEMA 1 Collections
**Investigator:** Mick O'Donovan
**Country:** United Kingdom
**Sample size:** 6548 cases, 518 controls
**Description:** The case sample included European ancestry schizophrenia cases recruited in the British Isles and described previously. All cases gave written informed consent. The study was approved by the Multicentre Research Ethics Committee in Wales and Local Research Ethics Committees from all participating sites. The control sample used the Wellcome Trust Case Control Consortium (WTCCC) sample described elsewhere, but included similar numbers of individuals from the 1958 British Birth Cohort and a panel of consenting blood donors (UK Blood Service).

Genomic Psychiatry Cohort (GPC)
**Consortium/Collaboration:** SCHEMA 1 Collections
**Investigator:** Carlos Pato
**Country:** USA
**Sample size:** 4914 cases, 7360 controls
**Description:** We recruited participants as part of the Genomic Psychiatry Cohort (GPC), a study based in the Los Angeles that recruited controls and cases living and being treated in local communities and healthcare delivery systems. This sample has been described elsewhere. Case participants were interviewed using the Diagnostic Interview for Psychosis and Affective Disorders (DI-PAD), a semi-structured clinical interview administered by mental health professionals. Inclusion criteria or cases included meeting lifetime diagnostic criteria for schizophrenia or schizoaffective disorder (any subtype) in accordance with the OPCRIT algorithms for DSM-IV and/or ICD-10 criteria, and/or DSM-5. Individuals reporting no lifetime symptoms indicative of psychosis or mania and who had no first-degree relatives with these symptoms were included as control participants. Exclusion criteria included any premorbid organic mental disorders and premorbid history of significant drug or alcohol dependence by DSM-IV/5 that confounds the diagnosis of schizophrenia. DNA was extracted from whole blood. All participants gave written informed consent and the IRB of the participating institutions approved the protocol.
**Acknowledgements:** We thank the Genomic Psychiatry Cohort (GPC) Consortium teams including: Evelyn J. Bromet, Celia Barreto Carvalho, Eric D. Achtyes, Maria Helena Azevedo, Roman Kotov, Douglas S. Lehrer, Dolores Malaspina, Stephen R. Marder, Helena Medeiros, Christopher P. Morley, Diana O. Perkins, Janet L Sobell, Peter F. Buckley, Fabio Macciardi, Mark H. Rapaport, James A. Knowles, and Ayman H. Fanous.

Sweden Schizophrenia Study
**Consortium/Collaboration:** SCHEMA 1 Collections

**Investigator:** Patrick F Sullivan
**Country:** Sweden
**Sample size:** 3961 cases, 5447 controls
**Description:** Fully described in PMID 23974872.
**Ethics/IRB approval:** Approved by all relevant IRBs.
**Acknowledgements:** Swedish Research Council (Vetenskapsrådet, award D0886501); NIMH R01 MH077139; NIMH U01 MH109528

CLOZUK
**Consortium/Collaboration:** SCHEMA 1 Collections
**Investigator:** Multiple
**Country:** United Kingdom
**Sample size:** 3605 cases, 0 controls
**Description:** We collected blood samples from those with treatment-resistant schizophrenia (TRS) in the UK through the mandatory clozapine blood-monitoring system for those taking clozapine, an antipsychotic licensed for TRS. Following national research ethics approval and in line with UK Human Tissue Act regulations we worked in partnership with the commercial companies that manufacture and monitor clozapine in the UK. We ascertained anonymous aliquots of the blood samples collected as part of the regular blood monitoring that takes place whilst taking clozapine due to a rare haematological adverse effect, agranulocytosis. The sample was assembled in collaboration with Leyden Delta (Nijmegen, Netherlands), a major company involved in the supply Treatment Access System (ZTAS), provided whole-blood samples and anonymised phenotypic information. Both Clozaril® and Zaponex® are bioequivalent brands of clozapine licensed in the UK.

Spain
**Consortium/Collaboration:** SCHEMA 1 Collections
**Investigator:** Celso Arango
**Country:** Spain
**Sample size:** 1651 cases, 1162 controls
**Description:** Participants were recruited as part of the sample collection of CIBERSAM (Network Biomedical Research Centre in Mental Health), an institution based in Spain that recruits in-patients from psychiatric units at eight different hospitals in Spain. All patients recruited met the DSM-IV diagnostic criteria for schizophrenia, schizoaffective or schizophreniform disorder. Healthy controls were recruited after being screened for psychiatric illness. All participants provided written informed consent. Blood sample recruitment and preparation, participation in the PGC consortium and commitment to share the generated data in subsequent meta-analyses and further secondary proposals were approved by the different ethical committees at the hospitals involved in the recruitment.

UK/Ireland Controls
**Consortium/Collaboration:** SCHEMA 1 Collections
**Investigator:** Andrew Mcquillin
**Country:** United Kingdom
**Sample size:** 1396 cases, 1506 controls
**Description:** The UCL control sample consisted of 480 genomic DNA samples that were extracted from EBV transformed peripheral blood lymphocytes from unscreened healthy British blood donors (https://www.phe-culturecollections.org.uk/products/dna/hrcdna/hrcdna). The remaining DNA samples were extracted from whole blood samples from healthy volunteers of UK or Irish ancestry who were 50 interviewed with the initial clinical screening questions of the SADS-L and selected on the basis of not having a past or present personal history of any RDC-defined mental disorder. Heavy drinking and a family history of schizophrenia, alcohol dependence or bipolar disorder, were also used as exclusion criteria for controls. UK National Health Service multi-centre and local research ethics approvals were obtained and all subjects gave signed informed consent.5 Whole exome sequencing was done by the Broad Genomics Platform and data is available by application to the European Genome Phenome Archive (EGA) under accession number EGAS00001005851.
**Ethics/IRB approval:** The UCL control sample consisted of 480 genomic DNA samples that were extracted from EBV transformed peripheral blood lymphocytes from unscreened healthy British blood donors (https://www.phe-culturecollections.org.uk/products/dna/hrcdna/hrcdna). The remaining DNA samples were extracted from whole blood samples from healthy volunteers of UK or Irish ancestry who were 50 interviewed with the initial clinical screening questions of the SADS-L and selected on the basis of not having a past or present personal history of any

RDC-defined mental disorder. Heavy drinking and a family history of schizophrenia, alcohol dependence or bipolar disorder, were also used as exclusion criteria for controls. UK National Health Service multi-centre and local research ethics approvals were obtained and all subjects gave signed informed consent.5 Whole exome sequencing was done by the Broad Genomics Platform and data is available by application to the European Genome Phenome Archive (EGA) under accession number EGAS00001005851.
**Acknowledgements:** The UCL control sample consisted of 480 genomic DNA samples that were extracted from EBV transformed peripheral blood lymphocytes from unscreened healthy British blood donors (https://www.phe-culturecollections.org.uk/products/dna/hrcdna/hrcdna). The remaining DNA samples were extracted from whole blood samples from healthy volunteers of UK or Irish ancestry who were 50 interviewed with the initial clinical screening questions of the SADS-L and selected on the basis of not having a past or present personal history of any RDC-defined mental disorder. Heavy drinking and a family history of schizophrenia, alcohol dependence or bipolar disorder, were also used as exclusion criteria for controls. UK National Health Service multi-centre and local research ethics approvals were obtained and all subjects gave signed informed consent.5 Whole exome sequencing was done by the Broad Genomics Platform and data is available by application to the European Genome Phenome Archive (EGA) under accession number EGAS00001005851.

Ashkenazi Jewish schizophrenia study
**Consortium/Collaboration:** SCHEMA 1 Collections
**Investigator:** Ann Pulver
**Country:** United States
**Sample size:** 731 cases, 543 controls
**Description:** Samples from Johns Hopkins University were provided for sequencing.

SCHEMA 1 Collections
**Investigator:** David St. Clair, Mick O'Donovan
**Country:** United Kingdom
**Sample size:** 516 cases, 350 controls
**Description:** Ascertainment and inclusion/exclusion criteria for cases and controls have been previously described73. All participating subjects were born in the UK (95% Scotland) and gave written informed consent. Both local and multiregional academic ethical committee approved the human subjects protocol. The samples were genotyped at the Broad Institute.

McLean Schizophrenia Collection
**Investigator:** Bruce Cohen
**Country:** United States
**Sample size:** 428 cases, 229 controls

Cardiff COGs
**Consortium/Collaboration:** SCHEMA 1 Collections
**Investigator:** James Walters
**Country:** United Kingdom
**Sample size:** 330 cases, 0 controls
**Description:** Cases were recruited from community mental health teams in Wales and England on the basis of a clinical diagnosis of schizophrenia or schizoaffective disorder (depressed sub-type) as described previously125. Diagnosis was confirmed following a SCAN90 interview and review of case notes followed by consensus diagnosis according to DSM-IV51 criteria. The samples were genotyped at the Broad Institute. The UK Multicentre Research Ethics Committee (MREC) approved the study and all participants provided valid informed consent.

UK/Ireland Controls 2
**Consortium/Collaboration:** SCHEMA 1 Collections
**Investigator:** Douglas Blackwood
**Country:** UK/Ireland
**Sample size:** 276 cases, 59 controls
**Description:** Participants were recruited from clinical service around Edinburgh and Scotland and screened using the SADS-L. DNA samples were extracted from whole blood for genotyping and sequencing. Research was conducted after research ethics and NHS management approvals (Schizophrenia Working Group of the Psychiatric

Genomics, C). Whole exome sequencing was done by the Broad Genomics Platform and data is available by application to the European Genome Phenome Archive (EGA) under accession number EGAS00001005843.
**Ethics/IRB approval:** UK/Ireland controls 1 were collected with support from the Neuroscience Research Charitable Trust, the Central London NHS (National Health Service) Blood Transfusion Service and the National Institute of Health Research (NIHR) funded Mental Health Research Network.

The Study of Genetics in Mental Illness
**Consortium/Collaboration:** SCHEMA 1 Collections
**Investigator:** Michael Owen
**Country:** United Kingdom
**Sample size:** 274 cases, 0 controls

Estonia Biobank
**Consortium/Collaboration:** SCHEMA 1 Collections
**Investigator:** Tõnu Esko
**Country:** Estonia
**Sample size:** 219 cases, 0 controls
**Description:** The Estonian cohort comes from the population-based biobank of the Estonian Genome Project of University of Tartu (EGCUT)85. The project was conducted according to the Estonian Research Act and all participants provided informed consent (www.biobank.ee). In total, 52,000 individuals aged 18 years or older participated in this cohort (33% men, 67% women). The population distributions of the cohort reflect those of the Estonian population (83% Estonians, 14% Russians and 3% other). General practitioners (GP) and physicians in the hospitals randomly recruited the participants. A Computer-Assisted Personal interview was conducted over 1-2 ours at doctors' offices. Data on demographics, genealogy, educational and occupational history, lifestyle and anthropometric and physiological data were assessed. Schizophrenia was diagnosed prior to the recruitment by a psychiatrist according to ICD-1052 criteria and identified from the Estonian Biobank phenotype database. Controls were drawn from a larger pool of genotyped biobank samples by matching on gender, age and genetic ancestry. All the controls were population based and have not been sampled for any specific disease.

Cardiff
**Consortium/Collaboration:** SCHEMA 1 Collections
**Investigator:** Nicholas Craddock
**Country:** United Kingdom
**Sample size:** 46 cases, 0 controls
**Description:** Cases were all over the age of 17 yr, living in the UK and of European descent. Recruitment was undertaken throughout the UK and included individuals who had been in contact with mental health services and had a lifetime history of high mood. After providing written informed consent, participants were interviewed by a trained psychologist or psychiatrist using a semi-structured lifetime diagnostic psychiatric interview (Schedules for Clinical Assessment in Neuropsychiatry) and available psychiatric medical records were reviewed. Using all available data, best-estimate life-time diagnoses were made according to the RDC12. In the current study we included cases with a lifetime diagnosis of RDC bipolar 1 disorder, bipolar 2 disorder or schizo-affective disorder, bipolar type. Controls were recruited from two sources: the 1958 Birth Cohort study and the UK Blood Service (blood donors) and were not screened for history of mental illness. All cases and controls were recruited under protocols approved by the appropriate IRBs. All subjects gave written informed consent.

BPD
**Consortium/Collaboration:** SCHEMA 1 Collections
**Investigator:** Aiden Corvin
**Country:** Ireland
**Sample size:** 29 cases, 6 controls
**Description:** The case sample was collected primarily in the Dublin area and the ascertainment procedure has been previously described. The controls were recruited, from the same region through the Irish Blood Transfusion Services. All participants gave written, informed consent and the collections were approved through the Federated Dublin Hospitals and Irish Blood Transfusion Services Research Ethics Committees, respectively. DNA samples were genotyped at the Broad Institute.
**Ethics/IRB approval:** Detailed already via refs in papers above.

Spatiotemporal dynamics of contextual processing in schizophrenia, autism spectrum disorder, and obsessive-compulsive disorder
**Consortium/Collaboration:** SCHEMA 1 Collections
**Investigator:** Dana Manoach
**Country:** United States
**Sample size:** 26 cases, 27 controls

WGSPD
**Consortium/Collaboration:** WGSPD - Whole Genome Sequencing for Schizophrenia and Bipolar Disorder
**Investigator:** Nelson Freimer, Elliot Hong, Raquel Gur, Carlos Pato, Ann Pulver, David Glahn, Roel Ophoff
**Country:** Netherlands, United States, United Kingdom
**Sample size:** 4,688 cases, 3,564 controls
**Description:** The WGSPD study sought to increase the discovery of common and low frequency variants and rare coding variants found to be associated with the risk of bipolar disorder (BD) and schizophrenia (SZ) beyond individuals of European genetic ancestry, which had primarily been the focus of previous efforts. The group performed whole genome sequencing of African American and African ancestry individuals: 1,598 individuals with BD, 3,295 individuals with SZ, and 2,651 unaffected controls (InPSYght study). Data from the WGSPD grant were incorporated into SCHEMA meta-analysis.
**Acknowledgements:** The authors would like to acknowledge and thank the study participants and their families. The WGSPD is a public-private partnership between the National Institute of Mental Health (NIMH), the Stanley Center for Psychiatric Research, and researchers at 11 academic institutions across the USA. This work was supported by grants from the National Institute of Mental Health (NIMH): U01 MH105653 (M.B.), U01 MH105641 (S.A.M.), U01 MH105573 (C.N.P.), U01 MH105670 (D.B.G.), U01 MH105575 (M.W.S., A.J.W.), U01 MH105669 (M.J.D., K.E.), U01 MH105575 (N.B.F., D.H.G., R.A.O.), U01 MH105666 (A.P.), U01 MH105630 (D.C.G.), U01 MH105632 (J.B.), U01 MH105634 (R.E.G), U01 MH100239-03S1 (M.W.S., S.J.S., A.J.W.), R01 MH095454 (N.B.F.), the Simons Foundation: (SFARI #385110, M.W.S., S.J.S., A.J.W., D.B.G., SFARI #401457 (D.H.G)), and a gift from the Stanley Foundation (S.E.H.).

Taiwanese Trios
**Consortium/Collaboration:** SCHEMA 1 Collections
**Investigator:** Ming

Swedish Bipolar Collection (SWEBIC)
**Consortium/Collaboration:** SCHEMA 1 Collections
**Investigator:** Mikael Landén
**Country:** Sweden
**Sample size:** 6 cases, 637 controls
**Description:** The Swedish Bipolar Collection (SWEBIC) is a large-scale research study and biobank designed to investigate the genetic, environmental, and clinical factors contributing to bipolar disorder. Covering over 11,000 patients, SWEBIC has collected DNA and clinical data (often via the BipoläR registry) to identify risk factors, biomarkers for suicide attempts, and treatment responses

Avon Longitudinal Study of Parents and Children (ALSPAC)
**Consortium/Collaboration:** Control Collections
**Investigator:** Nicholas Timpson
**Country:** United Kingdom
**Sample size:** 0 cases, 2841 controls
**Description:** ALSPAC is a large geographically homogeneous prospective birth cohort from the southwest of England established to investigate environmental and genetic characteristics that influence health, development and growth of children and their parents.8-11 ALSPAC is now a three-generational study, comprising ‘G0’: the cohort of original pregnant women, the biological father and other carers/partners; ‘G1’: the cohort of index children and ‘G2’: the cohort of offspring of the index children.Full details of the cohort and study 51 design have been described previously and are available at http://www.alspac.bris.ac.uk. Please note that the study website contains details of all the data that are available through a fully searchable data dictionary and variable search tool (http://www.bristol.ac.uk/alspac/researchers/our-data/). Samples were selected for whole exome sequencing at the

Broad Institute from the G1 cohort (the cohort of index children) and were from subjects who were singletons/unrelated and of European/British ancestry, had blood-derived DNA available, and had been genotyped on a whole genome genotyping array. Subjects were also chosen based on the availability of phenotype data. Ethical approval for the study was obtained from the ALSPAC Ethics and Law Committee and the Local Research Ethics Committees. Consent for biological samples was collected in accordance with the Human Tissue Act (2004) and informed consent for the use of data collected via questionnaires and clinics was obtained from participants following recommendations of the ALSPAC Ethics and Law Committee at the time. Written informed consent was obtained from mothers at recruitment, from the main carers (usually the mothers) for assessments on the children from ages 7 to 16 years and, from age 16 years onwards, the children gave written informed consent at all assessments. Data is available via an application through the ALSPAC portal: http://www.bristol.ac.uk/alspac/researchers/access/.
**Ethics/IRB approval:** The UK Medical Research Council and Wellcome (Grant ref: 217065/Z/19/Z) and the University of Bristol provide core support for ALSPAC. This publication is the work of the authors who will serve as guarantors for the contents of this paper. We are extremely grateful to all the families who took part in this study, the midwives for their help in recruiting them, and the whole ALSPAC team, which includes interviewers, computer and laboratory technicians, clerical workers, research scientists, volunteers, managers, receptionists and nurses. The ALSPAC samples were sequenced with funding from the Stanley Center for Psychiatric Research, Broad Institute of MIT and Harvard

Dutch Controls
**Consortium/Collaboration:** Control Collections
**Investigator:** Danielle Posthuma
**Country:** Netherlands
**Sample size:** 0 cases, 789 controls
**Acknowledgements:** Dutch controls: Sequencing of the Dutch controls was funded by the Dalio Foundation

German Controls
**Consortium/Collaboration:** Control Collections
**Investigator:** Andreas Reif
**Country:** Germany
**Sample size:** 0 cases, 401 controls

Control Collection - NIDDK (IBD)
**Consortium/Collaboration:** Control Collections
**Investigator:** Mark Daly
**Country:** USA
**Sample size:** 0 cases, 8415 controls
**Acknowledgements:** NIDDK IBDGC: We thank the National Institute of Diabetes and Digestive and Kidney Diseases (NIDDK) IBD Genetics Consortium (IBDGC) supported by The Helmsley Charitable Trust and the Centers for Common Disease Genomics (NHGRI CCDG). Whole exome sequencing was done by the Broad Genomics Platform and supported by the NHGRI CCDG grant (UM1HG008895).
**Data availability:** phs001642.v2.p1

Control Collection - Type 2 Diabetes Knowledge Portal
**Consortium/Collaboration:** Control Collections
**Investigator:** Jose Florez
**Country:** USA
**Sample size:** 0 cases, 834 controls

USA Controls 1
**Consortium/Collaboration:** Control Collections
**Investigator:** Robert Yolken
**Country:** USA
**Sample size:** 0 cases, 112 controls
**Description:** Control samples were collected as part of a larger study about infectious agents and immune factors in serious mental illness. Psychiatric participants were recruited at a large psychiatric health system and nonpsychiatric

controls from the same geographic region. The diagnosis of non-psychiatric participants was confirmed with a structured clinical interview13,14 based on DSM-IV (American Psychiatric Association). All participants provided written informed consent. The study was approved by the IRB of the institution where the study was performed and included a data sharing agreement. Whole exome sequencing was done by the Broad Genomics Platform. Data is hosted on the Terra platform (http://app.terra.bio).
**Acknowledgements:** USA Controls 1: Sequencing of the USA controls was funded by the Dalio Foundation.

USA Controls 2
**Consortium/Collaboration:** Control Collections
**Investigator:** Jordan Smoller
**Country:** USA
**Sample size:** 0 cases, 3092 controls
**Description:** Samples were collected as part of the International Cohort Collection for Bipolar Disorder (ICCBD). 15,16 The Massachusetts General Hospital site of the ICCBD collected DNA from cases (patients with bipolar disorder) and controls by linking discarded blood samples to de-identified electronic health record (EHR) data. Cases and controls were identified by deriving EHR-based phenotyping algorithms applied to the Partners Healthcare Research Patient Data Registry (RPDR), described in detail previously.17 Full details of the algorithm are provided in 16. Samples were whole exome sequenced by the Broad Genomics Platform 52 and data from control samples were included in Epi25 analyses. Data is hosted on the Terra platform (http://app.terra.bio).
**Acknowledgements:** USA Controls 2: ICCBD sample and data collection was supported by R01 MH085542. Sequencing of the USA controls was funded by the Dalio Foundation.

Control Collections
**Investigator:** Rolf Adolfsson
**Country:** Sweden
**Sample size:** 0 cases, 376 controls
**Description:** Cases of European ancestry were ascertained from multiple different studies of schizophrenia (1992-2009). The diagnostic processes were similar between studies, and the final diagnosis is a best-estimate consensus lifetime diagnosis based on multiple sources of information such as clinical evaluation by research psychiatrists, different types of semi-structured interviews made by trained research nurses and research psychiatrists, medical records, course of the disease and data from multiple informants. Diagnosis was made in accordance with the Diagnostic and Statistical Manual of Mental Disorders-Version IV (DSM-IV)51 or International Classification of Diseases, 10th Revision (ICD-10)52 criteria. Controls were recruited from the Betula study, an ongoing longitudinal, prospective, population-based study from the same geographic area (North Sweden) that is studying aging, health, and cognition in adults53. All subjects (cases and controls) participated after giving written informed consent and the regional Ethical Review Board at the University of Umeå approved all original studies and participation in the PGC.

Autism Sequencing Consortium
**Consortium/Collaboration:** Control Collections
**Investigator:** Joseph Buxbaum
**Country:** United States
**Sample size:** 0 cases, 121 controls
**Description:** Founded in 2010, the Autism Sequencing Consortium (ASC) is an international group of scientists who share autism spectrum disorder (ASD) samples and genetic data.

Brazil Controls
**Consortium/Collaboration:** Control Collections
**Investigator:** Kathleen Barnes
**Country:** Brazil
**Sample size:** 0 cases, 400 controls

Colombia Controls
**Consortium/Collaboration:** Control Collections
**Investigator:** Kathleen Barnes
**Country:** Colombia

**Sample size:** 0 cases, 403 controls

Finnish Collection - AD, FINRISK, HEALTH2000, IBD
**Consortium/Collaboration:** Control Collections
**Investigator:** Aarno Palotie
**Sample size:** 0 cases, 5671 controls
**Description:** Collections include Finnish IBD and AD collections and population cohorts Health 2000 and FINRISK. FINRISK – The population-based FINRISK study has been followed up for IBD and other disease end-points using annual record linkage with the Finnish National Hospital Discharge Register, the National Causes-of-Death Register and the National Drug Reimbursement Register. Controls were chosen to have a high polygenic risk score for IBD without an IBD diagnosis. A detailed description of the FINRISK cohort can be found at Borodulin et al. The FINRISK controls were part of the FINRISK studies supported by THL (formerly KTL: National Public Health Institute) through budgetary funds from the government, with additional funding from institutions such as the Academy of Finland, the European Union, ministries and national and international foundations and societies to support specific research purposes. BB2017_2_Implementation_Agreement. Health 2000 and 2011 Surveys: Health 2000 Survey, a comprehensive combination of health interview and health examination survey, was carried out in years 2000-2001. The study captured a nationally representative sample of more than 8000 donors aged 30 and over living in the mainland Finland. The main aim of the Health 2000 Survey was to obtain information on the most important public health problems present in working-aged and aged population,  as well as investigate the population's functional and working capacity. More than 1000 participants of the Health 2000 Survey had previously also participated in the Mini-Finland Health Examination Survey twenty years earlier. A follow-up study of the Health 2000 Survey was conducted in 2011, in which extensive data was collected  from more than 5000 individuals using physiological measurements, interviews, and questionnaires and new samples were collected. The cohort is continuously followed-up by linkage to national health registers.
**Ethics/IRB approval:** BB2017_2_Implementation_Agreement, BB2017_12_Implementation_Agreement, AD 28/2007, Kuopio University Hospital for IBD 25/1995 Helsinki University Hospital
**Data availability:** DUOS

Control Collection - MIGEN (Ottawa)
**Consortium/Collaboration:** Control Collections
**Sample size:** 0 cases, 1717 controls
**Description:** MIGEN-Ottawa Controls – The Ottawa Heart Study is a cross-sectional case-control study designed to identify genes that predispose to angiographically defined coronary artery disease. All exome sequencing was performed at the Broad Institute of Harvard and MIT; sample sequence capture was performed using Agilent SureSelect Human All Exon Kit v2 and sequencing was performed on an Illumina HiSeq 2000 or 2500. Control data from this study was utilized.
**Data availability:** phs000294.v1.p1

Control Collection - MIGEN (Leicester)
**Consortium/Collaboration:** Control Collections
**Sample size:** 0 cases, 2248 controls
**Description:** MIGEN-Leicester Controls – cases were ascertained from two studies: (i) the British Heart Foundation Family Heart Study and (ii) the BRICCS Study. Control subjects were ascertained from the control subjects being recruited as part of the UK Aneurysm Growth Study (UKAGS). All exome sequencing was performed at the Broad Institute of Harvard and MIT; sample sequence capture was performed using Illumina's ICE Capture reagent and sequencing was performed on an Illumina HiSeq 2000 or 2500. Control data from this study was utilized.
**Data availability:** phs000294.v1.p1

Control Collection - 1000 Genomes
**Consortium/Collaboration:** Control Collections
**Country:** dbGAP
**Sample size:** 0 cases, 58 controls

Control Collection - dbGAP collections
**Consortium/Collaboration:** Control Collections
**Sample size:** 0 cases, 488

19. The “All of Us” Research Program | New England Journal of Medicine. https://www.nejm.org/doi/full/10.1056/NEJMsr1809937.
